# Directed Evolution of Adenine Base Editors with Increased Activity and Therapeutic Application

**DOI:** 10.1101/2020.03.13.990630

**Authors:** Nicole M. Gaudelli, Dieter K. Lam, Holly A. Rees, Noris M. Solá-Esteves, Luis A Barrera, David A. Born, Aaron Edwards, Jason M. Gehrke, Seung-Joo Lee, Alexander J Liquori, Ryan Murray, Michael S. Packer, Conrad Rinaldi, Ian M Slaymaker, Jonathan Yen, Lauren E. Young, Giuseppe Ciaramella

**Author notes:** Department of Hematology, St. Jude’s Hospital, Memphis, TN USA.

## Abstract

The foundational adenine base editors (*e.g.* ABE7.10) enable programmable C•G to T•A point mutations but editing efficiencies can be low at challenging loci in primary human cells. Here we further evolve ABE7.10 using a library of adenosine deaminase variants to create ABE8s. At NGG PAM sites, ABE8s result in ∼1.5x higher editing at protospacer positions A5-A7 and ∼3.2x higher editing at positions A3-A4 and A8-A10 compared with ABE7.10. Non-NGG PAM variants have a ∼4.2-fold overall higher on-target editing efficiency than ABE7.10. In human CD34+ cells, ABE8 can recreate a natural allele at the promoter of the γ-globin genes HBG1 and HBG2, with up to 60% efficiency, causing persistence of fetal hemoglobin. In primary human T cells, ABE8s achieve 98-99% target modification which is maintained when multiplexed across three loci. Delivered as mRNA, ABE8s induce no significant levels of sgRNA-independent off-target adenine deamination in genomic DNA and very low levels of adenine deamination in cellular mRNA.

---

Adenine base editors (ABE) allow the efficient programmable conversion of adenine to guanine in target DNA without creating double strand breaks (DSBs)^1-3^. ABE is a molecular machine comprising an evolved *E. coli* tRNA^ARG^ modifying enzyme, TadA, covalently fused to a catalytically impaired Cas9 protein (D10A nickase Cas9, nCas9) (Fig. 1a and 1b). A single guide RNA (sgRNA) directs ABE to a target genomic DNA sequence and, upon binding and stable R-loop formation, a short stretch of single-stranded nucleotides becomes accessible to TadA, an enzyme that chemically converts adenine to inosine. Inosine exclusively base-pairs with cytosine in DNA polymerase binding pockets^4^, resulting in an ABE-catalyzed A•T to G•C transition mutation at user-defined base-pairs following DNA replication or strand resection and nick repair (Fig. 1a).

**Fig. 1.**
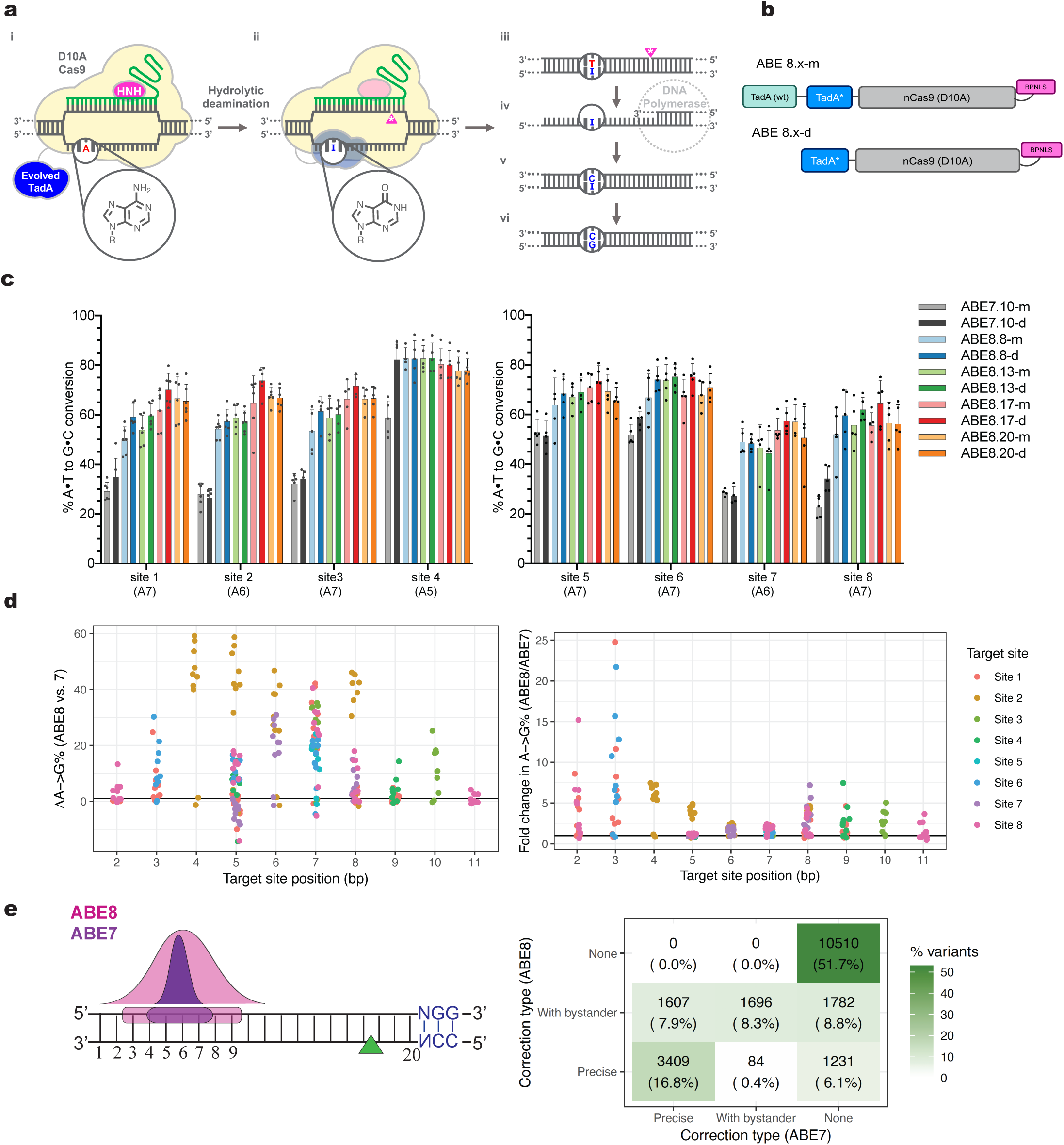
Eighth generation adenine base editors mediate superior A•T to G•C conversion in human cells. **a**, Adenine base editing overview. (i) ABE8 creates an R-loop at a sgRNA-targeted site in the genome. (ii) TadA* deaminase chemically converts adenine to inosine via hydrolytic deamination on the ss-DNA portion of the R-loop. (iii) nCas9 (D10A) nicks the strand opposite of the inosine containing strand. (iv) the inosine containing strand can be used as a template during DNA replication (v) inosine preferentially base pairs with cytosine in the context of DNA polymerases (ref). (vi) following replication, inosine may be replaced by guanosine. **b**, architecture of ABE8.x-m and ABE8.x-d. The nomenclature for ABE8.x-m/d used in this work is as follows: ABE8s are adenine base editors developed in this work resultant from an additional round of evolution (round 8) proceeding ABE7.10 evolution campaign (7 iterations of evolution conducted^1^. The “x” numerical value of ABE8.x indicate which mutations are included in the evolved TadA protomer of the corresponding ABE8 editor; each number represents a different set of mutations described in Supplementary Fig. 1. The indication of “m” or “d” denotes whether the ABE8 construct contains either an N-terminal wild-type TadA linked to the evolved TadA (“d”) or contains the TadA evolved variant only (“m”). **c**, A•T to G•C base editing efficiencies of core ABE8 constructs relative to ABE7.10 constructs in Hek293T cells across eight genomic sites. Values and error bars reflect the mean and s.d. of five independent biological replicates performed on different days. **d**, Absolute and fold changes in base editing between ABE8 and ABE7. Representation of average ABE8:ABE7 A•T to G•C editing in Hek293T cells across all As within the target of eight different genomic sites. Positions 2-12 denote location of a target adenine within the 20-nt protospacer with position 20 directly 5’ of the -NGG PAM. Each point is shown as a comparison to the median value for ABE7.10 editors at the same site and position. (left) The absolute difference (ABE8-ABE7) in editing at each position is shown (right) The ratio of ABE8:ABE7 editing is shown. Colors indicate the target site in which the observation was made. **e**, (left) ABE7.10 and ABE8 editor activity windows are shown. Numbers indicate the position within the protospacer. The location of an induced nick in the target DNA backbone is indicated by a triangle and corresponding PAM recognition sequence is shown. (right) Comparison of targetable sites, ABE7 vs. ABE8. Each box shows the number and percentage of pathogenic G->A or C->T SNV variants in the ClinVar database^48,49^ that can be targeted with ABE7 or ABE8. The analysis considers a 20-nt protospacer sequence and a Cas9 that can target NGG or NGA PAMs. The editing windows were assumed to be 5-7 for ABE7 and 4-8 for ABE8. Precise correction implies that only the pathogenic mutation is editable within the specified window in at least one possible spacer/PAM combination. If all possible correction strategies involve other modified bases modified, the corresponding variants are counted in the “with bystander” category.

To date, seventh generation ABEs (ABE7) have enabled efficient A•T to G•C conversion in the genomes of humans^5^, mice^6-8^, bacteria^1^, plants^9,10^, and a variety of other species, reviewed here^11^. Many therapeutic targets, however, may benefit from a more active ABE with a broader editing window or improved compatibility with non-NGG nCas9s^12-14^ as well as increased editing efficiencies in human cell lines^15^ or when used *in vivo*^8^. The need for a more active version of ABE7.10 is the greatest when target adenines are positioned on the outer edges of the canonical ABE editing window (positions 3, 4, 7 and 8).

## Results

### Evolution of ABE7.10 to create ABE8 constructs with higher activity

To diversify and expand our ABE toolbox, we further evolved the DNA-modifying TadA enzyme contained in ABE7.10 (TadA*7.10) for higher activity. Building on the previously developed bacterial selection strategy^1^, we increased the stringency of the selection system by requiring ABE to induce three concurrent A•T to G•C reversion edits to survive antibiotic selection (Supplementary Sequence 1). As an additional refinement to the protocol, we utilized a synthetic library of TadA sequences containing all 20 canonical amino acid substitutions at each position of TadA, with an average frequency of 1-2 nucleotide substitution mutations per library member. This chemically-synthesized library enables access to a greater sequence space than is achievable with error-prone PCR techniques used in previous studies^1^.

In the course of the ABE8 evolutionary campaign, we identified eight mutations within TadA* that were enriched with high frequency from ∼300 isolated clones (Supplementary Fig. 1). Six of the eight identified amino acid mutations required at least two nucleobase changes per codon; codons with two nucleobase changes were unobserved with the previously published TadA error-prone PCR libraries^1^. Two of the enriched mutations alter residues proximal to the active site of adenine deamination (I76 and V82) (Supplementary Fig. 2). In addition to the four previously reported mutations in the C-terminal alpha helix of TadA*7.10, we observed two new additional mutations within the same alpha-helix (Y147R and Q154R) (Supplementary Fig. 2). We confirmed this highly mutated alpha-helix is indeed necessary for robust product formation by demonstrating that, upon truncation, base editing efficiency was substantially reduced (Supplementary Fig. 3).

### Characterization of evolved ABE8s in mammalian cells

To test the activity of TadA* variants in mammalian cells, we utilized ABE codon optimization and NLS orientation with the most favorable on- and off-target profile (see Supplementary Note 1 and Supplementary Fig. 4). The eight enriched TadA* mutations were incorporated into ABE7.10 in various combinations, yielding forty new ABE8 variants (Supplementary Fig. 1). We also made two architectural variants of ABE8 where the TadA region of ABE is either a heterodimeric fusion of a wild-type (TadA) and evolved (TadA*) protomer or a single protomer of an engineered TadA, resulting in a ∼500 base-pair smaller editor. These architectural variants are referred to as ABE8.x-d and ABE8.x-m respectively (Fig. 1b).

First, these forty constructs were evaluated for their on-target DNA editing efficiencies relative to ABE7.10 across eight genomic sites containing target A bases in positions ranging from 2 to 20 (where NGG PAM = positions 21, 22, 23) within the canonical 20-nt *S. pyogenes* protospacer (Supplementary Fig. 5). In agreement with reports for ABE7.10^16-18^, we found that the N-terminal wild-type TadA construct was not necessary for robust DNA editing using ABE8. Indeed, constructs containing the N-terminal, wild-type TadA (ABE8.x-d) perform similarly in terms of editing window preference, total DNA editing outcome, and indel frequency relative to its economized architecture (ABE8.x-m) (Fig. 1c and Supplementary Fig. 5 and 6). Although intra-construct, TadA(wt):TadA*8 dimerization may not be necessary for ABE8 activity, these studies do not preclude the possibility of *in trans* TadA*8:TadA*8 dimerization occurring between ABE8 expressed base editors as has been observed in plant nuclei^18^.

Across all sites tested, ABE8s result in ∼1.5x higher editing at canonical positions (A5-A7) in the protospacer and ∼3.2x higher editing at non-canonical positions (A3-A4, A8-A10) compared with ABE7.10 (Fig. 1d). The sequence of the target, the position of the A within the target window and the sequence identity of the ABE8 construct itself are all factors that can impact the editing efficiency (Fig. 1c, 1d, Supplementary Fig. 5). Overall, the median change in editing across all positions, in all sites tested is 1.94-fold relative to ABE7.10 (range 1.34 - 4.49). The increased activity of ABE8s across the editing window enables reversion of an additional ∼3000 more disease-associated mutations identified in the ClinVar database (Fig. 1e). Whilst ABE8 broadens the therapeutic scope of adenine base editors, we computationally identified cases where ABE8 may create a bystander edit that could be avoided with ABE7.10, highlighting the need to select the appropriate adenine base editor tool depending on the target sequence and the desired outcome.

Next, from the large ABE8 pool of forty constructs, we selected a sub-set of ABE8 constructs (ABE8.8-m, ABE8.13-m, ABE8.17-m, ABE8.20-m, ABE8.8-d, ABE8.13-m, ABE8.17-d and ABE8.20-d) to evaluate in greater detail. These constructs represent ABE8s with distinct differences in editing performance amongst the 8 genomic sites as determined through a hierarchical clustering analysis (Supplementary Fig. 7). These ABE8s all significantly outperform ABE7.10 at all genomic sites tested (P-value = 0.0006871, two-tailed Wilcoxon rank sum test) and encompass a variety of combinations of mutations identified from the ABE8 directed evolution campaign (Supplementary Fig. 8 and 9).

### ABE8s with either non-NGG PAM nCas9 variants or catalytically dead *S. py.* Cas9

Although ABE variants recognizing non-NGG PAMs have been described ^12-14^, editing efficiencies of these constructs are decreased in many instances when compared to outcomes observed with *S. pyogenes* Cas9 targeting NGG PAM sequences^12-14^. To determine whether our eighth-generation evolved deaminase also increases the editing efficiencies at target sites bearing non-NGG PAMs, we created ABE8 editors that replace *S. pyogenes* Cas9 with an engineered *S. py.* variant, NG-Cas9 (PAM: NG)^19^ or *Staphylococcus aureus* Cas9 (SaCas9, PAM: NNGRRT)^*20*^. Encouragingly, we observed median increases in A•T to G•C editing frequencies of 1.6 and 2.0 fold, respectively, when comparing ABE8 variants to ABE7.10 for both SpCas9-NG (NG-ABE8.x-m/d) and SaCas9 (Sa-ABE8.x-m/d) (Fig. 2a and b and Supplementary Figs. 10-13). Similar to SpCas9-ABE8, the greatest differences in editing efficiencies between ABE7.10 and ABE8 constructs for the non-NGG PAM variants are observed at target As located at the periphery of the preferred position in the editing window (*S. pyogenes*: positions 4-8; *S. aureus*: positions 6-13; summarized here^3^). ABE8 orthologs utilizing non-NGG PAMs broaden the targeting scope for efficient A base editing.

**Fig. 2.**
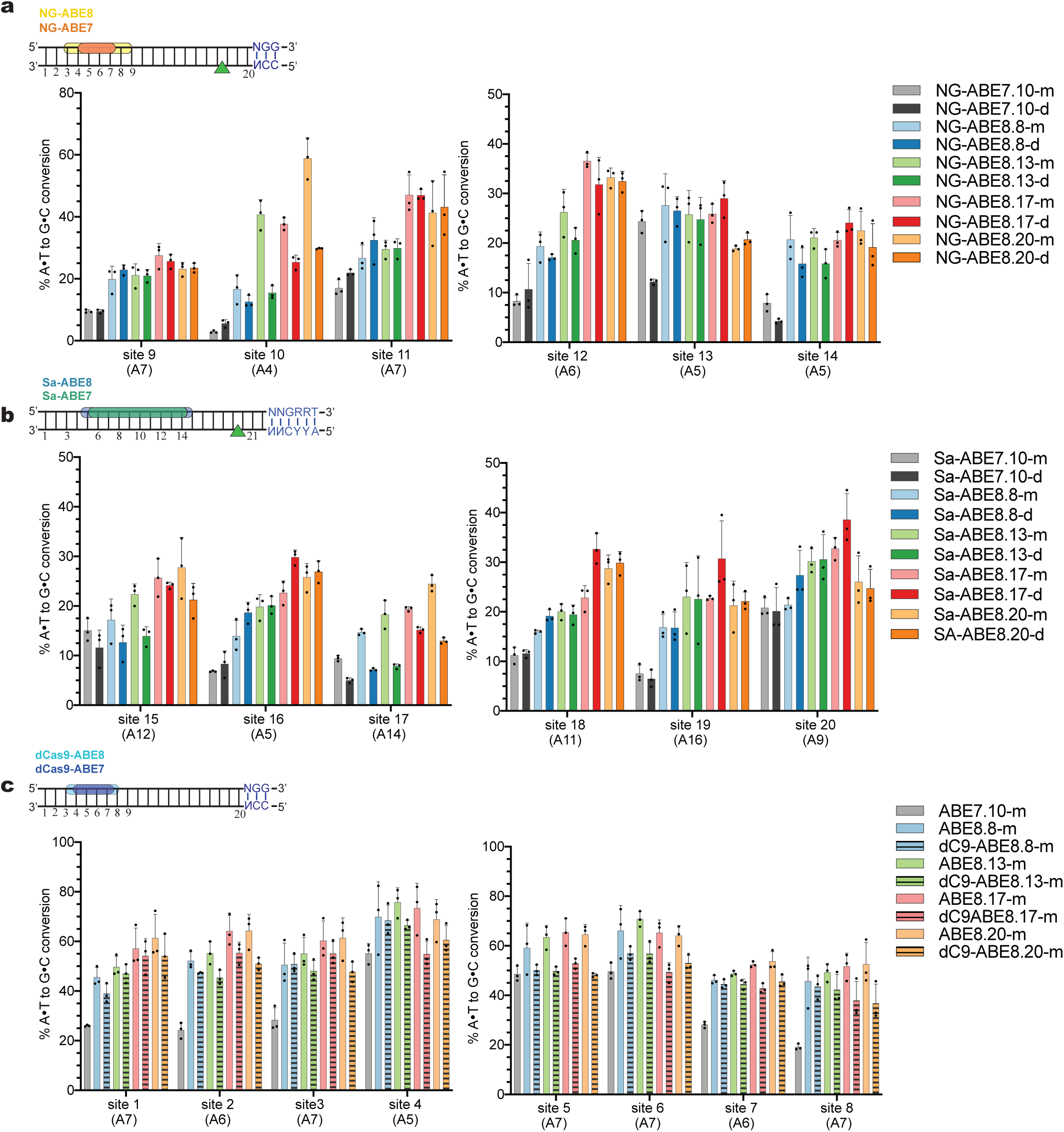
Cas9 PAM-variant ABE8s and catalytically dead Cas9 ABE8 variants mediate higher A•T to G•C conversion than corresponding ABE7.10 variants in human cells. **a-c**, A•T to G•C conversion in Hek293T cells with **a**, NG-Cas9 ABE8s (-NG PAM) **b**, Sa-Cas9 ABE8s (-NNGRRT PAM) and **c**, catalytically inactivated, dCas9-ABE8s (D10A, H840A in *S. pyogenes* Cas9). Values and error bars reflect the mean and s.d. of three independent biological replicates performed on different days. ABE7.10 and ABE8 editor activity windows are shown. Numbers indicate the position within the protospacer. The location of an induced nick in the target DNA backbone is indicated by a triangle and corresponding PAM recognition sequence is shown.

For applications where minimizing indel formation is necessary, we explored the effect of replacing the catalytically impaired D10A nickase mutant of Cas9, with a catalytically “dead” version of Cas9 (D10A + H840A)^21^ in the core eight ABE8 constructs (“dC9-ABE8.x-m/d”). By replacing the nickase with dead Cas9 in ABEs, we observed a >90% reduction in indel frequency to rates barely above background (0.3-0.8%) for dC9-ABE8 variants relative to ABE7.10 while maintaining a significantly higher (2.1-fold), on-target DNA editing efficiency (Fig. 2c, Supplementary Figs. 14-16). Encouragingly, dC9-ABE8 variants only have a median 14% reduction in on-target DNA editing efficiencies relative to nickase-active ABE8s.

Another class of undesired ABE-mediated genome edits at an on-target locus can be an ABE-dependent cytosine to uracil (C•G to T•A) conversions^16,22^. At the eight target sites tested, we measured the 95^th^ percentile of C-to-T editing to be 0.45% with ABE8 variants and 0.15% with ABE7.10-d or -m, indicating that on-target cytosine deamination with ABEs can occur but the frequencies are generally very low (Supplementary Fig. 17). Together, these data indicate that ABE8s retain high specificity for A-to-G conversion.

### Application of ABE8s to therapeutic targets in primary human cells

Next, we evaluated the ABE8 constructs in human hematopoietic stem cells (HSC). *Ex vivo* manipulation and/or editing of HSCs prior to administration to patients as a cell therapy is a promising approach for the treatment of hematological disorders. It has been previously demonstrated that ABEs can introduce a T•A to C•G substitution at the −198 position of the promoter region of HBG1/2^1^. This naturally occurring allele, referred to as the “British mutation” yields Hereditary Persistence of Fetal Hemoglobin (HPFH) resulting in moderately increased levels (3.5-10%)^23^ of *γ*-globin into adulthood, which can mitigate the defects in *β*-globin seen in sickle cell disease and *β*-thalassemia^24^. With the goal of reproducing the HPFH phenotype and evaluating the clinical relevance of ABE8, we isolated CD34+ hematopoietic stem cells from two donors and transfected them with mRNA encoding ABE8 and ABE7.10 editors and end-modified sgRNA placing the target A at position 7 within the protospacer.

We found that average ABE8 editing efficiencies at the −198 HBG1/2 promoter target site were 2-3x higher than either ABE7.10 construct at early time points (48h), and 1.3-2-fold higher than either ABE7.10 at the later time (144h) (Fig. 3a, Supplementary Fig. 18 and 19). Next, the amount of *γ*-globin protein produced following ABE treatment and erythrocyte differentiation was quantified by UPLC (Supplementary Tables 5-25). We observed a 3.5-fold average increase in % *γ*-globin/*α*-globin expression in erythrocytes derived from the ABE8 treatment groups when compared to mock treated cells and a statistically significant increase (10.0% absolute change, 1.3-fold relative change), when comparing the median *γ*-globin levels across all ABE8-treated vs. ABE7-treated samples (P-value = 0.004954, Wilcoxon-Mann-Whitney U test, two-tailed; Fig. 3b, Supplementary Fig. 20). Additionally, we investigated the genotype-phenotype correlation between the alleles generated by each editor and the corresponding *γ*-globin induction level. We identified a model that is highly correlated with our data (R^2^ = 0.84, Fig. 3c, Supplementary Note 2) and this model suggests both the 7G and 7G+8G alleles contribute to *γ*-globin induction. At least 20% of HbF-expressing cells is hypothesized to be required to ameliorate symptoms of sickle cell disease and likely slightly higher levels of HbF are needed for *β*-thalassemia.^25,26^ Encouragingly, the *γ*-globin levels observed following ABE8 editing are consistent with HbF levels higher than these thresholds and greater than levels achieved with ABE7.10.

**Fig. 3.**
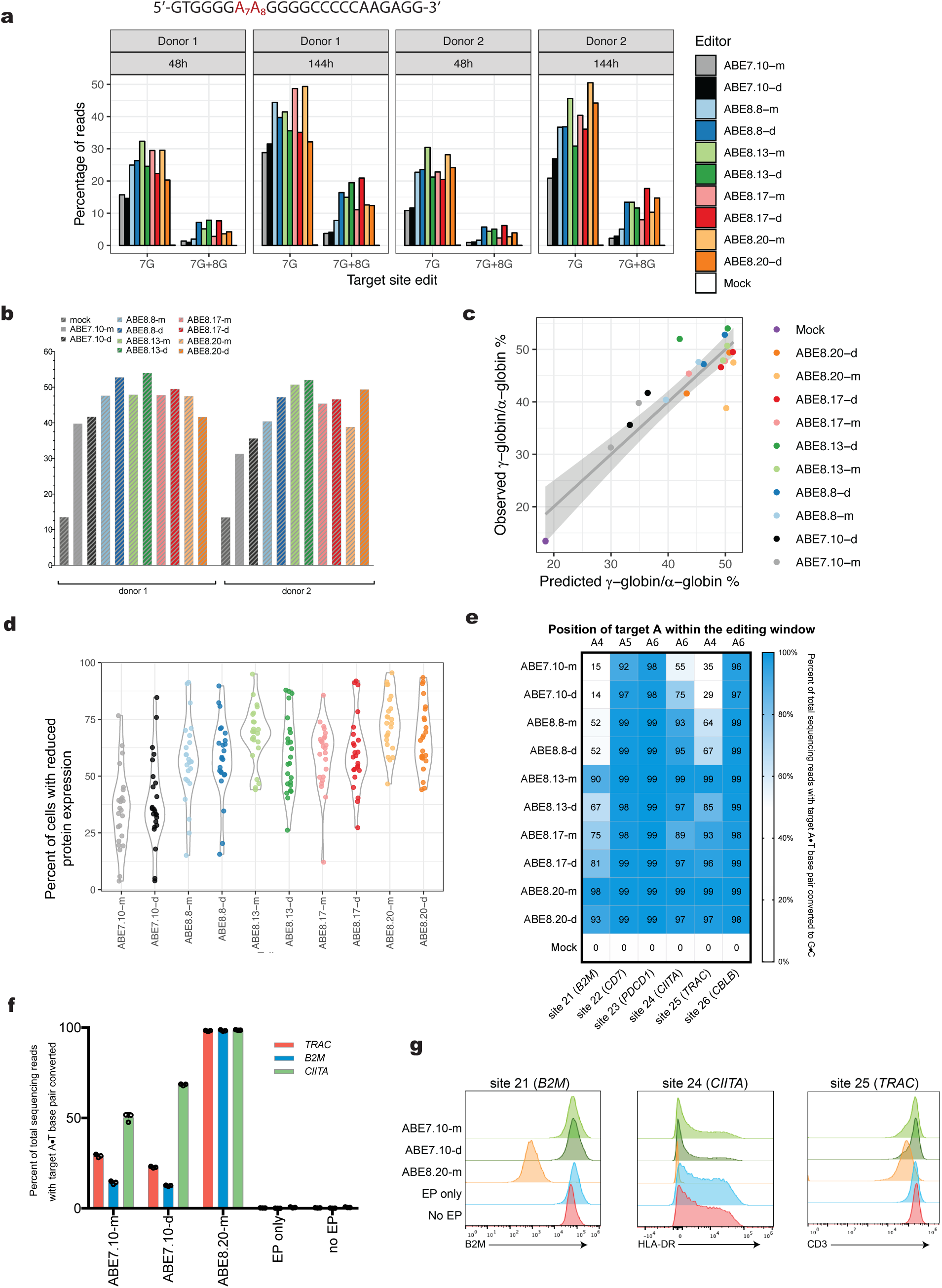
A•T to G•C conversion and phenotypic outcomes in primary human cells. **a**, graphical representation of distribution of total sequencing reads that contain either G_7_ or combined (G_7_ + G_8_) alleles. −198 target sequence denoted above; target adenines highlighted in red. **b**, percentage of *γ*-globin formed as a fraction of alpha-globin. Values shown from two different donors, post ABE treatment and erythroid differentiation. **c**, comparison of predicted vs. observed gamma-globin induction using a linear regression model that accounts for the contributions of the 7G and 7G + 8G alleles across samples. This model is described in more detail in Supplementary Note 2**. d**, violin plot of reduced protein expression as measured by flow cytometry after primary human T cells were electroporated with the indicated mRNA and 24 individual sgRNAs targeting six genes. Individual values shown represent the mean percent of cells with reduced protein expression from two replicates of cells edited with the indicated mRNA and one of the 24 sgRNAs tested. **e**, NGS analysis of A•T to G•C conversion at six target sites by eight ABE8 mRNAs and ABE7.10-m/d. Values shown reflect the mean of three independent biological replicates. The position of the edited nucleotide for each target site is shown above the heat map. **f**, NGS analysis of A•T to G•C conversion at site 21 (*B2M*), site 24 (*CIITA*), and site 25 (*TRAC*) after primary human T cells were electroporated with the indicated mRNA and three sgRNAs in multiplex editing format. **g**, protein expression of the three target genes as measured by flow cytometry on the cell populations in (e) five days post-electroporation. Data shown is a representative plot from two donors, repeated three times for each donor.

We next evaluated the activities of ABE8 in primary human T cells. Genetically modified T cells have demonstrated clinical efficacy in some therapeutic applications^27^, and there is an increasing body of evidence suggesting that the therapeutic potential of adoptive T cell therapies may be significantly enhanced by disruption of multiple genes in the same cell to achieve desirable cellular phenotypes.^28,29^ Approaches using nucleases to introduce indel mutations in target genes, thereby disrupting their expression in T cells^30,31^, are effective but simultaneous creation of multiple DSBs in a target cell can result in genomic rearrangements and toxicities with variable frequencies^32,33^. Because ABEs function by making single nucleotide genomic changes without creating DSBs, multiplexed base editing with ABE8 is an attractive approach for creating genetically modified T cells. To this end, we first asked whether ABE8 could be used to prevent the expression of single genes relevant to the creation of T cell therapies by targeting conserved sequence motifs at mRNA splice sites, a strategy successfully used with cytosine base editors^30^. We screened eight of the highest-performing ABE8s, in addition to ABE7.10, for activity by individually transfecting primary human T cells with mRNA encoding each editor and 24 sgRNAs targeting six total genes, and measured protein knockdown by flow cytometry as a proxy for genomic editing (Fig. 3d). Across all sgRNAs, ABE7.10s induced protein knockdown with between 2%-85% efficiency (median of 20.7% and 26.4% for ABE7.10-m and ABE7.10-d, respectively). Although all ABE8s outperformed their ABE7.10 counterparts, ABE8.20-m consistently produced the highest protein knockdown efficiencies (range of 4%-96%, median of 60%, Fig. 3d). We then measured the genomic editing efficiencies for each editor and the best performing target site for each gene (Fig. 3e, sites identified in Supplementary Fig. 21) using NGS. We found that ABE7.10s edited the six target sites with between 14-98% efficiency, while ABE8.20-m edited each of the same sites with between 98-99% efficiency, consistent with ABE8.20-m possessing improved editing capabilities. At most T cell target sites tested, ABE8 editors increased conversion of a single base relative to ABE7.10. In the case of the *CBLB* target (site 26), ABE8 editors created a different primary edit (4G+6G as opposed to 6G only, Supplementary Fig. 22) compared with ABE7.10. It should be appreciated that because the intention of targeting *CBLB* is to disrupt the target splice site, and thereby abrogate expression of the gene, the creation of a bystander edit at position 4 is inconsequential to the phenotypic outcome of the edit.

To determine whether ABE8 is capable of efficient multiplexed editing, we next sought to edit three genes simultaneously in primary human T cells. We targeted *B2M, CIITA*, and *TRAC*, three genes that when disrupted confer reduced cell surface expression of MHC class I, MHC class II, and the T cell receptor^30,34,35^ respectively, phenotypes that are hypothesized to reduce alloreactivity and immune recognition in the context of allogeneic cell therapies. ABE8.20-m edited each individual target with 98.1%, 98.3%, or 98.6% efficiency, improvements of 3.4, 6.9, and 1.4-fold over the best performing ABE7.10 (Fig. 3f). DNA editing efficiency correlated with reduced cell surface expression of B2M, HLA-DR, and CD3 (Fig. 3g). However, >98% genomic editing of the *TRAC* locus by ABE8.20-m resulted in only moderately reduced trafficking of the T cell receptor to the cell surface, indicating that modification of splice sites by ABE8 does not always fully abrogate mRNA splicing, and that protein expression must also be stringently evaluated for each sgRNA. Taken together, ABE8.20-m demonstrates the potential for adenine base editing to create T cell therapies featuring multiple highly efficient genetic edits which can confer a range of desirable therapeutic attributes.

### Off target analyses of ABE8s in mammalian cells

As with all base editors, ABE8s have the potential to act at off-target loci in the genome and transcriptome^1,2,16,17,36-41^. Guided by prior publications on this topic, we undertook an extensive assessment of the off-target deamination effects of ABE8s. To assess guide RNA-dependent DNA off-target base editing, we sequenced twelve off-target loci published to be cleaved by Cas9 when paired with three sgRNAs^42^. We confirmed through sequencing that when treated with Cas9 plasmid and the appropriate sgRNA, all of these twelve sgRNA-guided off-target loci yield detectable indel frequencies in our HEK293T cells (Supplementary Fig. 23). We also assessed unguided cellular mRNA deamination by base editors by targeted sequencing of 125-nt regions of two cellular mRNAs^17^.

We began by assessing whether previously-published mutations^16,17,40,41^, designed to mitigate spurious cellular RNA deamination for ABE7.10 is also compatible with our ABE8s (we chose ABE8.17-m to probe this question). All of the installed RNA off-target minimizing mutations decreased the on-target editing frequencies of ABE8.17-m to differing extents, with V106W^17^ and F148A^41^ impairing ABE8 the least (Supplementary Fig. 24a and 24b). Of these, only V106W was able to substantially reduce the level of off-target RNA and sgRNA-guided DNA editing (Supplementary Fig. 24c, d, and e). Accordingly, we proceeded with this construct since these data indicate that inclusion of V106W can enable high on-target editing efficiency with reduced DNA- and RNA-off-target deamination events.

To comprehensively assess the off-target editing associated with ABE8s, we compared the on- and off-target editing profiles of ABE7 and ABE8s when delivered as plasmids or as mRNA constructs (Supplementary Fig. 25). We note that although plasmid delivery of genome editing agents is widely used for base editing research purposes and represents a worst-case scenario for off-target editing effects, mRNA delivery is more effective for editing primary human cells^43^, and the delivery method for a base editor is a critical determinant of its off-target profile^36^. Encouragingly, we observe that the on-target efficiency of both ABE7 and ABE8 base editors is comparable between plasmid and mRNA delivery of ABE, despite the more transient lifetime of mRNA versus plasmid over-expression (Supplementary Fig. 25a, b).

When plasmid overexpression is used as a delivery modality for base editors, ABE8 constructs exhibit a 3- to 6-fold greater sgRNA-dependent DNA off-target editing frequencies than ABE7.10 (Supplementary Fig. 25c). Remarkably, when ABEs are delivered using mRNA, the guide-dependent DNA off-target editing associated with ABE8 constructs is decreased by an average of 1.5-fold (for ABE7.10-d) and 2.2-fold (for ABE8.20-m, the most heavily modified ABE8 relative to ABE7.10-d). Notably, off-target base editing activity associated with non-repetitive sgRNAs HEK2 and HEK3 was reduced from above 14% with plasmid delivery to below 0.4% by mRNA delivery (Supplementary Fig 25c,d). Thus, for therapeutic and other applications requiring high DNA editing specificity, we emphasize the benefits of mRNA delivery, careful choice of sgRNA^42,44^, and consideration of V106W inclusion in TadA to substantially reduce DNA off-target editing frequencies, in this case to that of the negative control for the non-repetitive sgRNAs HEK2 and HEK3 (Supplementary Fig 25d).

Next, we measured the levels of spurious deamination of cellular mRNA^16,17,40,41^ associated with ABE8s. To generate data on all the constructs of interest in a high-throughput manner, we began with targeted amplification and high throughput sequencing of two cellular RNAs in HEK293T cells treated with ABEs. Consistent with previous publications^16,17,40,41^ treatment with ABE7, and, to a greater extent, ABE8 lead to detectable A-to-I deamination in cellular RNA when delivered by plasmid overexpression (Supplementary Fig 24e). However, when delivered as an mRNA construct, the level of mRNA deamination was reduced by an average of 34-fold (in the case of ABE7.10-d) to 134-fold (ABE8.17-m) (Supplementary Fig. 25e and f), indicating that mRNA delivery can effectively reduce the frequency of cellular RNA editing.

To interrogate spurious cellular RNA deamination more thoroughly, we performed whole transcriptome sequencing of both HEK293T and human T cells treated with ABE7.10-d, ABE8.17-m, ABE8.20-m and ABE8.17-m+V106W-encoding mRNAs (Fig. 4a for T cells and Supplementary Fig. 26 for HEK293T cells). In both cell types, transcriptome-wide sequencing revealed a detectable increase in cellular adenine deamination in cells treated with ABE7.10-d, ABE8.17-m and ABE8.20-m relative to a Cas9 control (Fig.4a and Supplementary Fig. 26). However, we find that the elevated frequency of mRNA deamination is mitigated by inclusion of the V106W mutation in the ABE8.17m+V106W-treated samples (Fig. 4a for T cells and Supplementary Fig. 26 for HEK293T cells), indicating that careful choice of editor and delivery modality can mitigate and, in some cases, eliminate off-target cellular RNA deamination arising from ABE treatment for applications where transient RNA editing is of concern.

**Fig. 4.**
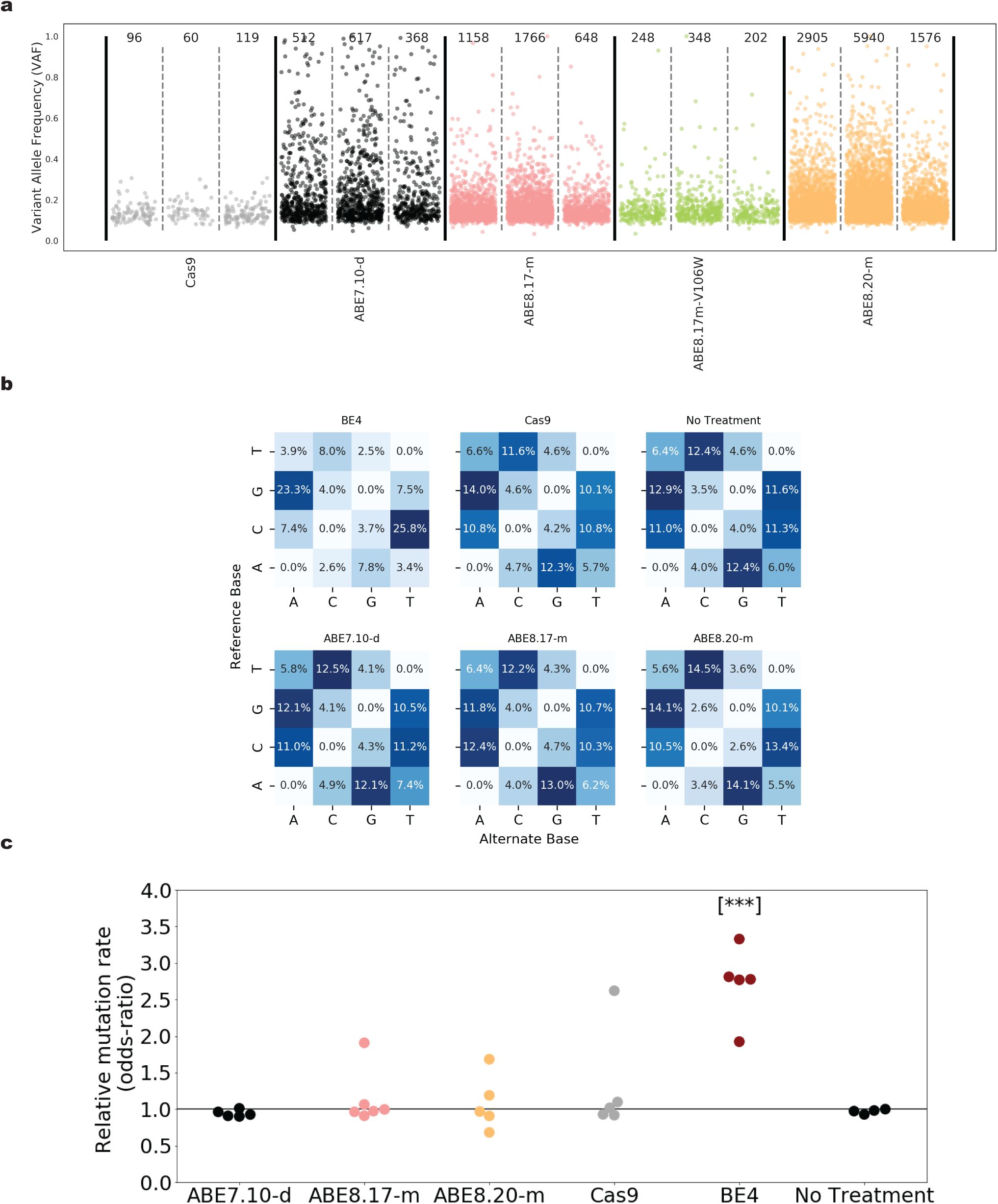
Whole transcriptome and whole genome sequencing data from cells treated with base editor mRNAs. **a**, Strip plot representing the variant allele frequency of transcriptome wide A-to-G mutations in RNA observed in three different T cell donors. Total A-to-G mutations indicated above each sample, which correspond to the sample size in each group. **b**, Mutational classification of all somatic mutations observed genome wide in single cell expanded cell populations. Each plot is from the sample from each treatment group with the median number of total mutations. All samples are shown in **c**, For each editor, the odds ratios quantify the fold change in mutation frequencies for the editor-induced mutation type (C-to-T for BE4 and A-to-G for others) and all other mutation types in each treatment replicate compared to an untreated control. [***] indicates a significant p-value (P = 0.010, one-sided Wilcoxon-Mann-Whitney U test) for a one-sided Mann-Whitney U test between treatment group and untreated control group. All five biological replicates for each condition tested are shown.

Finally, we performed whole genome sequencing (WGS) to assess the degree to which ABE8s may induce genome-wide point mutations in a sgRNA-independent manner (referred to here as ‘spurious deamination’) as has been previously reported for CBEs, but not ABE7.10 or Cas9^37,38,45,46^. Accordingly, we transfected HEK293T cells with base editor-encoding mRNA and sgRNA targeting *B2M* (Site 21) and, after seventy-two hours of incubation, we used fluorescence activated cell sorting (FACS) to isolate B2M-negative single cells that had been successfully base edited. Single-sorted cells were individually clonally expanded to generate sufficient genomic DNA to perform Illumina-based whole genome sequencing. Importantly, all treated samples were confirmed to be bi-allelically edited at the on-target *B2M* locus, indicating receipt of active base editor or Cas9. Consistent with previous results from ABE7.10-d-treated animals in embryo injection mouse experiments^37,38,45,46^, we found no detectable increase (P-value = 0.911, one-sided Wilcoxon-Mann-Whitney U test, Supplementary Table 26 and Supplementary Table 27) in genome-wide A-to-G mutations upon treatment with ABE7.10-d (Fig. 4b, 4c and Supplementary Fig. 27). Concurrently, we identified a statistically significant increase in C-to-T mutations in BE4-treated samples compared to untreated controls (P-value = 0.010, one-sided Wilcoxon-Mann-Whitney U test, Supplementary Table 26), consistent with previous reports^37,38,45,46^. We observed no statistically significant increase in A->G% mutation rates across the ABE8.17-m, ABE8.20-m or Cas9 groups when compared to untreated samples (P-value = 0.375, 0.643, and 0.27, respectively, one-sided Wilcoxon-Mann-Whitney U test, Fig. 4c and Supplementary Fig. 27, Supplementary Table 26). However, we note that individual samples in most treatment groups exhibited substantially higher or lower relative rates of A-to-G or C-to-T mutations, including a sample with elevated A-to-G mutation rates in the Cas9-treated group. We discuss potential sources of variability between clonally expanded cell populations in Supplementary Note 3. In summary, these results indicate that treatment with ABE7 or ABE8 does not lead to substantially elevated mutation rates as observed in this study and others for CBEs such as BE3 and BE4. These findings further characterize the DNA specificity of ABEs and are encouraging both for their use as research tools and in therapeutic applications.

## Discussion

Using directed evolution, we identified forty new ABE8s that have higher editing efficiencies on the ss-DNA R loop of Cas9 relative to ABE7.10 in mammalian cells. Creating these variants required both increasing the stringency of the bacterial selection and utilizing a fully chemically synthesized library of TadA variants in which all amino substitutions were represented within the library at each site across the open reading frame of the adenine deaminase.

We tested our forty ABE8 variants at eight sequence-diverse genetic loci in mammalian cells and were able to identify a core set of ABE8s with distinct editing properties, capturing the diverse editing outcomes across all forty ABE8 constructs generated.

The ABE8s evolved in this study are a powerful suite of new tools to add to the ABE editor toolbox, enabling robust, programmable A•T to G•C DNA editing at sites that were previously difficult to edit or at target As located beyond the reaches of ABE7.10’s editing window. In addition to the superior editing performance of canonical ABE8s, these molecular instruments have also been successfully implemented as non-NGG PAM variants, along with catalytically dead Cas9 variants, demonstrating their modularity and versatility. Analogous to editing outcomes with ABE7.10, ABE8s also maintain high product purity and generate low indel frequencies.

Additionally, we demonstrate that our ABE8s enable robust editing in both transformed and human primary cells when delivered as an mRNA construct, retaining their beneficial on-target properties relative to ABE7.10, as was observed with plasmid. In CD34+ cells, we successfully use mRNA encoding ABE8s to target and edit the HBG1/2 promoter region, resulting in higher upregulation of fetal gamma-globin compared to outcomes with ABE7.10. Furthermore, we show for the first time, highly efficient, multiplexed editing in T cells using ABE8, which can be used to confer beneficial properties in the context of cell therapies such as CAR-Ts.

Crucial toward realistic therapeutic use of a more potent ABE editor we evaluated the off-target effects of cells treated with ABE8s. We show here that mRNA delivery of our ABE8s result in minimal spurious editing of intracellular mRNA and that this transient editing can be mitigated with a V106W substitution in TadA*^17^. Most importantly, we experimentally demonstrate that treatment of mammalian cells with our ABE8s causes no significant genome-wide, guide-independent adenine deamination relative to untreated cells.

Taken all together, the technological advancement of ABE8s reported here represent a significant step forward in the use of adenine base editors as a viable therapeutic gene editing modality for correcting the most prevalent class of disease-associated SNPs in humans, editing regulatory elements to re-activate gene expression in CD34+ cells, and multiplexed editing for adoptive T cell therapies with desirable properties.

## Acknowledgements

Steve Cavnar is thanked for support with creating digital renderings of base editing mechanisms. Dana Levasseur and Bo Yan are acknowledged for technical advice and support with CD34+ experiments. Bob Gantzer, Jeremy Decker, and Matt Humes are thanked for NGS support. Scott Haskett is recognized and thanked for his FACS expertise and sorting mammalian cells used in WGS experiments.

## Author contributions

NMG conceived of and directed the work, conducted evolution and Hek293T experiments, performed analyses, and wrote the manuscript. DKL and NMS conducted mammalian cell transfections. HAR conducted all off-target studies and analyses except for whole transcriptome and whole genome sequencing analyses, that were conducted by LEY. MSP and JY designed CD34+ experiments and synthesized mRNA constructs. CR, JY, and AJL carried out all CD34+ experiments and associated UHPLC procedures. AE, RM, and JMG conducted all T-cell experiments and analyses. LAB conducted all statistical analyses of NGS data and generated figures. DAB created PyMol figures. SL and IMS designed truncation experiments. GC supervised the research. LEY processed, analyzed and generated figures for whole-transcriptome and WGS. NMG, HAR, MSP, JMG, LAB and GC all edited the manuscript.

## Methods

### General Methods

All cloning was conducted via USER enzyme (New England Biolabs) cloning methods^47^ and templates for PCR amplification were purchased as bacterial or mammalian codon optimized gene fragments (GeneArt). Vectors created were transformed into Mach T1^R^ Competent Cells (Thermo Fisher Scientific) and maintained at −80 C for long-term storage. All primers used in this work were purchased from Integrated DNA Technologies and PCRS were carried out using either Phusion U DNA Polymerase Green MultiPlex PCR Master Mix (ThermoFisher) or Q5 Hot Start High-Fidelity 2x Master Mix (New England Biolabs). All plasmids used in this work were freshly prepared from 50 mL of Mach1 culture using ZymoPURE Plasmid Midiprep (Zymo Research Corporation) – a kit that involves an endotoxin removal procedure. Molecular biology grade, Hyclone water (GE Healthcare Life Sciences) was used in all assays, transfections, and PCR reactions to ensure exclusion of DNAse activity. Amino acid sequences for core ABE8 editors and sgRNA used can be found in Supplementary Sequence 2-9 and Supplementary Table 1.

### Generation of input bacterial TadA* libraries for directed evolution

The TadA*8.0 library was designed to encode all 20 amino acids at each amino acid position in the previously published^1^ TadA*7.10 open reading frame. Each TadA*8.0 library member contained ∼1-2 new coding mutations and was chemically synthesized and purchased from Ranomics Inc (Toronto, Canada). The TadA*8.0 library was PCR amplified with Phusion U Green MultiPlex PCR Master Mix and USER-assembled into the previously described^1^ bacterial vector optimized for ABE directed evolution.

### Bacterial evolution of TadA variants

Directed evolution of ABE containing the TadA*8 library was conducted as previously described^1^ with the following changes: 1. *E. coli* 10 betas (New England Biolabs) were used as the evolution host and 2. survival on kanamycin relied on correction of three genetic inactivating components (*e.g.* survival required reversion of two stop mutations and one active site mutation in kanamycin). The kanamycin resistance gene sequence, containing selection mutations, used for ABE8 evolution in this work can be found in Supplementary Sequence 1. After overnight co-culturing of selection plasmid and editor in 10 beta host cells, the library cultures were plated on 2xYT-agar medium supplemented with plasmid maintenance antibiotic and increasing concentrations of selection antibiotic, kanamycin (64-512 *μ*g/mL). Bacteria were allowed to grow for 1 day and the TadA*8 portion of the surviving clones were Sanger sequenced after enrichment, as previously described. Identified TadA*8 mutations of interest were then were then incorporated into mammalian expression vector via USER assembly.

### General HEK293T mammalian culture conditions

Cells were cultured at 37 °C with 5% CO_2_. HEK293T cells [CLBTx013, American Type Cell Culture Collection (ATCC)] were cultured in Dulbecco’s modified Eagles medium plus Glutamax(10566-016, Thermo Fisher Scientific) with 10% (v/v) fetal bovine serum (A31606-02, Thermo Fisher Scientific).

### Hek293T plasmid transfection and gDNA extraction

HEK293T cells were seeded onto 48-well well Poly-D-Lysine treated BioCoat plates (Corning) at a density of 35,000 cells/well and transfected 18-24 hours after plating. Cells were counted using a NucleoCounter NC-200 (Chemometec). To these cells were added 750 ng of base editor or nuclease control, 250 ng of sgRNA, and 10 ng of GFP-max plasmid (Lonza) diluted to 12.5 *μ*L total volume in Opti-MEM reduced serum media (ThermoFisher Scientific). The solution was combined with 1.5 *μ*L of Lipofectamine 2000 (ThermoFisher) in 11 *μ*L of Opti-MEM reduced serum media and left to rest at room temperature for 15 min. The entire 25 *μ*L mixture was then transferred to the pre-seeded Hek293T cells and left to incubate for ∼120 h. Following incubation, media was aspirated and cells were washed two times with 250 *μ*L of 1x PBS solution (ThermoFisher Scientific) and 100 *μ*L of freshly prepared lysis buffer was added (100 mM Tris-HCl, pH 7.0, 0.05% SDS, 25 *μ*g/mL Proteinase K (Thermo Fisher Scientific). Transfection plates containing lysis buffer were incubated at 37 °C for 1 hour and the mixture was transferred to a 96-well PCR plate and heated at 80 °C for 30 min.

### Hek293T mRNA lipofection

HEK293T cells were plated on 48-well poly-D-lysine coated plates (Corning) 16 to 20 hours before lipofection at a density of 30,000 cells per well in DMEM + Glutamax medium (Thermo Fisher Scientific) without antibiotics. 500 ng Cas9 or base editor expression mRNA was combined with 100ng of chemically modified synthetic sgRNA (2’-O-methyl analogs and 3’ phosphorothioate linkages at the first three 5’ and final three 3’ terminal RNA residues, Synthego) into a total volume of 15 μl with OPTIMEM + Glutamax. This was combined with 10 μl of lipid mixture, comprising 1.0 μl Lipofectamine MessengerMax and 9.0 μl OPTIMEM + Glutamax per well. Cells were harvested 3 days after transfection and either DNA or RNA was harvested and processed as described below.

### Next generation sequencing (NGS) of genomic DNA samples

Genomic DNA samples were amplified and prepared for high throughput sequencing as previously reported.^1^ Briefly, 1 *μ*L of gDNA was added to a 25 *μ*L PCR reaction containing Phusion U Green Multiplex PCR Master Mix and 0.5 *μ*M of each forward and reverse primer. Following amplification, PCR products were barcoded using unique Illumina barcoding primer pairs. Barcoding reactions contained 0.5 *μ*M of each illumina forward and reverse primer, 2 *μ*L of PCR mixture containing amplified genomic site of interest, and Q5 Hot Start High-Fidelity 2x Master Mix in a total volume of 25 *μ*L. All PCR conditions were carried out as previously published^1^. Primers used for site-specific mammalian cell genomic DNA amplification are listed in Supplementary Table 4. DNA concentration was quantified using a NanoDrop 1000 Spectrophotometer (ThermoFIsher Scientific) and sequenced on an Illumina MiSeq Instrument according to the manufacturer’s protocols.

### Targeted NGS data analysis

All targeted NGS data were analyzed by performing four general steps: (1) Illumina demultiplexing, (2) read trimming and filtering, (3) alignment of all reads to the expected amplicon sequence, and (4) generation of alignment statistics and quantification of editing rates. Each step is described in more detail in Supplementary Note 4. In an additional analysis, conducted using the same computational analysis approach, we show the haplotypes generated by ABE7 and ABE8 at different genetic loci (Supplementary Fig. 28).

### Treatment of HEK293T cells for whole genome sequencing, including preparation of genomic DNA and clonal isolation of edited cells

Cells were lipofected with base editor or Cas9-encoding mRNA combined an sgRNA targeting a region in B2M which, when successfully targeted by ABE, CBE or Cas9 leads to disruption of B2M (sgRNA target sequence: 5’-CTTACCCCACTTAACTATCT-3’^29^, Synthego) either through splice site disruption (ABE, Cas9) or incorporation of a stop codon (CBE), as described above. 24 hours post-transfection, cells were split 3:8 into a new plate to encourage cell growth. Three days post-transfection, HEK293T cells were harvested with TryplE Express (ThermoFisher), washed 1X with FACS buffer (PBS, 1% BSA, both ThermoFisher) and chilled at 4 °C for 15 minutes. The cells were then pelleted (1500 *g, 5 mins) and resuspended in a solution of FACS buffer with a 1:100 dilution of PE anti-human B2-microglobin (Biolegend 316306). Cells were incubated for 30 mins in the dark at 4 °C. Cells were then washed 3 times with FACS buffer by centrifugation (1500 *g, 5 mins) and resuspended in FACS buffer. Single, B2M-negative cells were sorted into 96-well plates except from untreated cells for which B2M-positive cells were sorted into 96-well plates. Representative FACS plots are shown in Supplementary Fig. 29. Nine days post sorting, wells were inspected and those containing single colonies were marked and treated with TryplE Express to promote cell growth. After four days of additional growth, genomic DNA was harvested from cells using Agincourt DNAdvance kit (Beckmann Coulter), according to the manufacturer’s instructions.

Genomic DNA was fragmented and adapter-ligated using the Nextera DNA Flex Library Prep Kit (Illumina) using the 96-well plate Nextera indexing primers (Illumina), according to the manufacturer’s instructions. Library size and concentration was confirmed by Fragment Analyzer (Agilent) and sent to Novogene for whole genome sequencing using an Illumina HiSeq.

### Analysis of whole transcriptome and whole genome sequencing data

All targeted NGS data were analyzed by performing four general steps: (1) alignment, (2) duplicate marking, (3) variant calling (4) background filtration of variants to remove artifacts and germline mutations. Each step is described in more detail in Supplementary Notes 5 and 6. The mutation reference and alternate alleles are reported relative to the plus strand of the reference genome.

### Analysis of DNA and RNA off-target editing for ABE architecture and ABE8 constructs

HEK293T cells were plated on 48-well poly-D-lysine coated plates (Corning) 16 to 20 hours before lipofection at a density of 30,000 cells per well in DMEM + Glutamax medium (Thermo Fisher Scientific) without antibiotics. 750 ng nickase or base editor expression plasmid DNA was combined with 250ng of sgRNA expression plasmid DNA in 15 μl OPTIMEM + Glutamax. This was combined with 10 μl of lipid mixture, comprising 1.5 μl Lipofectamine 2000 and 8.5 μl OPTIMEM + Glutamax per well. Cells were harvested 3 days after transfection and either DNA or RNA was harvested. For DNA analysis, cells were washed once in 1X PBS, and then lysed in 100 μl QuickExtract™ Buffer (Lucigen) according to the manufacturer’s instructions. For RNA harvest, the MagMAX™ mirVana™ Total RNA Isolation Kit (Thermo Fisher Scientific) was used with the KingFisher™ Flex Purification System according to the manufacturer’s instructions.

Targeted RNA sequencing was performed largely as previously described.^1^ cDNA from the isolated RNA using the SuperScript IV One-Step RT-PCR System with EZDnase (Thermo Fisher Scientific) according to the manufacturer’s instructions. The following program was used: 58 °C for 12 min; 98 °C for 2 min; followed by PCR cycles that varied by amplicon: for CTNNB1 and IP90: 32 cycles of [98 °C for 10 s; 60 °C for 10 sec; 72 °C for 30 sec]. No RT controls were run concurrently with the samples. Following the combined RT-PCR, amplicons were barcoded and sequenced using an Illumina Miseq as described above. The first 125nt in each amplicon, beginning at the first base after the end of the forward primer in each amplicon, was aligned to a reference sequence and used for our analysis of maximum A-to-I frequencies in each amplicon.

Off-target DNA sequencing was performed using previously published primers^1,2^ listed in Supplementary Table 4 using a two-step PCR and barcoding method to prepare samples for sequencing using Illumina Miseq sequencers as above.

### mRNA production for ABE editors used in CD34+ cells, T cells and HEK293T cells

All adenine base editor mRNA was generated using the following synthesis protocol. Editors were cloned into a plasmid encoding a dT7 promoter followed by a 5’UTR, Kozak sequence, ORF, and 3’UTR. The dT7 promoter carries an inactivating point mutation within the T7 promoter that prevents transcription from circular plasmid. This plasmid templated a PCR reaction (Q5 Hot Start 2X Master Mix), in which the forward primer corrected the SNP within the T7 promoter and the reverse primer appended a polyA tail to the 3’ UTR. The resulting PCR product was purified on a Zymo Research 25ug DCC column and used as mRNA template in the subsequent in vitro transcription. The NEB HiScribe High-Yield Kit was used as per the instruction manual but with full substitution of N1-methyl-pseudouridine for uridine and co-transcriptional capping with CleanCap AG (Trilink). Reaction cleanup was performed by lithium chloride precipitation. Primers used for amplification can be found in Supplementary Table 3.

The Cas9 mRNA used here was purchased from Trilink (CleanCap Cas9 mRNA 5moU) and the CBE mRNA used in the whole genome sequencing experiment was generated in-house.

### CD34+ cell preparation

Mobilized peripheral blood was obtained and enriched for Human CD34+ HSPCs and frozen in single-use aliquots (HemaCare, M001F-GCSF/MOZ-2). The CD34+ cells were thawed and put into X-VIVO 10 (Lonza) containing 1% Glutamax (Gibco), 100ng/mL of TPO (Peprotech), SCF (Peprotech) and Flt-3 (Peprotech) and cultured for 48 hours prior to electroporation.

### Electroporation of CD34+ cells

48 hours post thaw, the cells were spun down to remove the X-VIVO 10 media and washed in MaxCyte buffer (HyClone) with 0.1% HSA (Akron Biotechnologies). The cells were then resuspended in cold MaxCyte buffer at 1,250,000 cell per mL and split into multiple 20µL aliquots. The ABE mRNA (0.15 *μ*M) and −198 HBG1/2 sgRNA (4.05 *μ*M) were then aliquoted as per the experimental conditions and raised to a total of 5µL in MaxCyte buffer. 20µL of cells was then added into the 5µL RNA mixture in groups of 3 and loaded into each chamber of an OC25×3 MaxCyte cuvette for electroporation and electroporated with program “HSC-4”. After receiving the charge, 25µL was collected from the chambers and placed in the center of the wells in a 24-well untreated culture plate. The cells recovered for 20 minutes in an incubator (37°C, 5% CO_2_). After the 20 minutes recovery, X-VIVO 10 containing 1% Glutamax, 100ng/mL of TPO, SCF and Flt-3 was added to the cells for a concentration of 1,000,000 cells per mL. The cells were then left to further recover in an incubator (37°C, 5% CO_2_) for 48hrs.

### Erythrocyte differentiation post ABE electroporation

Following 48 h post electroporation rest (day 0 of culture), the cells were spun down and moved to “Phase 1” IMDM media (ATCC) containing 5% human serum, 330µg/mL transferrin (Sigma), 10µg/mL human insulin (Sigma), 2U/mL heparin sodium (Sigma), 3U/mL EPO (Peprotech), 100ng/mL SCF (Peprotech), 5µg/mL IL3 and 50µM hydrocortisone (Sigma) at 20,000 cells per mL. On day 4 of culture, the cells were fed 4x volume of the same media. On day 7, the cells were spun down and moved to “Phase 2” IMDM media containing 5% human serum (Sigma), 330µg/mL transferrin, 10µg/mL human insulin, 2U/mL heparin sodium, 3U/mL EPO and 100ng/mL SCF at 200,000 cells per mL. On day 11, cells were spun down and moved to “Phase 3” IMDM media containing 5% human serum, 330µg/mL of transferrin, 10µg/mL human insulin, 2U/mL of heparin sodium and 3U/mL of EPO at 1,000,000 cells per mL. On day 14, the cells were spun down and resuspended in the same media as day 11 but at 5,000,000 cells per mL. On day 18, the differentiated red blood cells were collected in 500,000 cell aliquots, washed once in 500µL DPBS (Gibco) and frozen at −80°C for 24 hours before UHPLC processing.

### Preparation of red blood cell sample for UHPLC analysis

Frozen red blood cell pellets were thawed at room temperature. Pellets were diluted to a concentration of 5 × 10^4^ cells/μL with ACK lysis buffer (Gibco). Samples were mixed by pipette and incubated at room temperature for 5 min. Samples were then frozen at −80 °C for 5 min, allowed to thaw, and vortex mixed for 30 seconds. Samples were then lysed by centrifugation at 6,700 g for 10 min. The supernatant was carefully removed (without disturbing cell debris pellet) and transferred to a new plate where a 10-fold dilution in ultrapure water was done for UHPLC analysis.

### Ultra-high performance liquid chromatography (UHPLC) Analysis

Reverse-phase separation of globin chains was performed using a UHPLC system configured with a binary pump and UV detector (Thermo Fisher Scientific, Vanquish Horizon). The Waters AQUITY Peptide BEH C18 VanGuard pre-column (2.1 x 5 mm, 1.7 μm beads, 300 Å pore size) followed by ACQUITY Peptide BEH C18 Column (2.1 x 150 mm, 1.7 μm beads, 300 Å pore size) (Waters Corp) were used for the separation with a column temperature of 60 °C. Elution was preformed using 0.1% trifluoroacetic acid (TFA) in water (**A**) and 0.08% TFA in acetonitrile (**B**) with a flow rate of 0.25 mL/min. Separation of the globin chains was achieved using a linear gradient of 40-52%**B** 0-10 min; 52-40%**B** 10-10.5 min; and 40%**B** to 12 min. Sample injection volume was 10μL. UV spectra at a wavelength of 220nm with a data rate of 5Hz was collected throughout the analysis. Globin chain identities were confirmed through LC/MS analysis of hemoglobin standards.

### Genomic DNA extraction for CD34+ cells

Immediately following ABE electroporation, an aliquot of cells was cultured in X-VIVO 10 media (Lonza) containing 1% Glutamax (Gibco), 100ng/mL of TPO (Peprotech), SCF (Peprotech) and Flt-3 (Peprotech). Following 48 h and 144 h post culturing, 100,000 cells were collected and spun down. 50 µL of Quick Extract (Lucigen) was added to the cell pellet and the cell mixture was transferred to a 96-well PCR plate (Bio-Rad). The lysate was heated for 15 minutes at 65°C followed by 10 minutes at 98°C. The cell lysates were stored at −20°C.

### Generation of T cells

Frozen, bulk PBMCs obtained from healthy donors were thawed and cultured in a T-cell growth media (TCGM) consisting of X-VIVO15 (Lonza) supplemented with 5% human serum, type AB (Valley Biomedical), 2mM of GlutaMAX (Gibco), 10mM of HEPES buffer solution (Gibco), and 250IU/mL of recombinant human interleukin-2 (rhIL-2, CellGenix GmbH). Cells were activated with soluble human anti-CD3 (clone OKT3, Miltenyi Biotec) and human anti-CD28 (clone 15E8, Miltenyi Biotec) and cultured at 37°C in a 5% CO_2_ incubator.

### Electroporation of primary human T cells

At either 72hr or 96hr post T cell activation, cells were spun down at 500g for 5 mins. Supernatant was removed and cells were then washed once with DPBS (Gibco) and spun again. DPBS was removed and cells were resuspended in P3 primary cell electroporation buffer (Lonza) at a concentration of 50e6 cells/mL. Two micrograms of ABE8 mRNA and one microgram of 5’/3’ end-modified sgRNA (Synthego) were added to 1e6 cells (20uL), that were then electroporated using a Lonza 4-D Nucleofector with 96-well Shuttle™ add-on (Lonza). Sequences of sgRNA can be found in Supplementary Table 2. Post electroporation, 100uL of TCGM media was used to quench the reaction, and cells were subsequently transferred to a single well of a G-Rex® 24-well plate (Wilson Wolf) containing 8mL of pre-warmed TCGM +IL-2. Plates were then placed in an incubator (37°C, 5% CO_2_) until further analysis.

### Flow Cytometry of primary human T cells

To assess editing efficiency,1 × 10^6^ cells were taken from culture five days post electroporation and stained with the following primary anti-human antibodies: Cbl-b (Clone D3C12, Cell Signaling Technologies) followed by AlexaFluor 647 F(ab’)2 goat anti rabbit IgG (H+L) (Invitrogen), CD3 (Clone UCHT1, PE, Biolegend) CD7 (Clone CD7-6B7, FITC, Biolegend), HLA-DR (Clone L243, PE Biolegend), B2M (Clone 2M2, PE, Biolegend), CD279 (Clone eBioJ105, PE, Biolegend). Data was acquired using an Attune NxT Flow Cytometer and analyzed using FlowJo Single Cell Analysis Software v10.6.1 (FlowJo, LLC). Examples of gating strategies have been included in Supplementary Figure 30.

### Genomic DNA extraction for human T cells

Following incubation, ∼1 × 10^6^ of treated T cells were spun down, washed with PBS and resuspended in 200 *μ*L of Quick Extract (Lucigen) lysis buffer and cells were lysed according to the manufacture’s protocol. Genomic DNA was directly used in subsequent PCR amplification steps.

## Data availability

Plasmids encoding the core ABE8s used in this work are available through Addgene. High-throughput sequencing data are deposited in the NCBI Sequence Read Archive (PRJNA574182).

## Code accessibility

All software tools used for data analysis are publicly available. Detailed information about versions and parameters used, as well as shell commands, are provided in Supplementary Note 4.

