## Supplementary Information for "Directed Evolution of Adenine Base Editors with Increased Activity and Therapeutic Application"

Beam Therapeutics
Cambridge, MA, 02139

*Correspondence should be addressed to:

Nicole M. Gaudelli:
Giuseppe Ciaramella:

### Table of Contents

**Supplementary Figure 1:** Genotypes of all ABE8s described in this work…………………………………………………….4
**Supplementary Figure 2:** Three perspectives of the *E. coli* TadA deaminase aligned with the *S. aureus* TadA complexed with tRNA$^{Arg2}$ ……………………………………………………………………………………………………………...…5
**Supplementary Figure 3:** Median A•T to G•C conversion and corresponding INDEL formation of TadA, C-terminal alpha-helix truncation ABE constructs in HEK293T cells ……………………………………………………………………….6
**Supplementary Figure 4**: Comparison between the on- and off-target editing frequencies between ABE7.10, ABEmax and ABEmax with one BPNLS ……………………………………………………………………………………………...7
**Supplementary Figure 5**: Median A•T to G•C conversion of all 40 ABE8 constructs across 8 genomic sites ……………….8
**Supplementary Figure 6**: Median indel % of all 40 ABE8 constructs across 8 genomic sites ………………….……………9
**Supplementary Figure 7**: Dendrogram of ABE8s ………………………………………………………………………………10
**Supplementary Figure 8:** Median A•T to G•C conversion of core eight ABE8 constructs across 8 genomic sites …..…...11
**Supplementary Figure 9** : Median indel frequency of core 8 ABE8s tested at 8 genomic sites ……………………………12
**Supplementary Figure 10:** Median A•T to G•C conversion of core NG-ABE8 constructs 9 (-NG PAM) ………………..…13
**Supplementary Figure 11**: Median indel frequency of core NG-ABE8s tested ……………..……………………………..……14
**Supplementary Figure 12:** Median A•T to G•C conversion of core Sa-ABE8 constructs (-NNGRRT PAM) ……………...15
**Supplementary Figure 13**: Median indel frequency of core Sa-ABE8s tested ……………………..………………………..16
**Supplementary Figure 14:** Median A•T to G•C conversion of core dC9-ABE8-m constructs ……………..…………………17
**Supplementary Figure 15:** Median A•T to G•C conversion of core dC9-ABE8-d constructs ……………………………..…18
**Supplementary Figure 16**: Median indel frequency of core dC9-ABE8s tested ……………..………………………………19
**Supplementary Figure 17:** C•G to T•A editing with Hek293T cells treated with ABE8s and ABE7.10 …………………….20
**Supplementary Figure 18:** A•T to G•C conversion of CD34+ cells treated with ABE8 at the -198 promoter site upstream of HBG1/2 ……………………………………………………………………………………………………………………...……21
**Supplementary Figure 19:** Indel frequency of CD34+ cells treated with ABE8 at the -198 site of the γ-globin promoter……21
**Supplementary Figure 20**: Median gamma-globin levels across all ABE8-treated vs. ABE7-treated samples ……………..…22
**Supplementary Figure 21:** Protein knockdown measured by flow cytometry by ABE editors in primary T cells ……………23
**Supplementary Figure 22**: Variant alleles created by ABEs at six different targets in primary T cells ……………………..24
**Supplementary Figure 23:** Median indel frequencies at 12 previously identified sgRNA-dependent Cas9 off-target loci in human cells ……………………………………………………………...………………………………………………25
**Supplementary Figure 24:** DNA on-target, sgRNA-dependent DNA off-target editing and sgRNA-independent off-target mRNA editing by ABE8 constructs containing mutations designed to reduced spurious cellular RNA editing……........……26
**Supplementary Figure 25:** Median DNA on-target and sgRNA-dependent DNA off-target editing by ABE8 constructs using plasmid or mRNA delivery ……………………………..……………………………...……………………………….27
**Supplementary Figure 26:** Whole transcriptome sequencing in HEK293T cells treated with indicated mRNA …………..28
**Supplementary Figure 27:** Mutational classification plots for each sample sent for whole genome sequencing …………29
**Supplementary Figure 28**: Alleles created by ABEs across 8 different genomic sites in HEK293Ts ………………………..30

### Table of Contents Continued

**Supplementary Figure 29:** Representative examples of gates used to flow sort B2M-positive and B2M-negative cells prior to whole genome sequencing ……………………………………………………………………………..………………31
**Supplementary Figure 30:** Examples of gates used for assessment of protein knockdown in T cells ……….……………. 32

**Supplementary Table 1**: Sequences of sgRNA in Hek293T cells ……………………………………………………...33
**Supplementary Table 2**: Primers used for *in vitro* transcription of ABE8s ……………………………………………….34
**Supplementary Table 3**: Sequences of sgRNAs used in primary cells ………………………………………………….34
**Supplementary Table 4**: Primers used for mammalian cell genomic DNA and RNA amplification ……….……..……35-37
**Supplementary Table 5-25**: UHPLC traces for gamma globin quantification and integration tables ………..………..…38-59
**Supplementary Table 26**: P-values resulting from comparisons between relative cytosine and adenine deamination-induced mutation frequencies amongst treatment groups subjected to whole genome sequencing……………………….60
**Supplementary Table 27**: Total number of mutations, relative to untreated control, detected in each whole genome sequencing sample………………………………………………………………………………………………….61

**Supplementary Note 1**: Discussion of codon optimization and NLS choice for ABE8 designs …………….………..….62
**Supplementary Note 2**: Modeling $\gamma$-globin induction as a function of editor-induced allele frequencies …………………..63
**Supplementary Note 3:** Discussion of whole genome sequencing data ………………….…………………..………64-65
**Supplementary Note 4:** Targeted NGS analysis details…………………..………………………………....………66-67
**Supplementary Note 5:** Whole transcriptome sequencing analysis details……….……….……..…..………...……68
**Supplementary Note 6:** Whole genome sequencing analysis details…………………..……………………….…….…69

**Supplementary Sequence 1**: DNA sequence of inactivated kanamycin resistance gene used for ABE8 evolution ……….70
**Supplementary Sequence 2-9:** Amino acid sequences of core ABE8 editors ……………………………………….……71-75

**Supplementary References**: ……………………...…………………….....…………………………………………...………76

| | residue identity in evolved TadA | | | | | | | | | | | | | | | | | |
|---|---|---|---|---|---|---|---|---|---|---|---|---|---|---|---|---|---|---|
| | 23 | 36 | 48 | 51 | 76 | 82 | 84 | 106 | 108 | 123 | 146 | 147 | 152 | 154 | 155 | 156 | 157 | 166 |
| **ABE7.10** | **R** | **L** | **A** | **L** | **I** | **V** | **F** | **V** | **N** | **Y** | **C** | **Y** | **P** | **Q** | **V** | **F** | **N** | **T** |
| ABE8.1-m | | | | | | | | | | | | T | | | | | | |
| ABE8.2-m | | | | | | | | | | | | R | | | | | | |
| ABE8.3-m | | | | | | | | | | | | | | S | | | | |
| ABE8.4-m | | | | | | | | | | H | | | | | | | | |
| ABE8.5-m | | | | | | S | | | | | | | | | | | | |
| ABE8.6-m | | | | | | | | | | | | | | | | | | R |
| ABE8.7-m | | | | | | | | | | | | | | R | | | | |
| ABE8.8-m | | | | | | | | | | H | | R | | R | | | | |
| ABE8.9-m | | | | | Y | | | | | | | R | | R | | | | |
| ABE8.10-m | | | | | | | | | | | | R | | R | | | | R |
| ABE8.11-m | | | | | | | | | | | | T | | R | | | | |
| ABE8.12-m | | | | | | | | | | | | T | | S | | | | |
| ABE8.13-m | | | | | Y | | | | | H | | R | | R | | | | |
| ABE8.14-m | | | | | Y | S | | | | | | | | | | | | |
| ABE8.15-m | | | | | | S | | | | | | R | | | | | | |
| ABE8.16-m | | | | | | S | | | | H | | R | | | | | | |
| ABE8.17-m | | | | | | S | | | | | | | | R | | | | |
| ABE8.18-m | | | | | | S | | | | H | | | | R | | | | |
| ABE8.19-m | | | | | | S | | | | H | | R | | R | | | | |
| ABE8.20-m | | | | | Y | S | | | | H | | R | | R | | | | |
| ABE8.1-d | | | | | | | | | | | | T | | | | | | |
| ABE8.2-d | | | | | | | | | | | | R | | | | | | |
| ABE8.3-d | | | | | | | | | | | | | | S | | | | |
| ABE8.4-d | | | | | | | | | | H | | | | | | | | |
| ABE8.5-d | | | | | | S | | | | | | | | | | | | |
| ABE8.6-d | | | | | | | | | | | | | | | | | | R |
| ABE8.7-d | | | | | | | | | | | | | | R | | | | |
| ABE8.8-d | | | | | | | | | | H | | R | | R | | | | |
| ABE8.9-d | | | | | Y | | | | | | | R | | R | | | | |
| ABE8.10-d | | | | | | | | | | | | R | | R | | | | R |
| ABE8.11-d | | | | | | | | | | | | T | | R | | | | |
| ABE8.12-d | | | | | | | | | | | | T | | S | | | | |
| ABE8.13-d | | | | | Y | | | | | H | | R | | R | | | | |
| ABE8.14-d | | | | | Y | S | | | | | | | | | | | | |
| ABE8.15-d | | | | | | S | | | | | | R | | | | | | |
| ABE8.16-d | | | | | | S | | | | H | | R | | | | | | |
| ABE8.17-d | | | | | | S | | | | | | | | R | | | | |
| ABE8.18-d | | | | | | S | | | | H | | | | R | | | | |
| ABE8.19-d | | | | | | S | | | | H | | R | | R | | | | |
| ABE8.20-d | | | | | Y | S | | | | H | | R | | R | | | | |

**Supplementary Figure 1 | Genotypes of 40 ABE8s described in this work.** Residue position in the evolved *E. coli* TadA portion of ABE are indicated. Mutational changes in ABE8 are shown when distinct from ABE7.10 mutations. ABE8s highlighted are the “core 8” chosen for further studies.

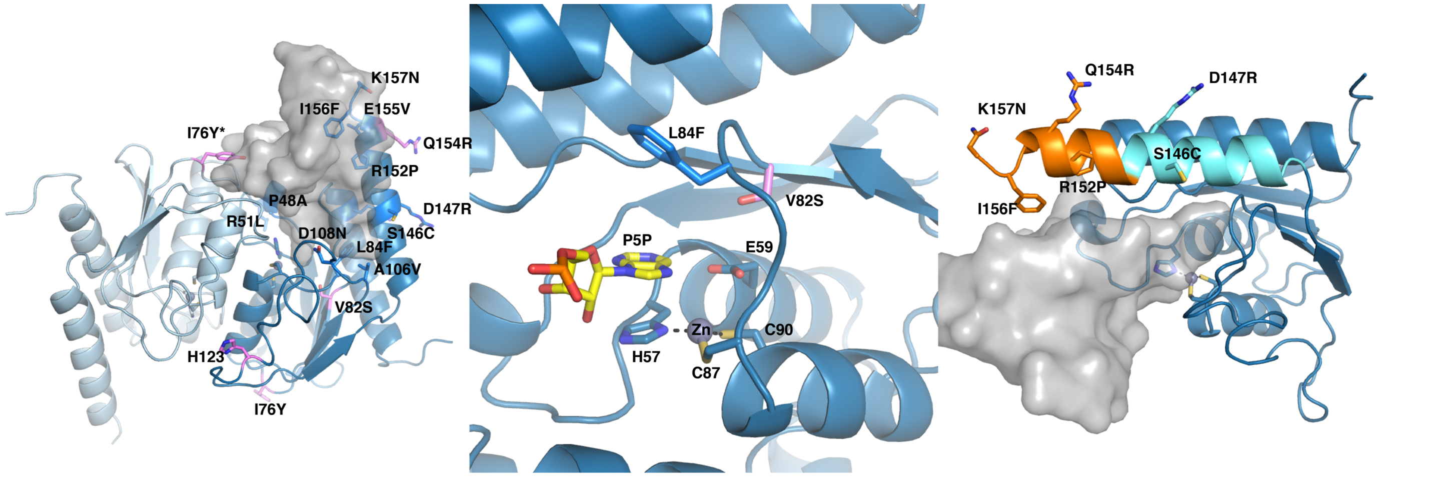


**Supplementary Figure 2 | Three perspectives of the *E. coli* TadA deaminase (PDB 1Z3A) aligned with the *S. aureus* TadA (not shown) complexed with tRNA$^{Arg2}$ (PDB 2B3J).** Mutations identified in the eighth round of evolution are highlighted in pink. Key residues throughout ABE8 and the active site are labeled. The region of the C-terminal alpha helix investigated in this study is highlighted in orange.

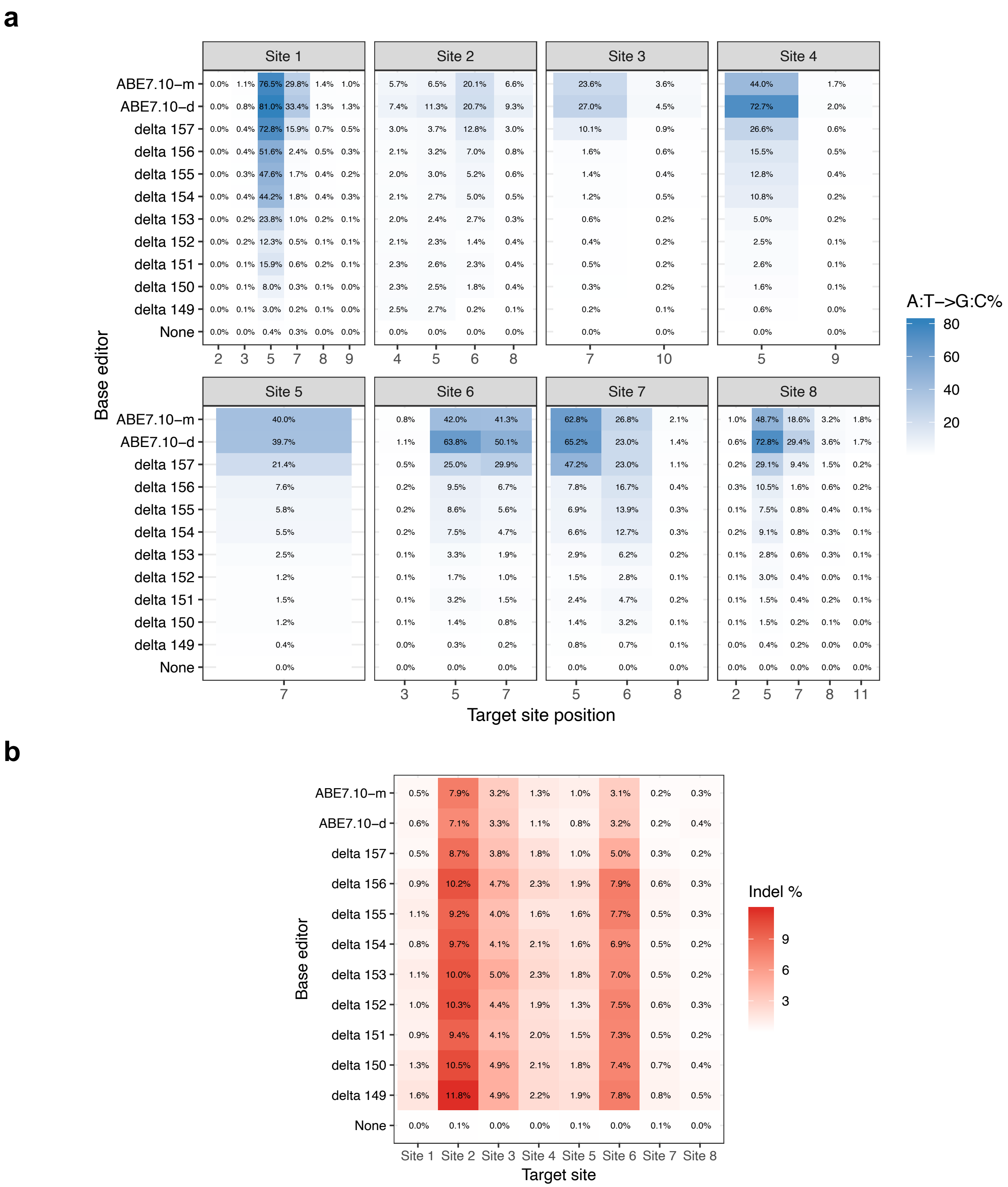

**Supplementary Figure 3 | Median A•T to G•C conversion and corresponding indel formation of TadA, C-terminal alpha-helix truncation ABE constructs in HEK293T cells. a,** A•T to G•C median editing conversion across 8 genomic sites and **b,** Indel formation. Delta residue values correspond to deletion position in TadA. Median values are shown, generated from n=3 biological replicate.

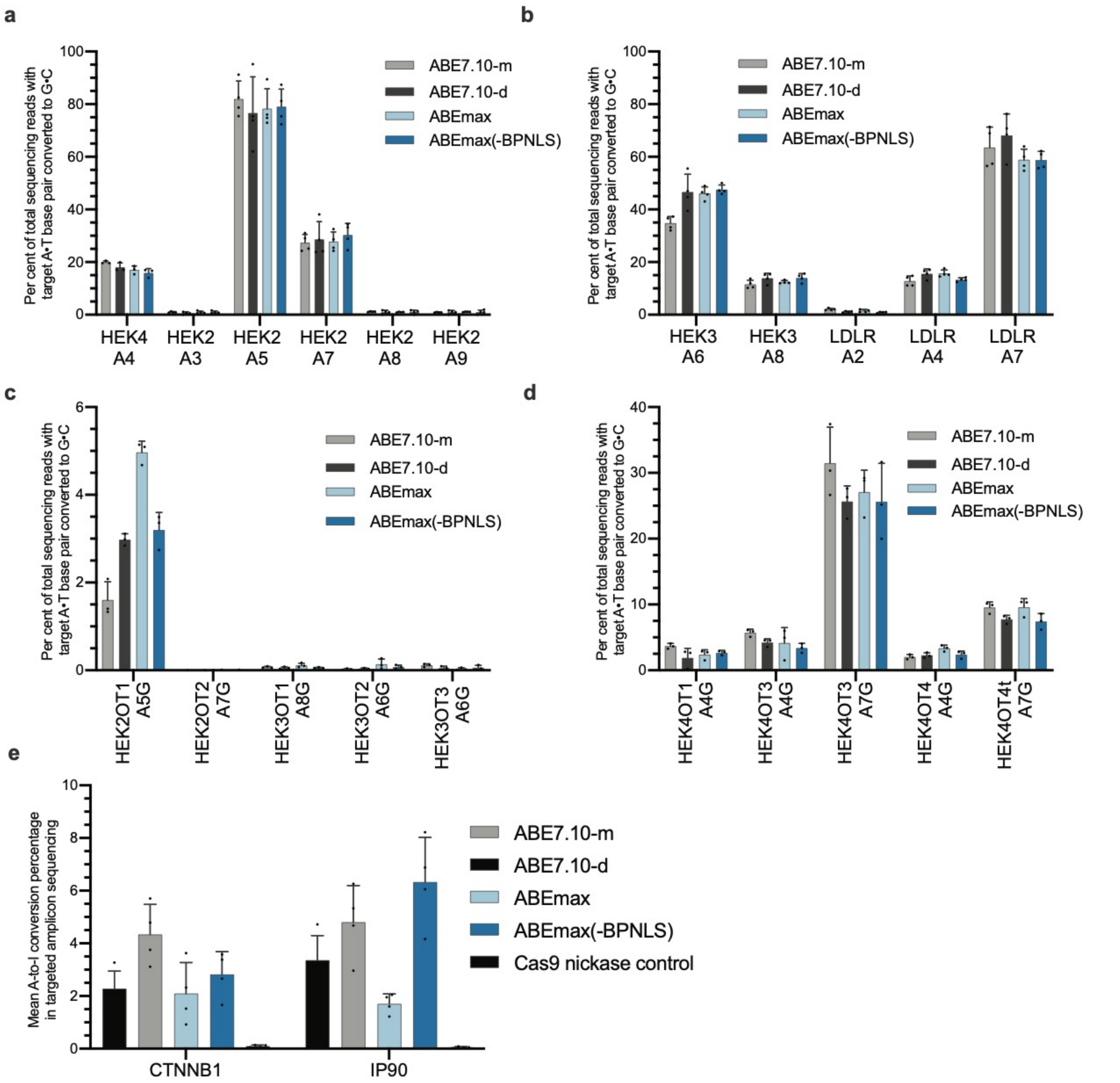


**Supplementary Figure 4 | Comparison between the on- and off-target editing frequencies between ABE7.10, ABEmax and ABEmax with one BPNLS in Hek293T cells**. ABEmax(-BPNLS) is ABEmax with only a single BPNLS at the C-terminal end of the construct. **a** and **b**, on-target DNA editing frequencies, **c** and **d**, sgRNA-guided DNA-off-target editing frequencies and **e,** Maximum cellular RNA off-target editing frequencies. Individual data points are shown and error bars represent s.d. for n=3 or n=4 independent biological replicates, as can be determined for each sample by the number of included data points on the graph. The bar for each condition represents the mean value. Replicates were performed on different days. See also Supplementary Note 1.

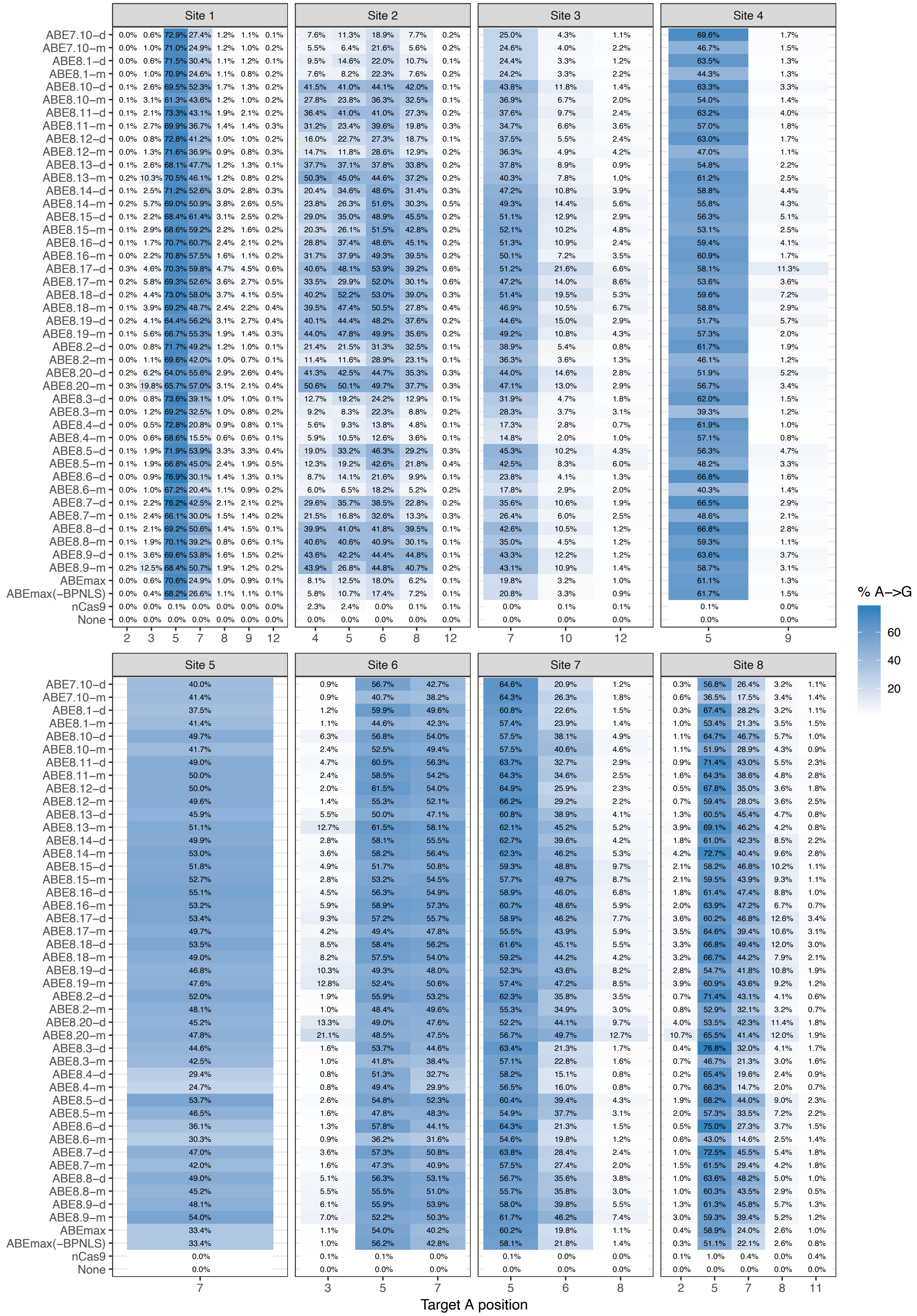


**Supplementary Figure 5: Median A•T to G•C conversion of all 40 ABE8 constructs in HEK293T cells across 8 genomic sites.** Median values were determined from n=3 or greater biological replicates.

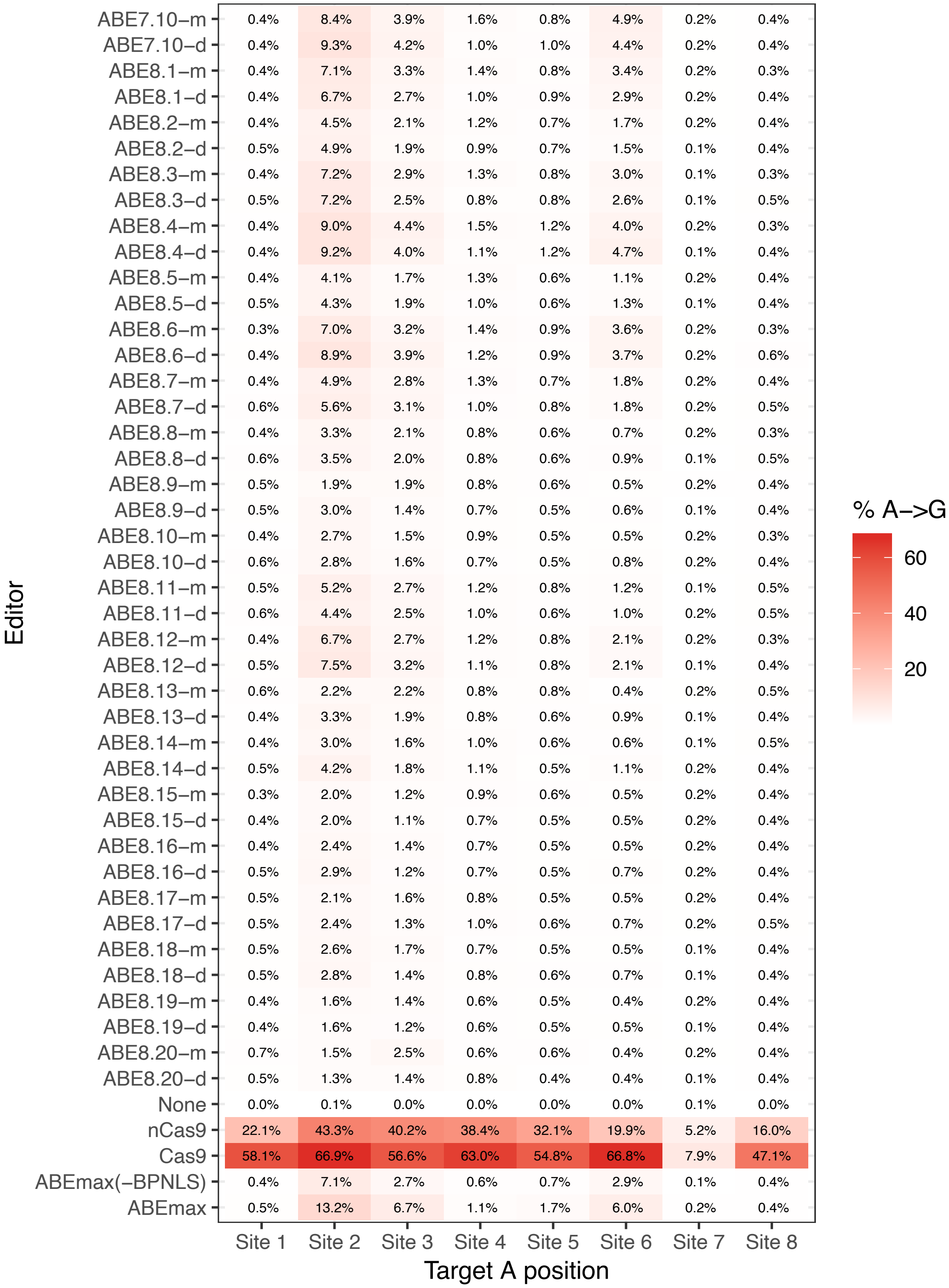


**Supplementary Figure 6 | Median indel % of all 40 ABE8 constructs in HEK293T cells across 8 genomic sites.** Median values were determined from two or greater biological replicates

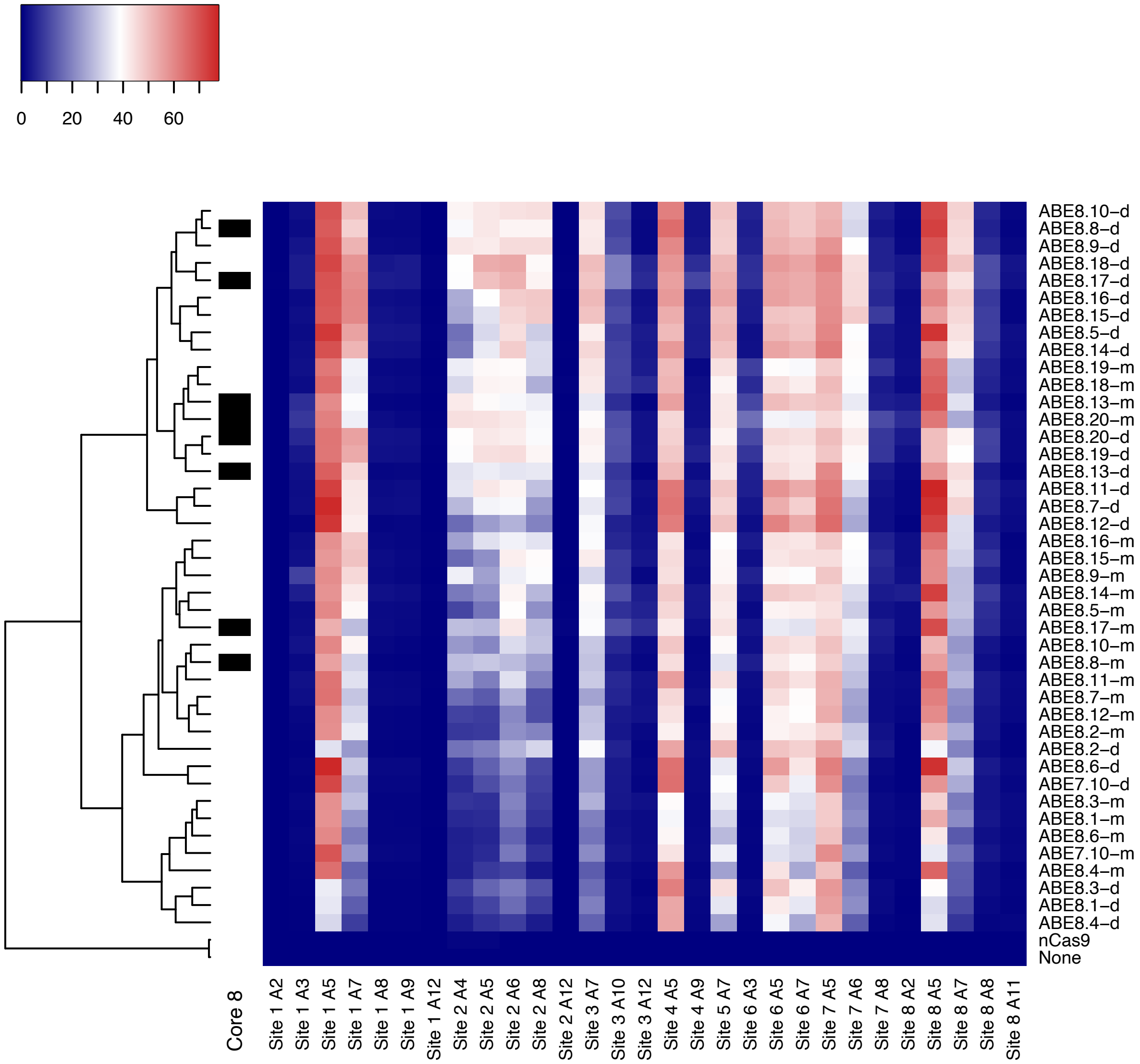


**Supplementary Figure 7 | Dendrogram of ABE8s.** Core ABE8 constructs selected for further studies highlighted in black

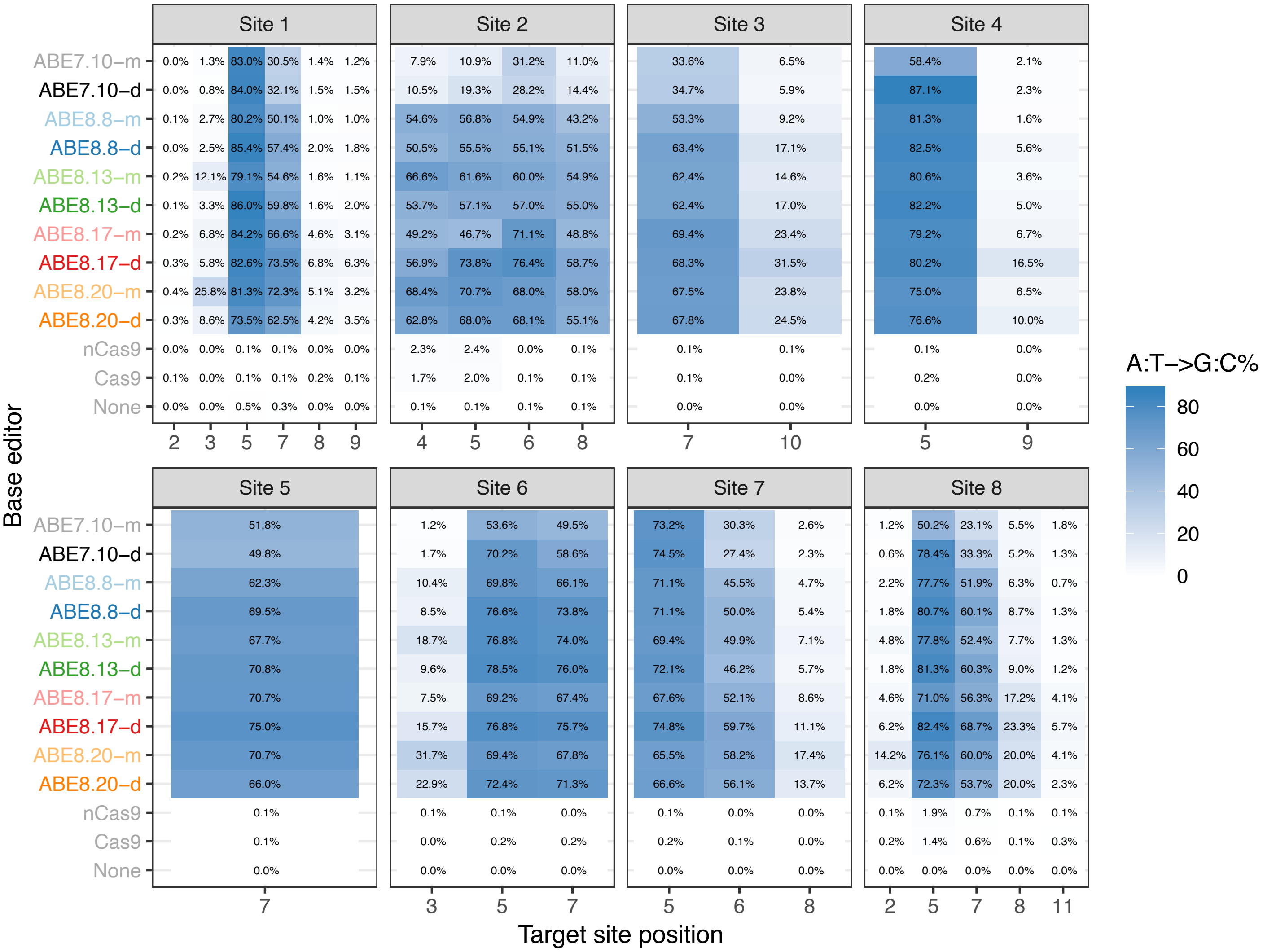

**Supplementary Figure 8: Median A•T to G•C conversion of core eight ABE8 constructs in HEK293T cells across eight genomic sites.** Median values were determined from n=3 or greater biological replicates.

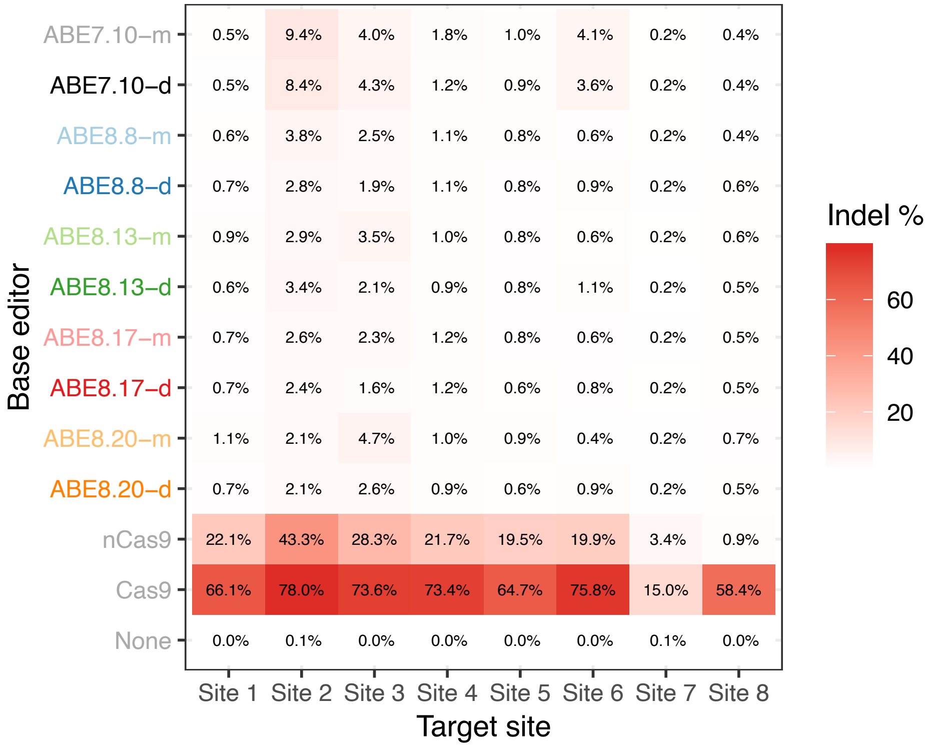


**Supplementary Figure 9 | Median indel frequency of core 8 ABE8s tested at 8 genomic sites in HEK293T cells.** Median values were determined from n=3 or greater biological replicates.

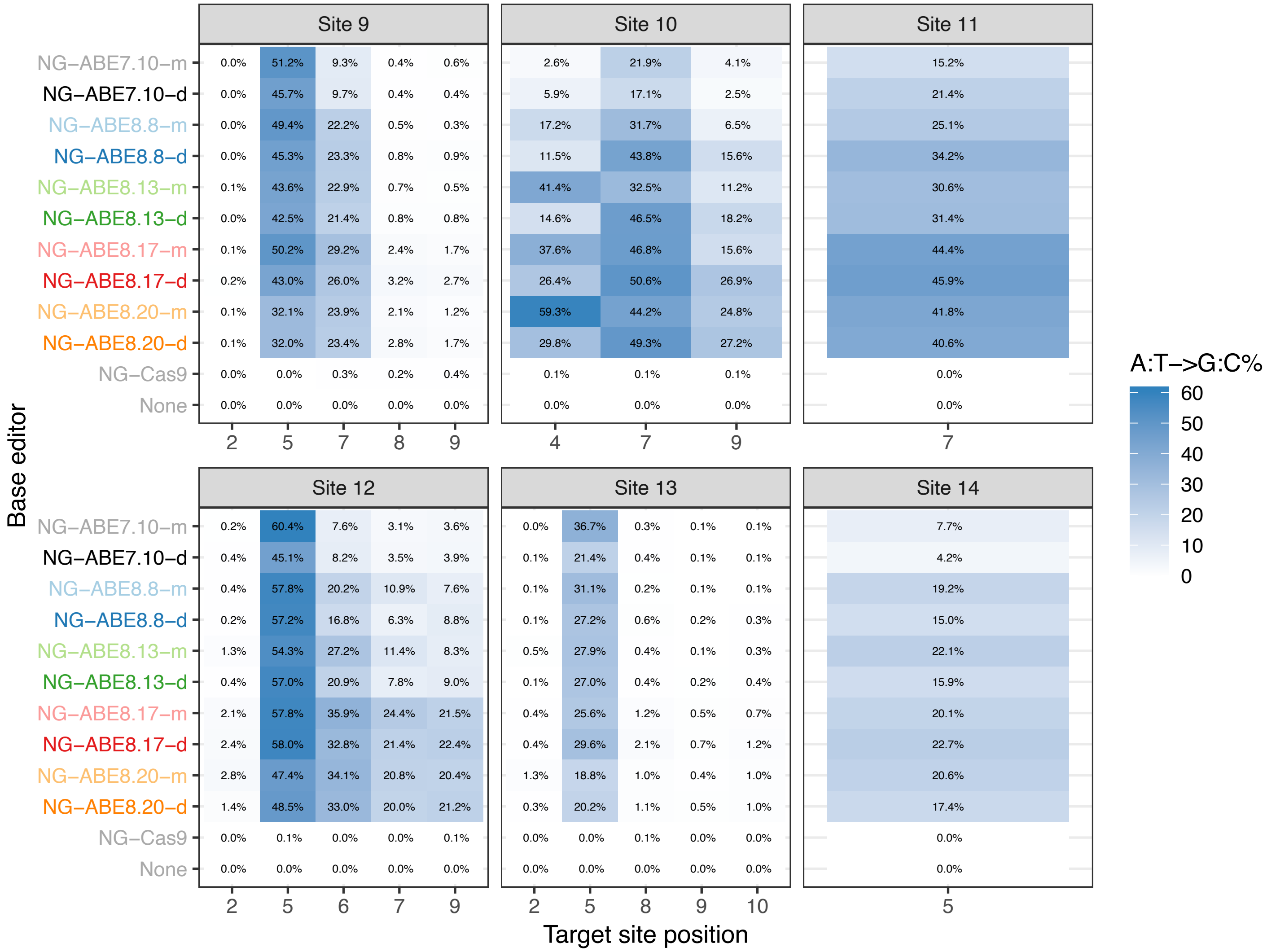


**Supplementary Figure 10 | Median A•T to G•C conversion of core NG-ABE8 constructs (-NG PAM) at six genomic sites in HEK293T cells.** Median values are shown, generated from n=3 biological replicates.

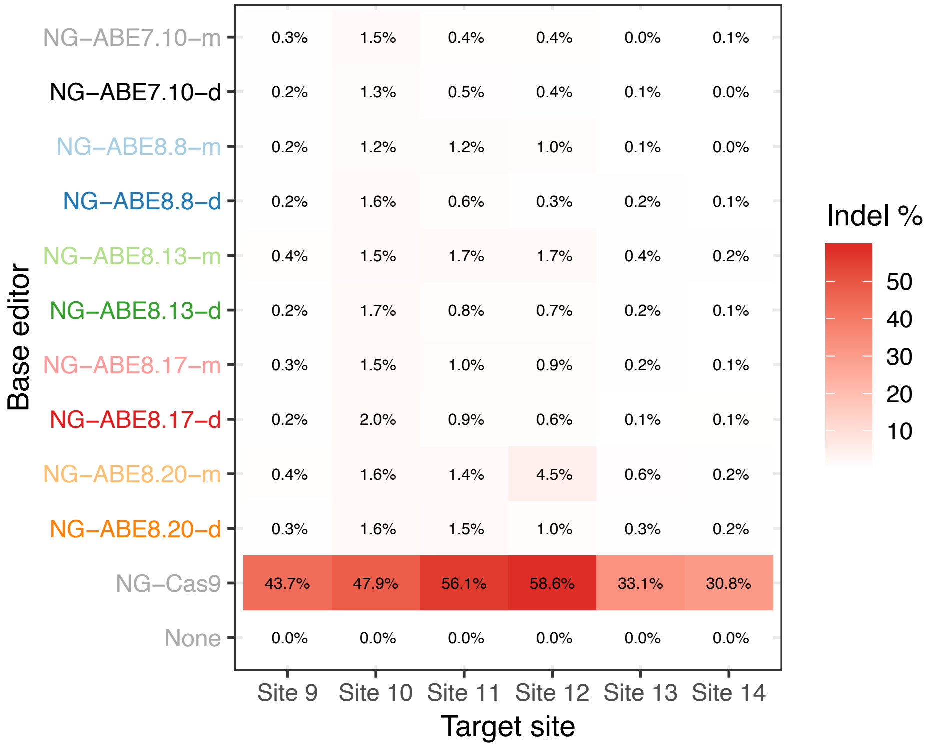


**Supplementary Figure 11 | Median indel frequency of core NG-ABE8s tested at six genomic sites in HEK293T cells.** Median values are shown, generated from n=3 biological replicates.

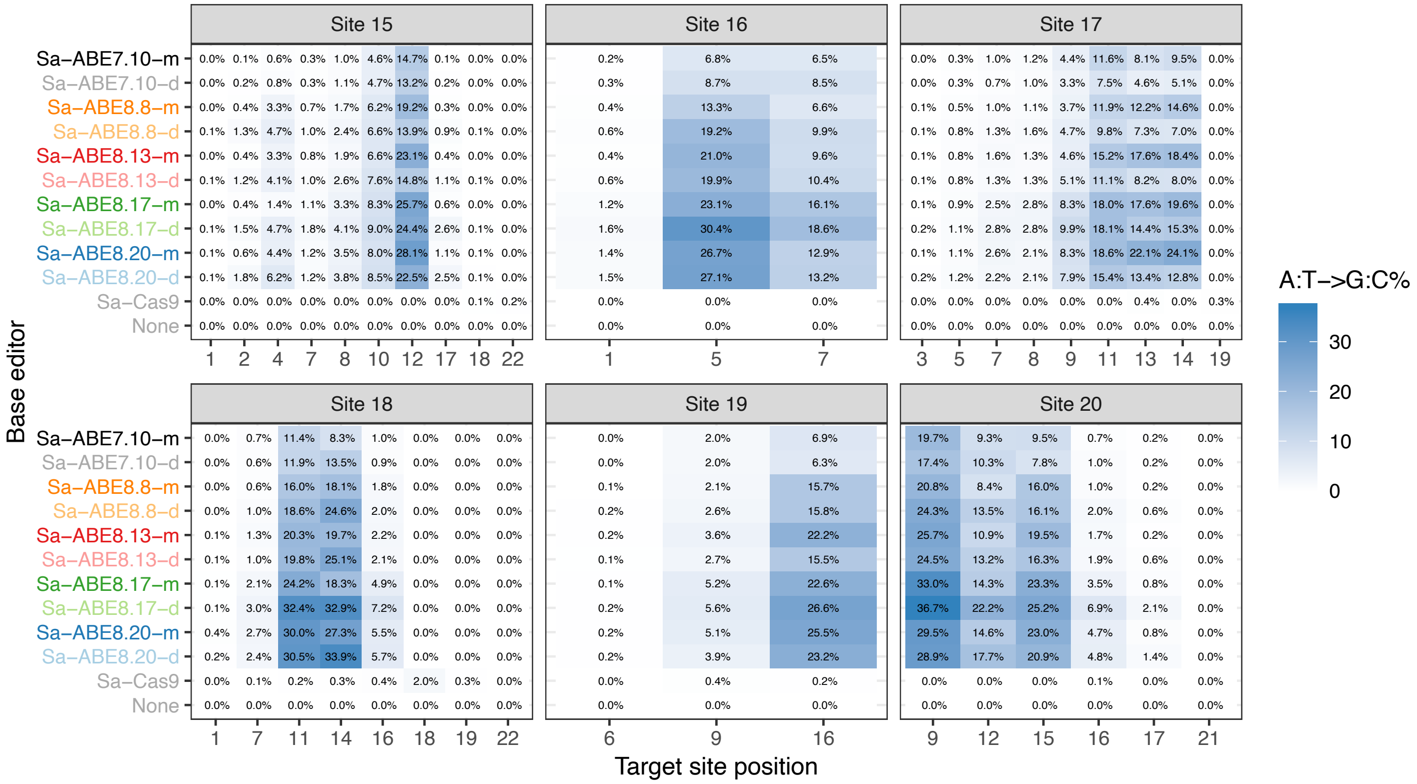


**Supplementary Figure 12 | Median A•T to G•C conversion of core Sa-ABE8 constructs (-NNGRRT PAM) at six genomic sites in HEK293T cells.** Site positions are numbered -2 to 20 (5' to 3') within the 22-nt protospacer. Position 20 is 5' to the NNGRRT PAM. Median values are shown, generated from n=3 biological replicates.

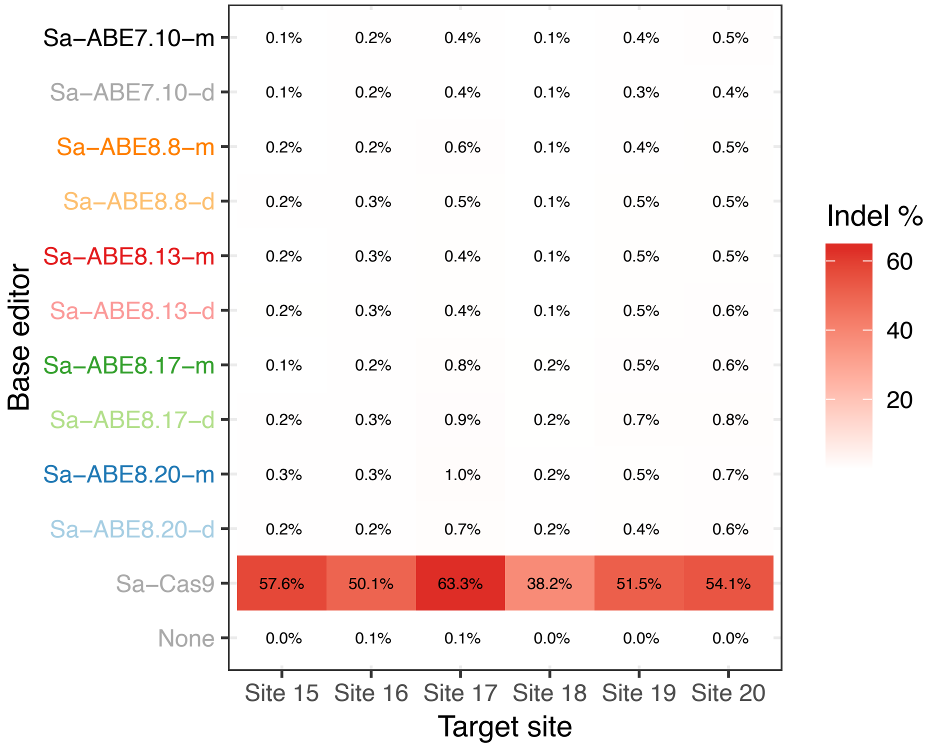


**Supplementary Figure 13 | Median indel frequency of core Sa-ABE8s tested at 8 genomic sites in HEK293T cells.** Median values are shown, generated from n=3 biological replicates.

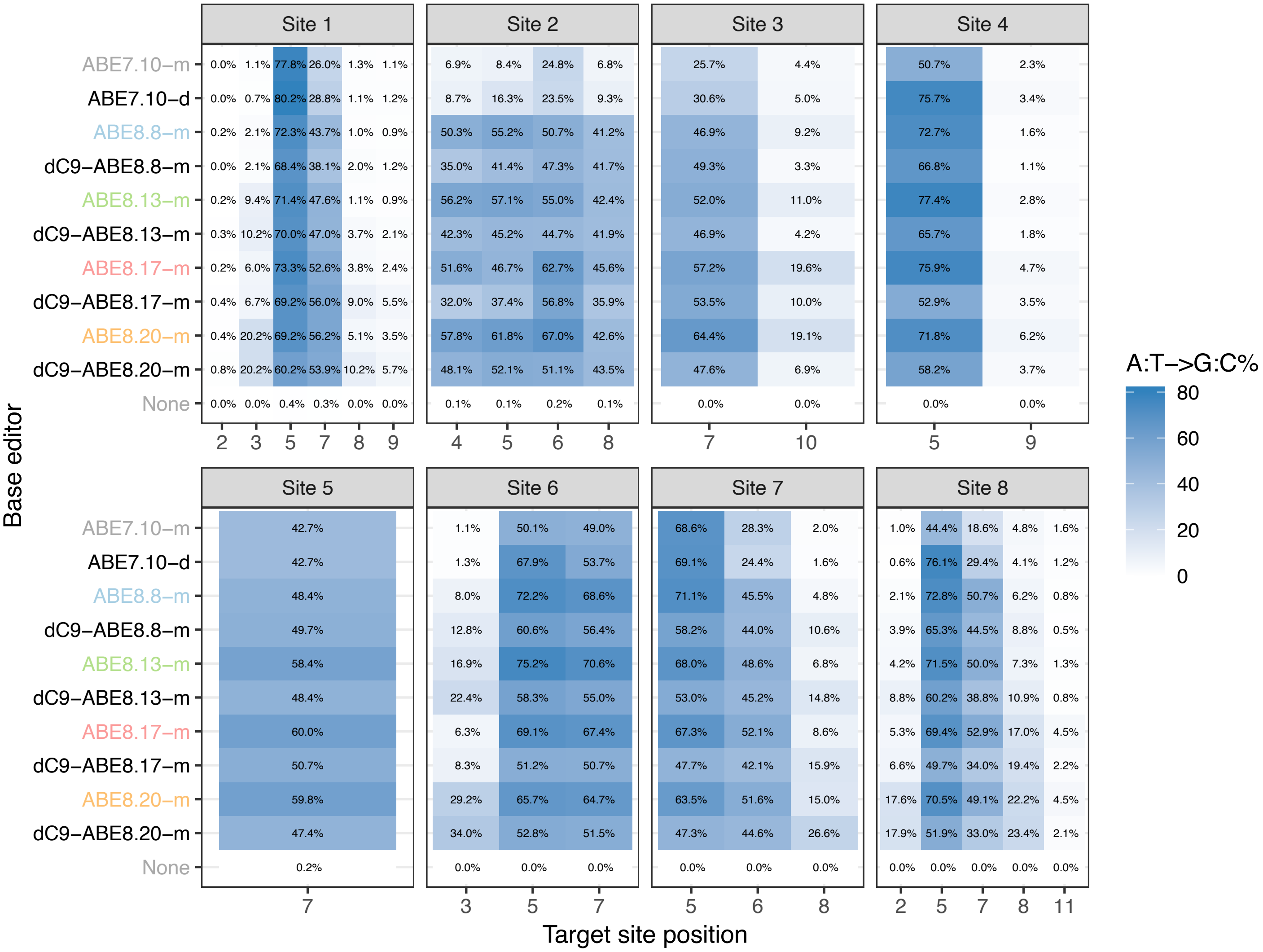


**Supplementary Figure 14 | Median A•T to G•C conversion of core dC9-ABE8-m constructs at eight genomic sites in HEK293T cells.** Dead Cas9 (dC9) is defined as D10A and H840A mutations within *S. pyogenes* Cas9. Median values are shown, generated from n≥3 biological replicates.

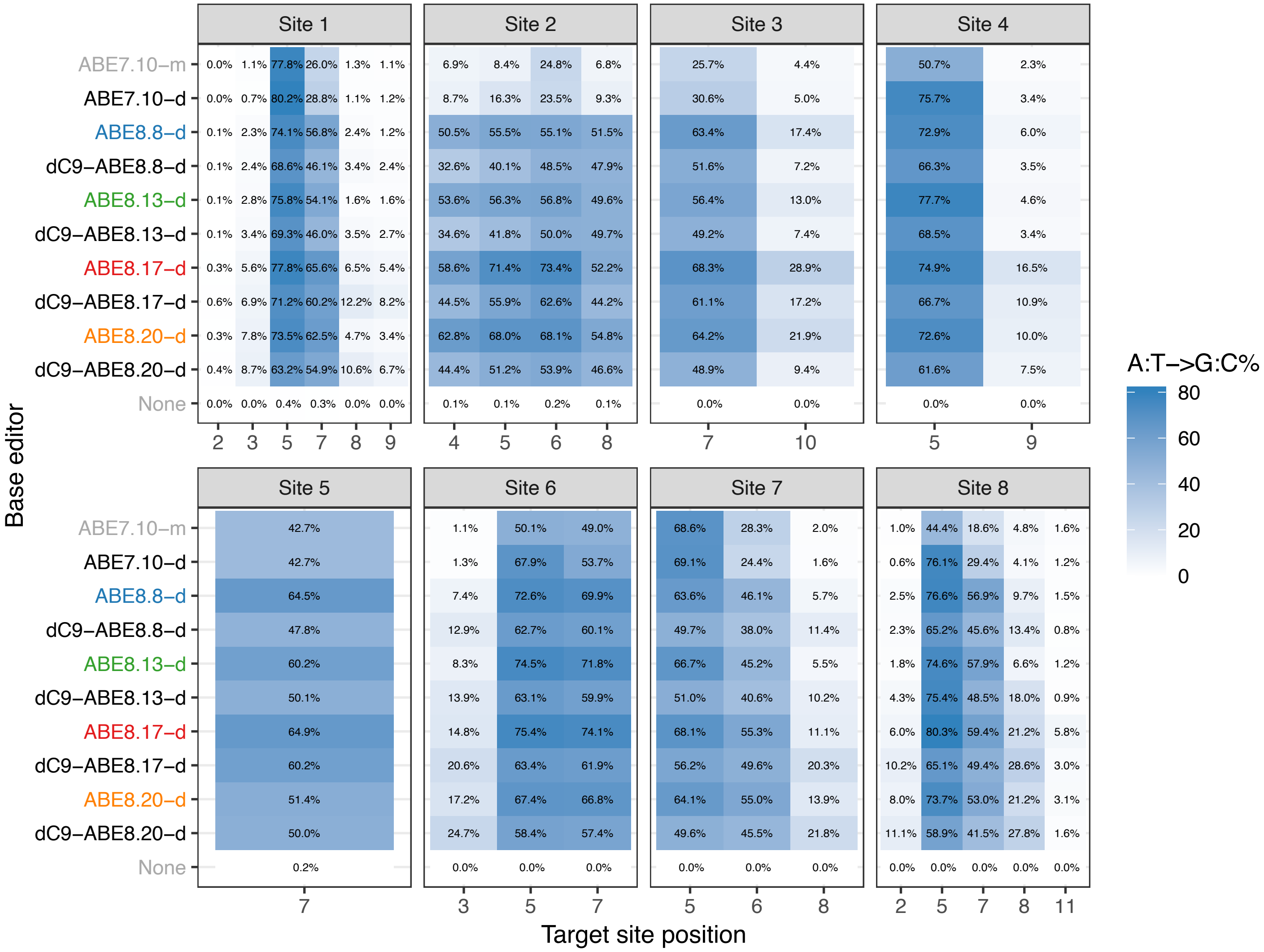


**Supplementary Figure 15 | Median A•T to G•C conversion of core dC9-ABE8-d constructs at eight genomic sites in HEK293T cells.** Dead Cas9 (dC9) is defined as D10A and H840A mutations within *S. pyogenes* Cas9. Median values are shown, generated from n≥3 biological replicates.

**a.**

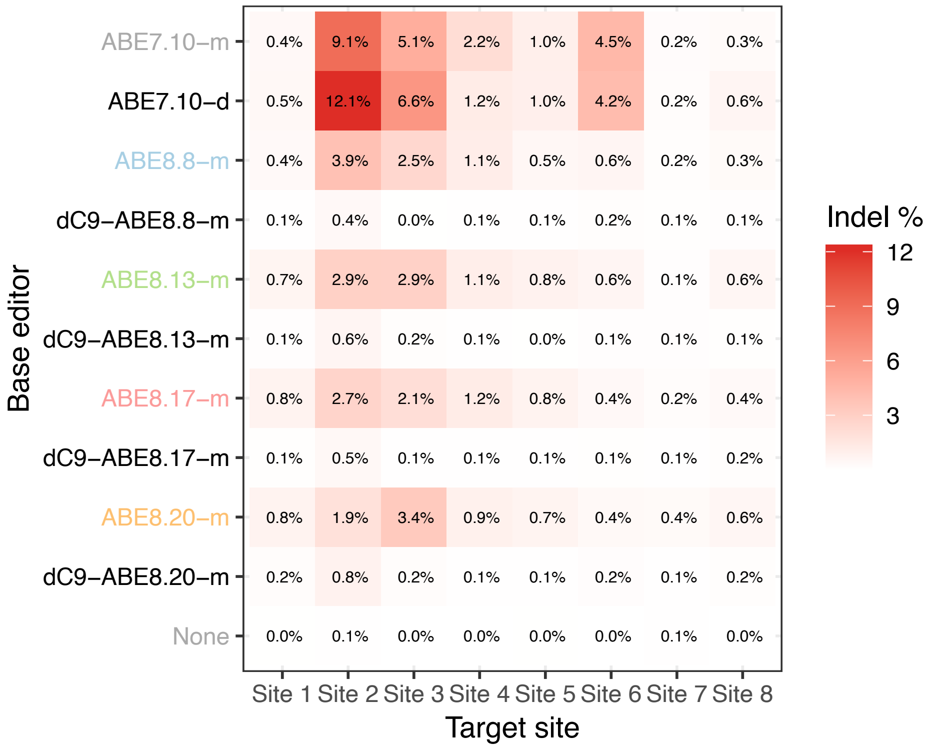


**b.**

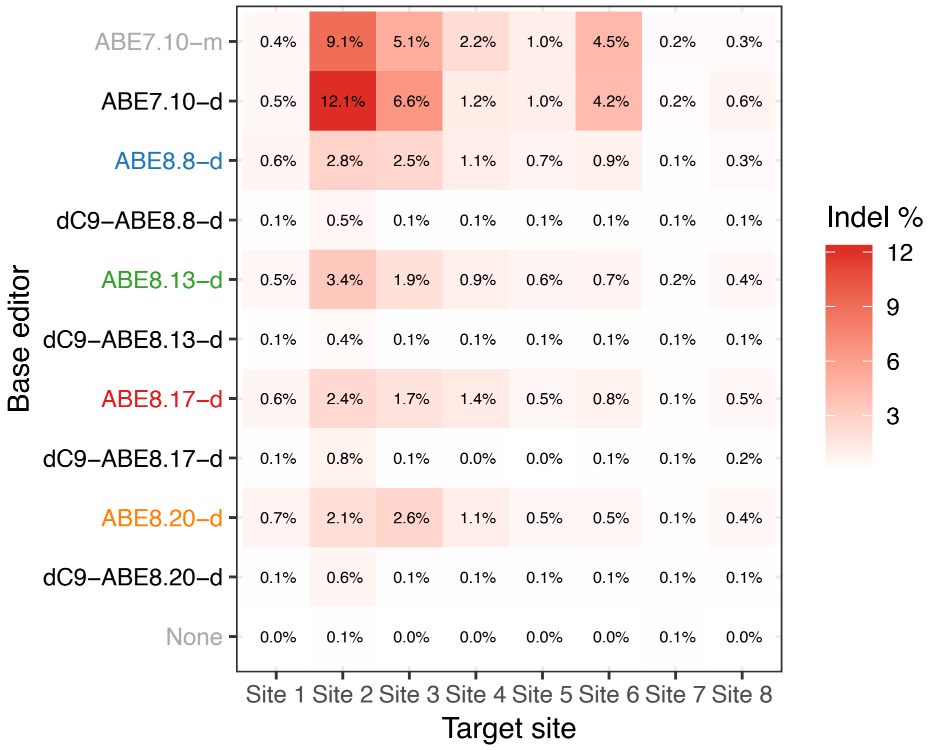


**Supplementary Figure 16 | Median indel frequencies of core dC9-ABE8s tested at 8 genomic sites in HEK293T cells. a,** Indel frequencies are shown for dC9-ABE8-m variants and compared to those from ABE7.10. **b,** Indel frequencies are shown for dC9-ABE8-d variants relative to ABE7.10. Median values are shown, generated from n≥3 biological replicates.

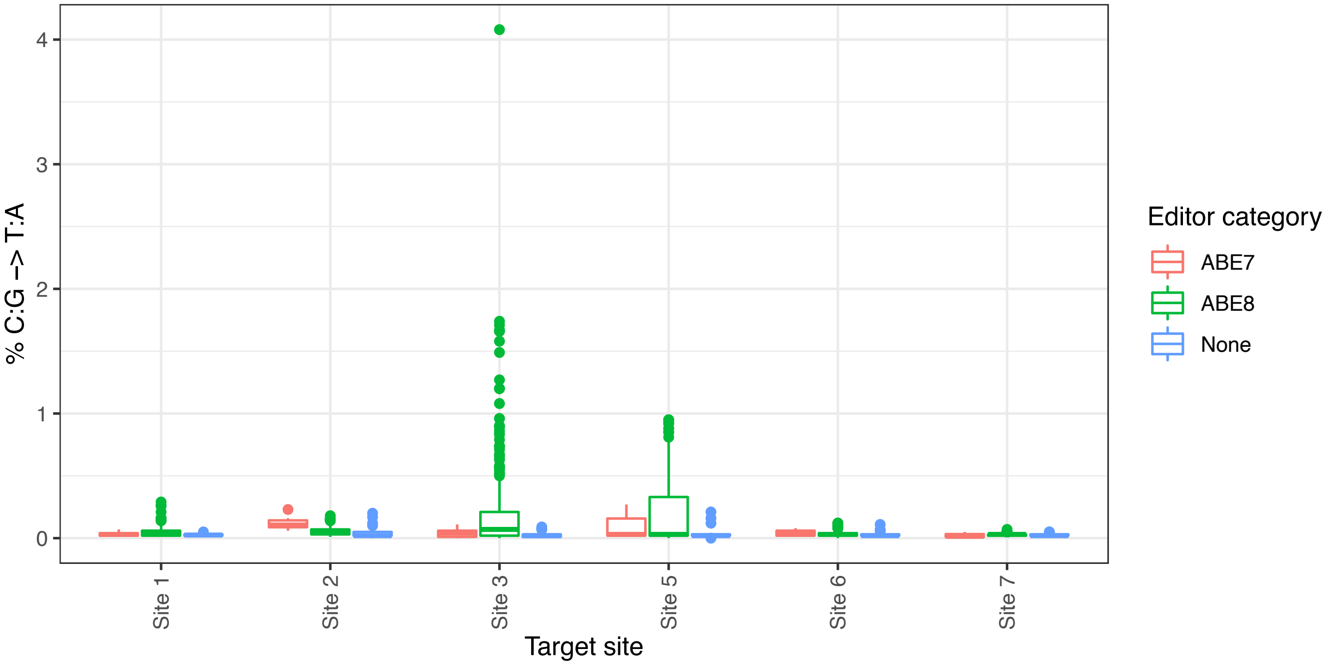


| site | target DNA sequence |
|---|---|
| **1** | **GAACACAAAGCATAGACTGC** |
| **2** | GGGAAAGACCCAGCATCCGT |
| **3** | GCTCCCATCACATCAACCGG |
| **5** | GGCTTCAGGTTCTAAATGAG |
| **6** | GCAGAGAGTCGCCGTCTCCA |
| **7** | GTGTAAGACCTCAAAAGCAC |

**Supplementary Figure 17 | C•G to T•A editing box plot with Hek293T cells treated with ABE8s and ABE7.10.** Editing frequencies for each site averaged across all C's within the target. Cytosines within the protospacer highlighted in blue below. The boundaries of the box indicate the first (bottom) and third (top) quartiles, while the band within the box indicates the median. Values that are farther than 1.5 times the interquartile range (|Q3-Q1|) from the median are marked as outliers and displayed individually. Whiskers extend from the edge of the box to the maximum and minimum values that are not considered outliers. The size of each group shown in the plot is: Site 1: n=384, Site 2: n=354, Site 3: n=732, Site 5: n=366, Site 6: n=366, Site 7: n=366.

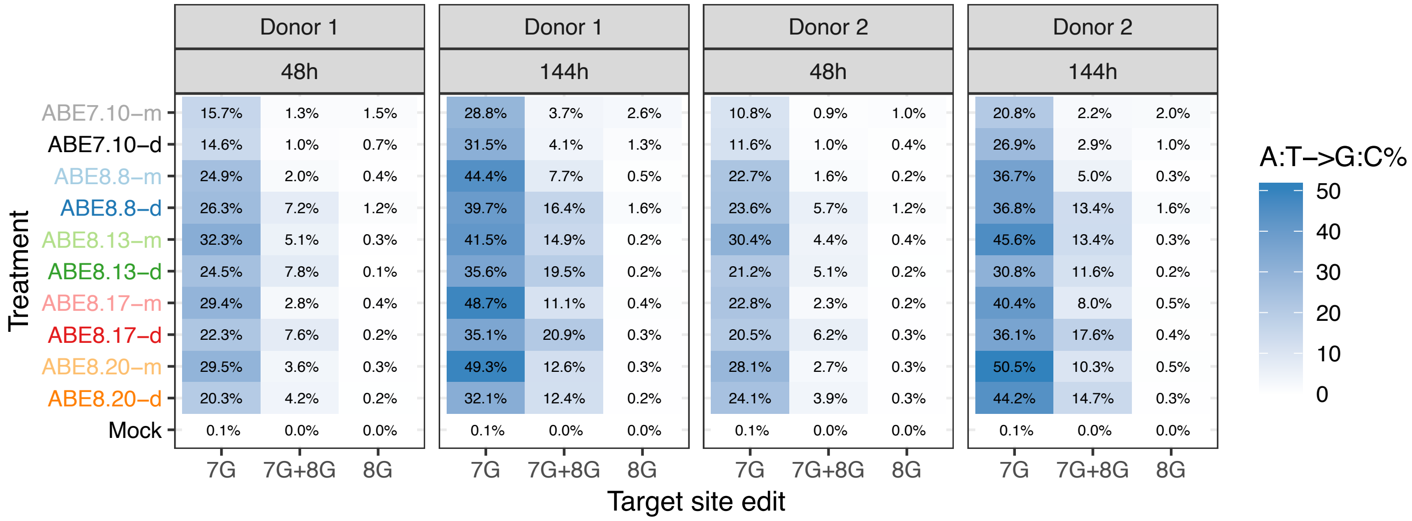


**Supplementary Figure 18 | A•T to G•C conversion of CD34+ cells treated with ABE8 at the -198 promoter site upstream of HBG1/2. a,** Heat map depicting A to G editing frequency of ABE8s in CD34+ cells from two donors at 48 and 144h post editor treatment. Note: Donor 2 is heterozygous for sickle cell disease.

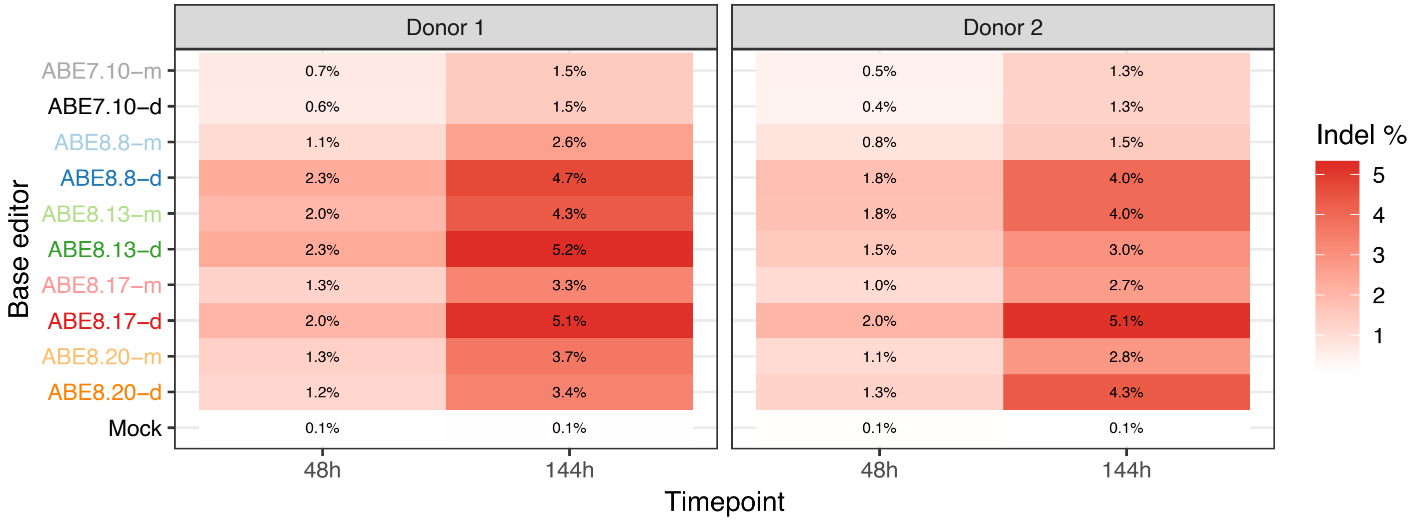


**Supplementary Figure 19 | Indel frequency of CD34+ cells treated with ABE8 at the -198 site of the gamma-globin promoter.** Frequencies shown from two donors at 48h and 144h time points. Note: Complete A•T to G•C conversion at the HBG1/2 -198 promoter target site investigated here, creates a poly-G stretch of 10-nt. Such homopolymer runs often increase the rate of PCR- and sequencing-induced errors, leading to the appearance of elevated indel frequencies at this site.

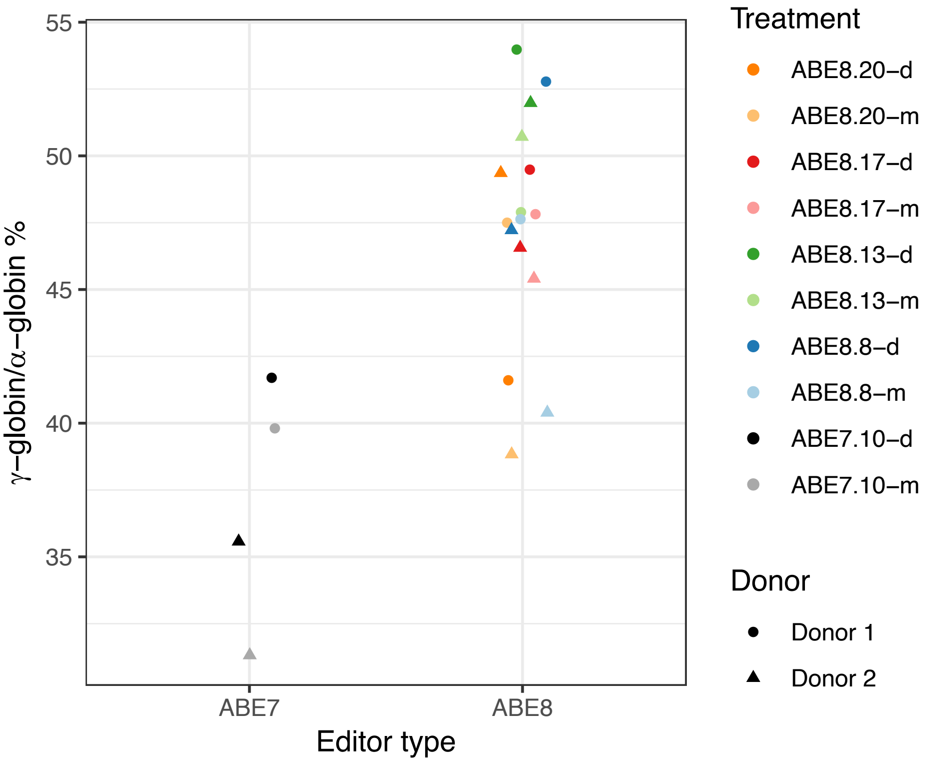


**Supplementary Fig. 20: Median gamma-globin levels across all ABE8-treated vs. ABE7-treated samples.** Each point represents the measured gamma-globin over alpha-globin percentage as measured by UPLC for a given base editor (color) and a given CD34+ cell donor (shape).

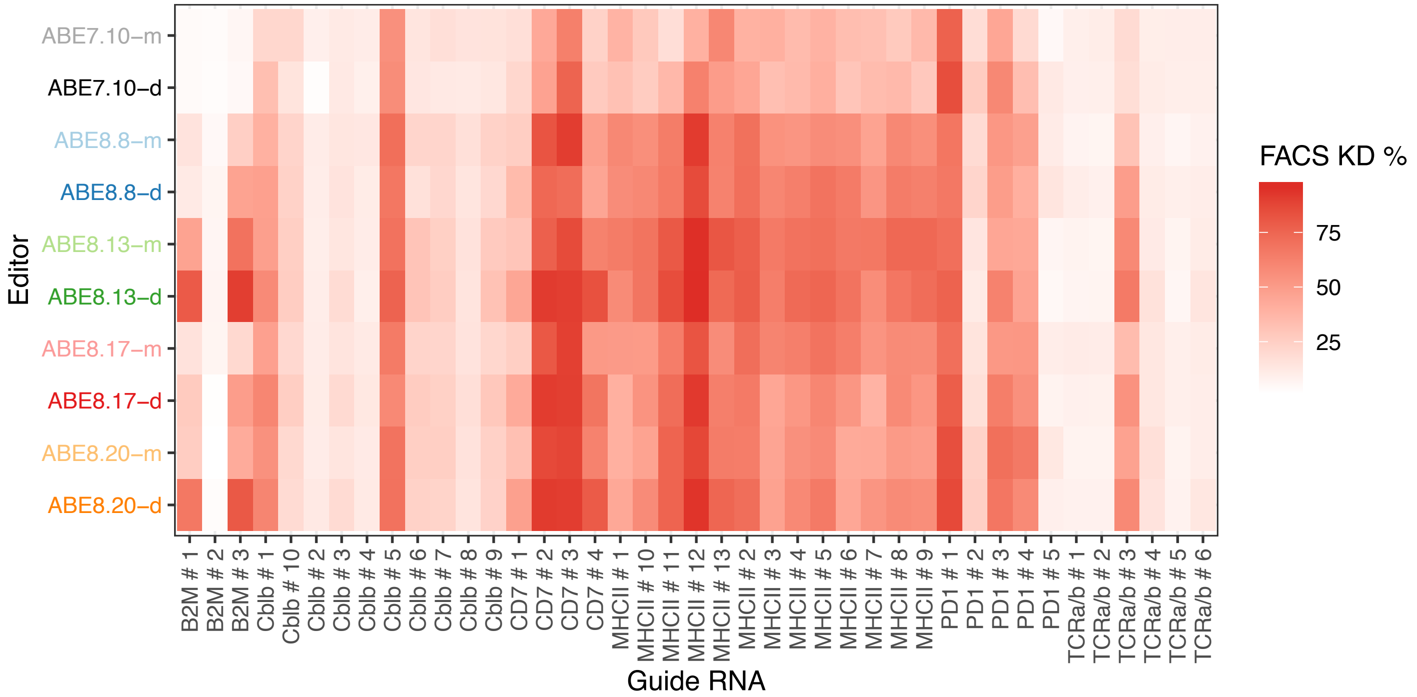


**Supplementary Figure 21 | Protein knockdown measured by flow cytometry by ABE editors in primary T cells.** Eight mRNAs encoding ABE8 editors and two mRNAs encoding ABE7.10-m/d were individually transfected into T cells with 41 sgRNAs targeting six genes and their effects on protein expression were measured using flow cytometry. Values shown are the mean of n=2 independent replicates.

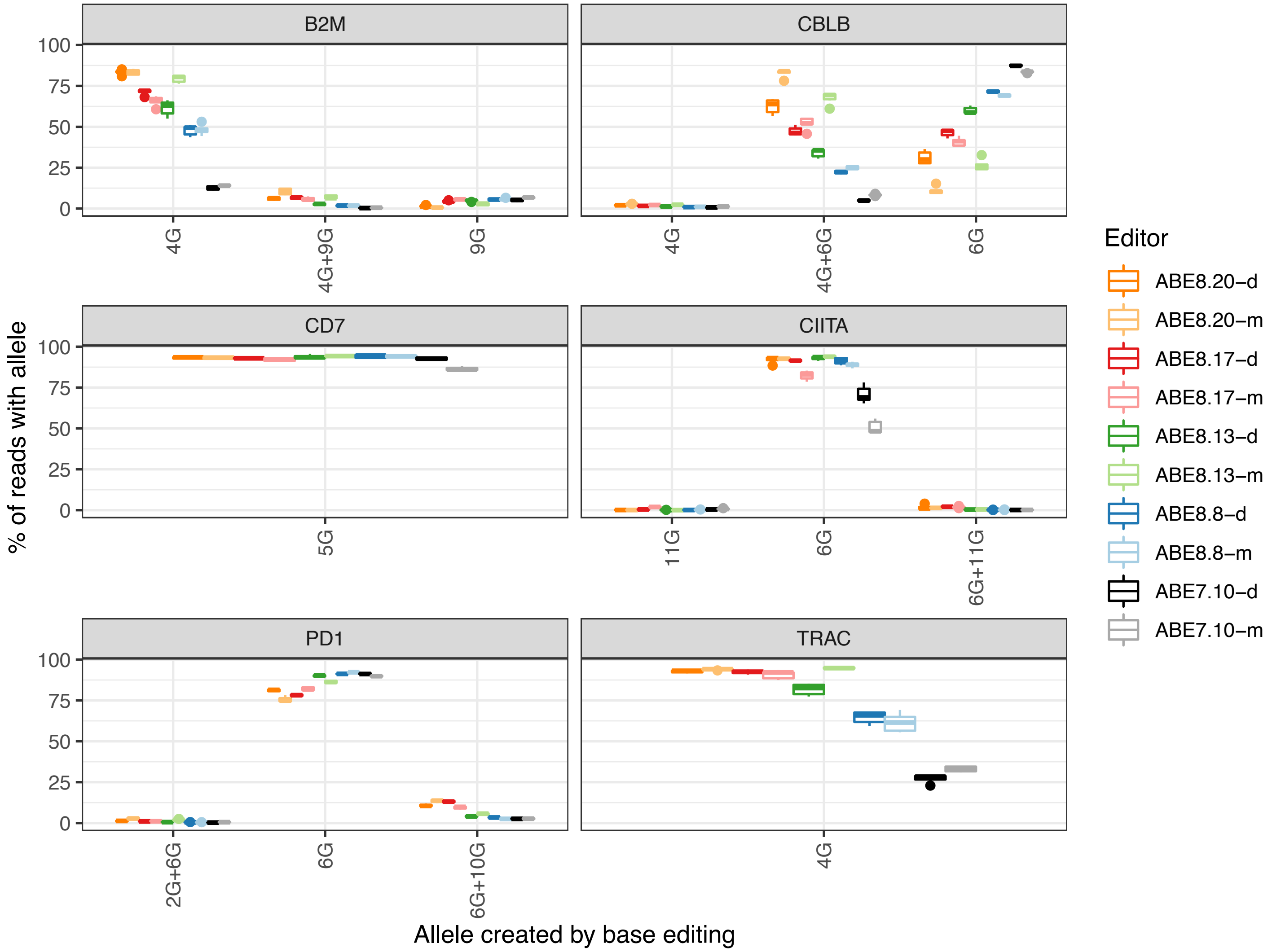


**Supplementary Figure 22: Variant alleles created by ABEs at six different targets in primary T cells to prevent the expression of single genes relevant to creation of T-cell therapies.** Each panel shows the percentage of mapped sequencing reads (y-axis) containing a particular combination of base substitution(s) for a given target site (x-axis). Each group shown as a box consists of 5 observations. The color of each bar corresponds to the base editor used. Only alleles that occurred with a frequency of >1% in at least one sample were included. The boundaries of the box indicate the first (bottom) and third (top) quartiles, while the band within the box indicates the median. Values that are farther than 1.5 times the interquartile range (|Q3-Q1|) from the median are marked as outliers and displayed individually. Whiskers extend from the edge of the box to the maximum and minimum values that are not considered outliers.

| | HEK2 OT1 | HEK2 OT2 | HEK3 OT1t | HEK3 OT2 | HEK3 OT3 | HEK3 OT4 | HEK3 OT5 | HEK4 OT1 | HEK4 OT2 | HEK4 OT3 | HEK4 OT4 | HEK4 OT5 |
|---|---|---|---|---|---|---|---|---|---|---|---|---|
| ABE7.10-d | 0.1 | $1.7\times10^{-2}$ | $1.4\times10^{-2}$ | $1.6\times10^{-2}$ | $1.4\times10^{-2}$ | $1.6\times10^{-2}$ | $1.5\times10^{-2}$ | 0.1 | $3.9\times10^{-2}$ | 17.6 | 0.1 | $1.5\times10^{-2}$ |
| ABE7.10-m | 0.1 | $2.2\times10^{-2}$ | $3.0\times10^{-2}$ | $2.0\times10^{-2}$ | $1.6\times10^{-2}$ | $1.5\times10^{-2}$ | $2.0\times10^{-2}$ | 0.1 | 0.1 | 20.9 | 0.1 | $2.1\times10^{-2}$ |
| ABE8.8-d | $3.9\times10^{-2}$ | $2.4\times10^{-2}$ | $1.0\times10^{-2}$ | $1.9\times10^{-2}$ | $1.4\times10^{-2}$ | $2.0\times10^{-2}$ | $2.4\times10^{-2}$ | 0.2 | 0.1 | 5.6 | 0.3 | $4.9\times10^{-2}$ |
| ABE8.8-m | $4.9\times10^{-2}$ | $2.0\times10^{-2}$ | $1.6\times10^{-2}$ | $2.4\times10^{-2}$ | $1.5\times10^{-2}$ | $2.6\times10^{-2}$ | $1.7\times10^{-2}$ | 0.2 | 0.2 | 6.2 | 0.3 | $8.3\times10^{-3}$ |
| ABE8.13-d | 0.1 | $2.0\times10^{-2}$ | $2.2\times10^{-2}$ | $1.8\times10^{-2}$ | $1.6\times10^{-2}$ | $1.8\times10^{-2}$ | $3.6\times10^{-2}$ | 0.2 | 0.1 | 6.3 | 0.3 | $1.6\times10^{-2}$ |
| ABE8.13-m | $2.4\times10^{-2}$ | $1.6\times10^{-2}$ | $1.0\times10^{-2}$ | $2.2\times10^{-2}$ | $9.7\times10^{-3}$ | $1.8\times10^{-2}$ | $1.8\times10^{-2}$ | 0.3 | 0.1 | 12.5 | 0.5 | $2.6\times10^{-2}$ |
| ABE8.17-d | $4.3\times10^{-2}$ | $1.9\times10^{-2}$ | $1.0\times10^{-2}$ | $1.6\times10^{-2}$ | $1.8\times10^{-2}$ | $2.0\times10^{-2}$ | $9.0\times10^{-3}$ | 0.1 | 0.1 | 9.9 | 0.3 | 0.1 |
| ABE8.17-m | $1.1\times10^{-2}$ | $2.2\times10^{-2}$ | $1.1\times10^{-2}$ | $3.4\times10^{-2}$ | $1.7\times10^{-2}$ | $1.9\times10^{-2}$ | $2.4\times10^{-2}$ | 0.2 | 0.1 | 9.3 | 0.4 | $2.2\times10^{-2}$ |
| ABE8.20-d | $4.2\times10^{-2}$ | $2.0\times10^{-2}$ | $1.3\times10^{-2}$ | $1.5\times10^{-2}$ | $1.6\times10^{-2}$ | $2.0\times10^{-2}$ | $3.2\times10^{-2}$ | 0.2 | 0.2 | 7.1 | 0.5 | $2.5\times10^{-2}$ |
| ABE8.20-m | 0.1 | $2.1\times10^{-2}$ | $1.3\times10^{-2}$ | $1.7\times10^{-2}$ | $1.2\times10^{-2}$ | $1.8\times10^{-2}$ | $1.7\times10^{-2}$ | 0.3 | 0.3 | 6.5 | 0.7 | $4.5\times10^{-2}$ |
| Cas9 nickase control | $4.3\times10^{-2}$ | $2.2\times10^{-2}$ | $8.0\times10^{-3}$ | $1.3\times10^{-2}$ | $2.3\times10^{-2}$ | $1.8\times10^{-2}$ | $1.7\times10^{-2}$ | 0.2 | 0.1 | 5.3 | 0.1 | $3.6\times10^{-2}$ |
| Cas9 nuclease | 8.4 | 0.2 | 17.1 | 22.0 | 2.6 | 0.3 | 0.4 | 28.6 | 21.6 | 95.0 | 18.0 | 35.2 |

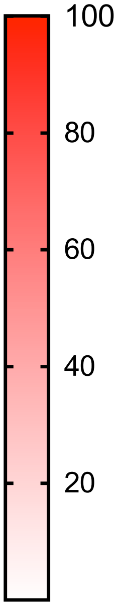


**Supplementary Figure 23 | Median indel frequencies at 12 previously identified sgRNA-dependent Cas9 off-target loci in human cells** Data shown is the median value from n=3 independent biological replicates, performed on different days. Constructs were administered to HEK293T cells using plasmid delivery.

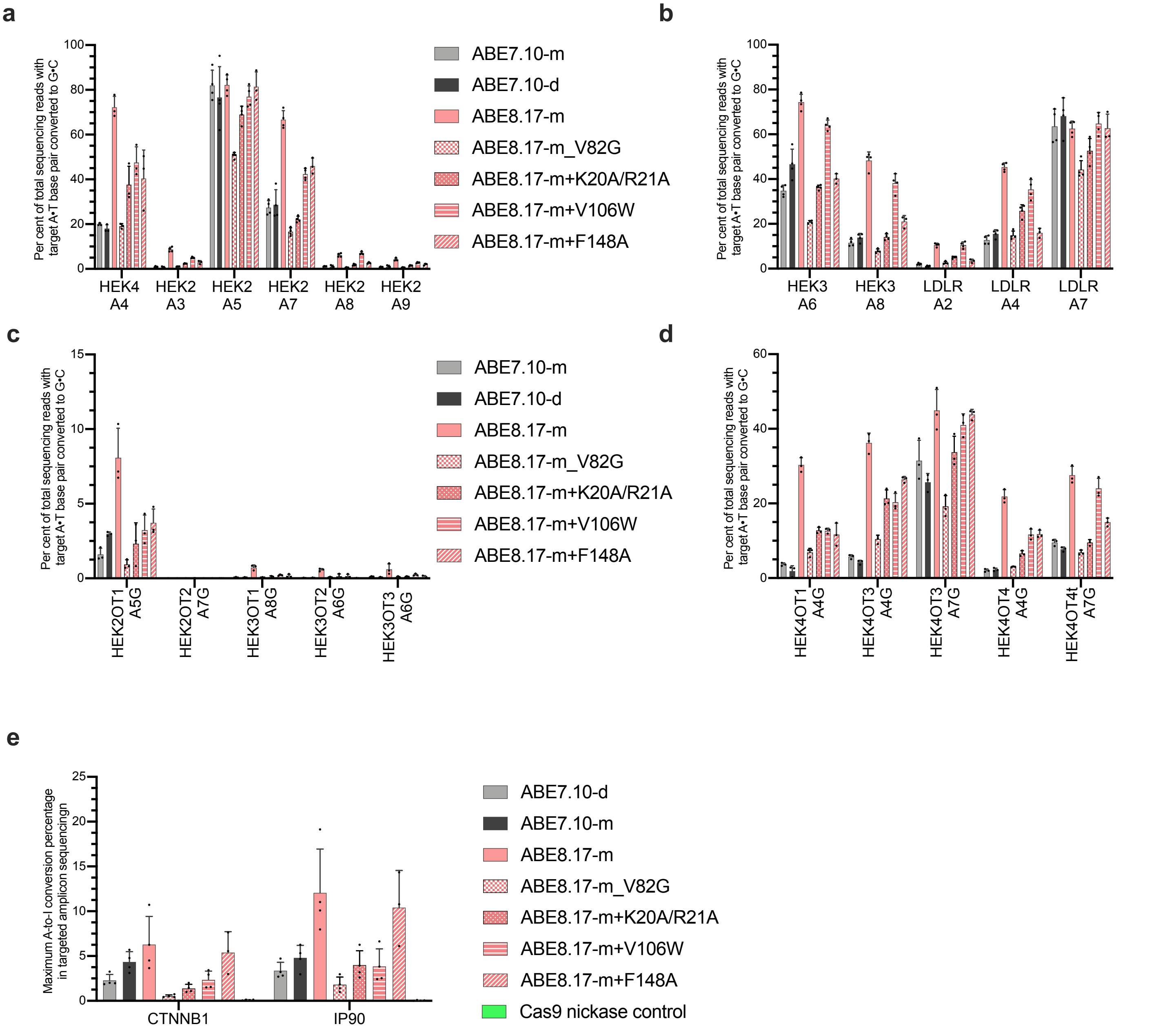


**Supplementary Figure 24 | DNA on-target, sgRNA-dependent DNA off-target editing and sgRNA-independent off-target mRNA editing by ABE8 constructs containing mutations designed to reduced spurious cellular RNA editing.** Constructs were administered to HEK293T cells using plasmid delivery. **a** and **b** show on-target editing frequencies at four genomic loci, **c** and **d** show off-target editing frequencies at known, sgRNA-dependent off-target editing loci and **e** shows the maximum level of A-to-I mutation in cellular mRNA samples isolated from cells treated with the indicated construct. Individual data points are shown and error bars represent s.d. for n=3 or n=4 independent biological replicates, as can be determined for each sample by the number of included data points on the graph. The bar for each condition represents the mean value.

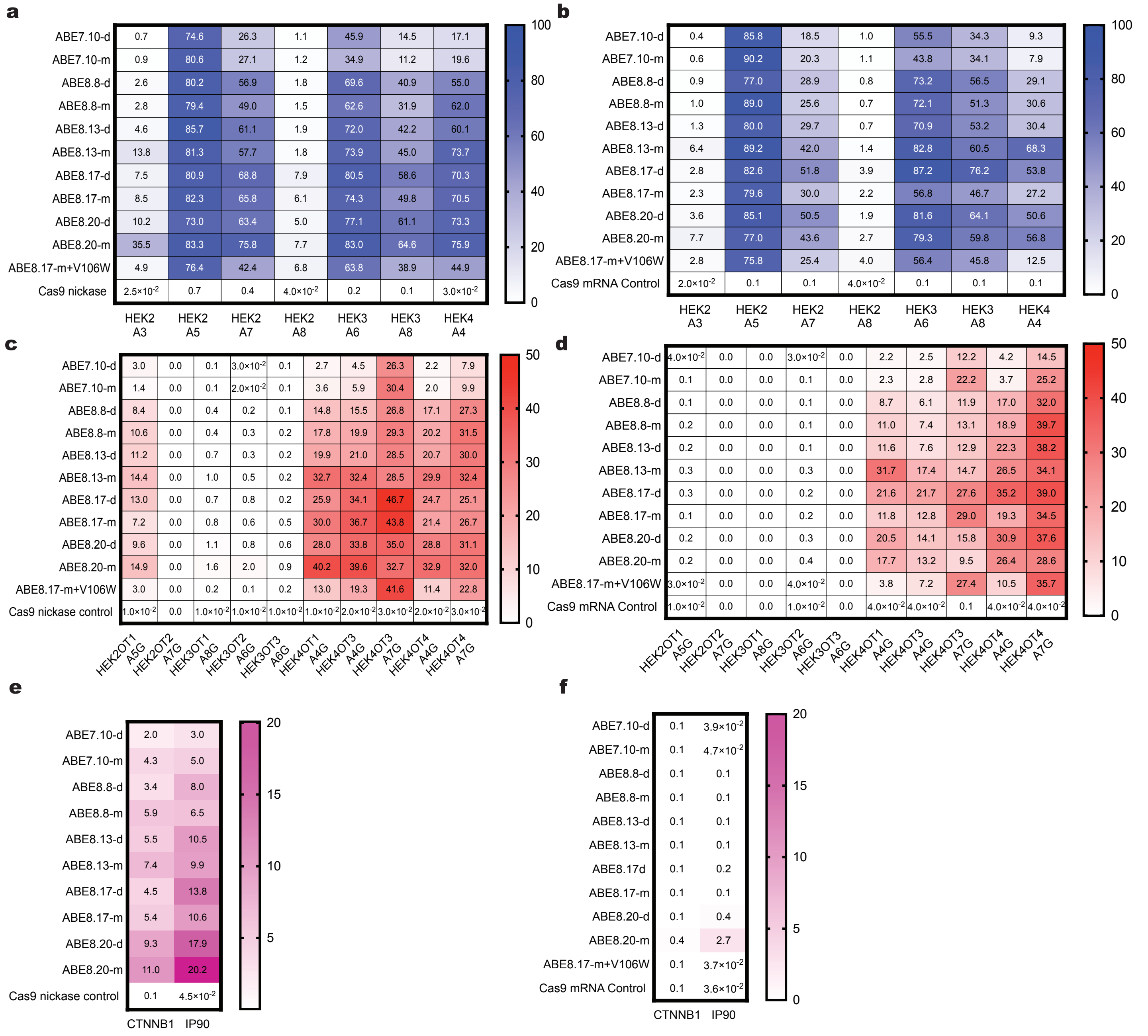


**Supplementary Figure 25 | DNA on-target base editing, sgRNA-dependent DNA off-target base editing and sgRNA-independent off-target mRNA editing by ABE8 constructs using plasmid or mRNA delivery a** and **b**: on-target DNA editing frequencies for core ABE 8 constructs as compared to ABE7. Constructs were delivered as plasmid in **a** and as mRNA in **b**, **c** and **d**: sgRNA-dependent off-target DNA editing frequencies for ABE8 as compared to ABE7. Constructs were delivered as plasmid in **c** and as mRNA in **d. e, f:** The maximum A-to-G editing frequency measured in a 125-nt region of the indicated amplicon**.** Constructs were delivered as plasmid in **e** and as mRNA in **f.** The median is shown for n=3 or n=4 independent biological replicates, performed on different days, other than in **d**, where some samples yielded <5,000 Miseq reads so were excluded; specifically HEK4OT3 (ABE7.10-d n=1 and ABE8.13-m n=2) and HEK4OT4 (ABE8.13-d, ABE8.17-d, ABE8.20-m and ABE8.20-d n=2) and in **f** which ABE8.8-m: n=2 for CTNNB1.

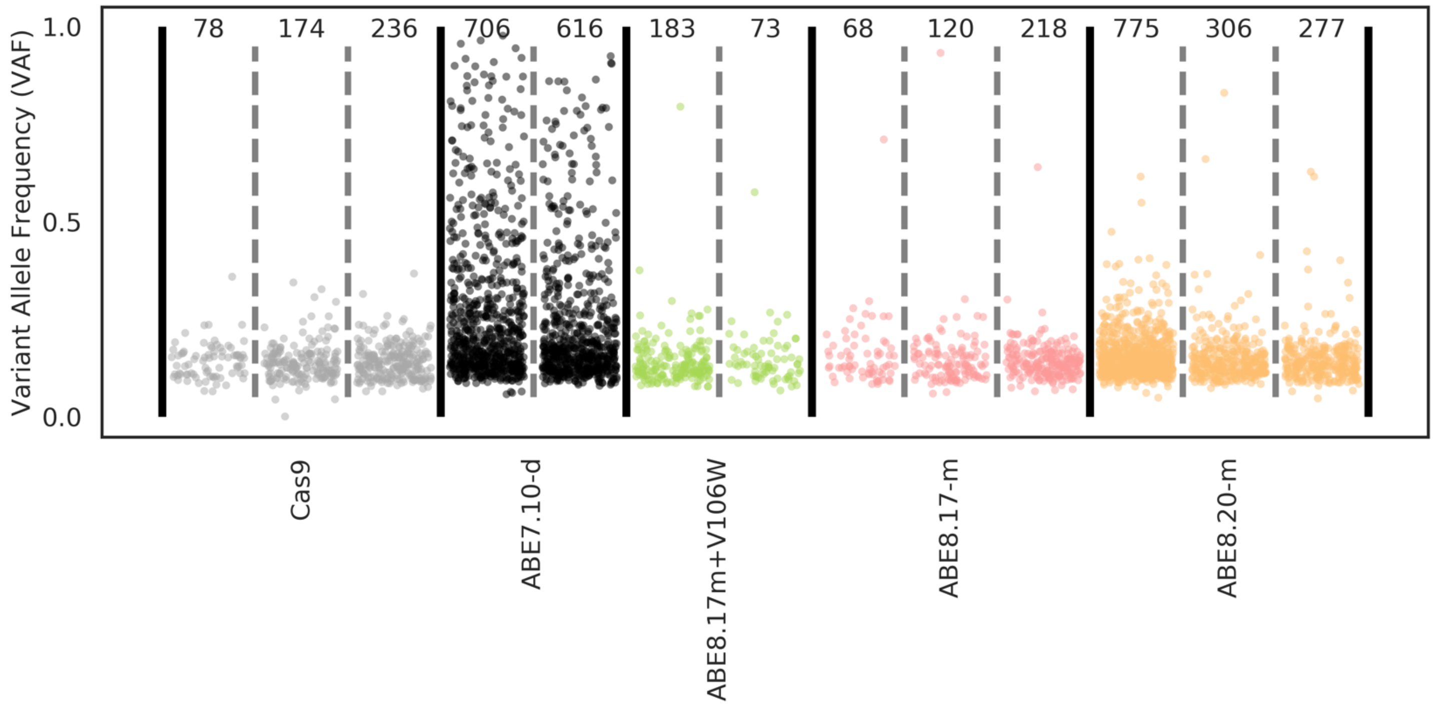


**Supplementary Figure 26 | Whole transcriptome sequencing in HEK293T cells treated with indicated mRNA.** Strip plot representing the variant allele frequency of transcriptome wide A->G mutations in RNA observed in replicate HEK293T cell experiments. Total A->G mutations are indicated at the top of the panel for each sample. These numbers correspond to the number of observations in each group.

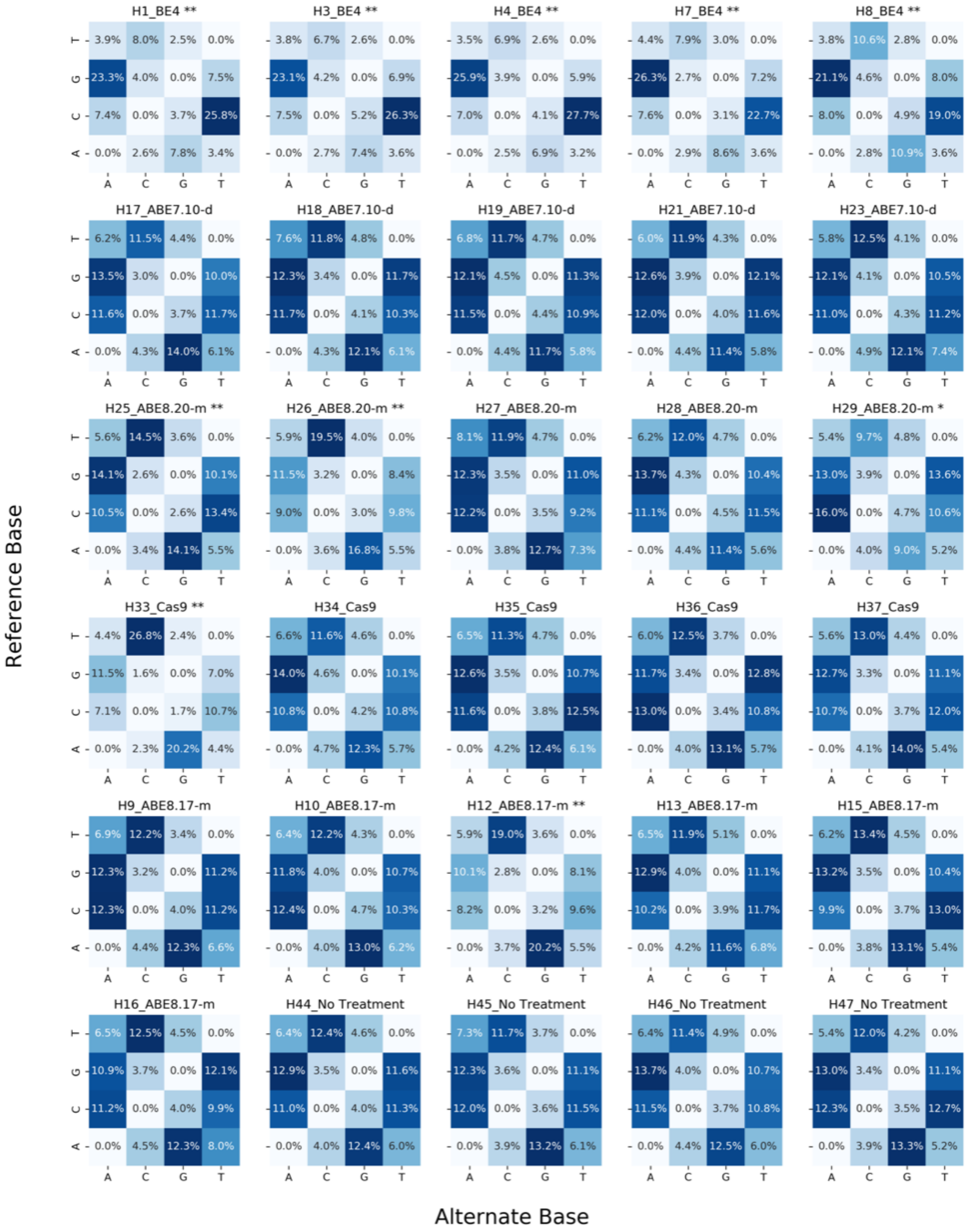


**Supplementary Figure 27 | Mutational classification plots for each sample sent for whole genome sequencing.** Distribution of mutation types in all whole genome sequenced samples. Samples significantly enriched for mutations of the type the base editor creates indicated with **. Sample with significantly less mutations of the type the editor creates indicated with *

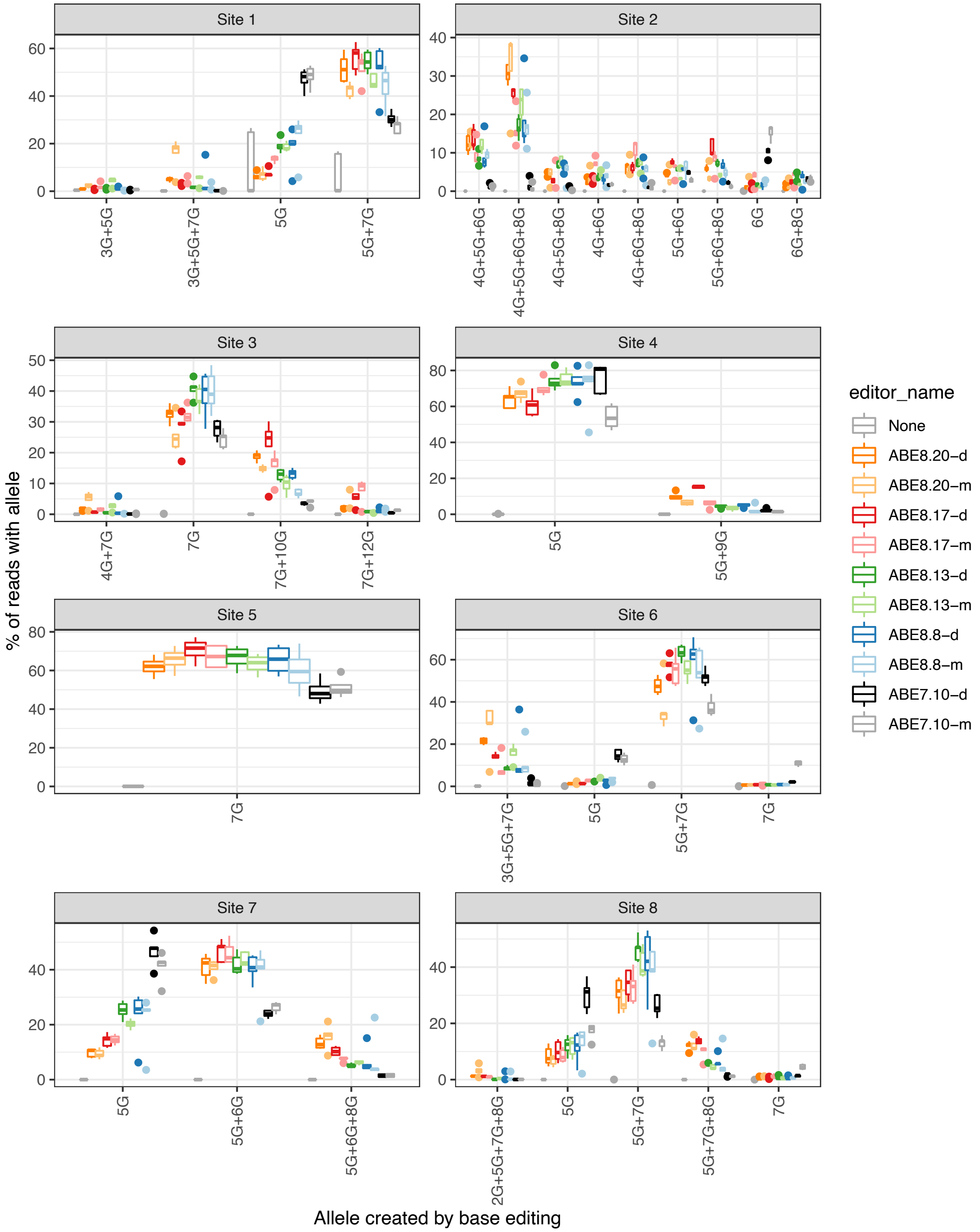


**Supplementary Figure 28 | Alleles created by ABEs across 8 different genomic sites in HEK293T cells.** Each panel shows the percentage of mapped sequencing reads (y-axis) containing a particular combination of base substitution(s) for a given target site (x-axis). The color of each bar corresponds to the base editor used. Only alleles that occurred with a frequency of >5% in at least one sample were included. Each group shown as a box includes 5 observations. The boundaries of the box indicate the first (bottom) and third (top) quartiles, while the band within the box indicates the median. Values that are farther than 1.5 times the interquartile range (|Q3-Q1|) from the median are marked as outliers and displayed individually. Whiskers extend from the edge of the box to the maximum and minimum values that are not considered outliers.

**a**

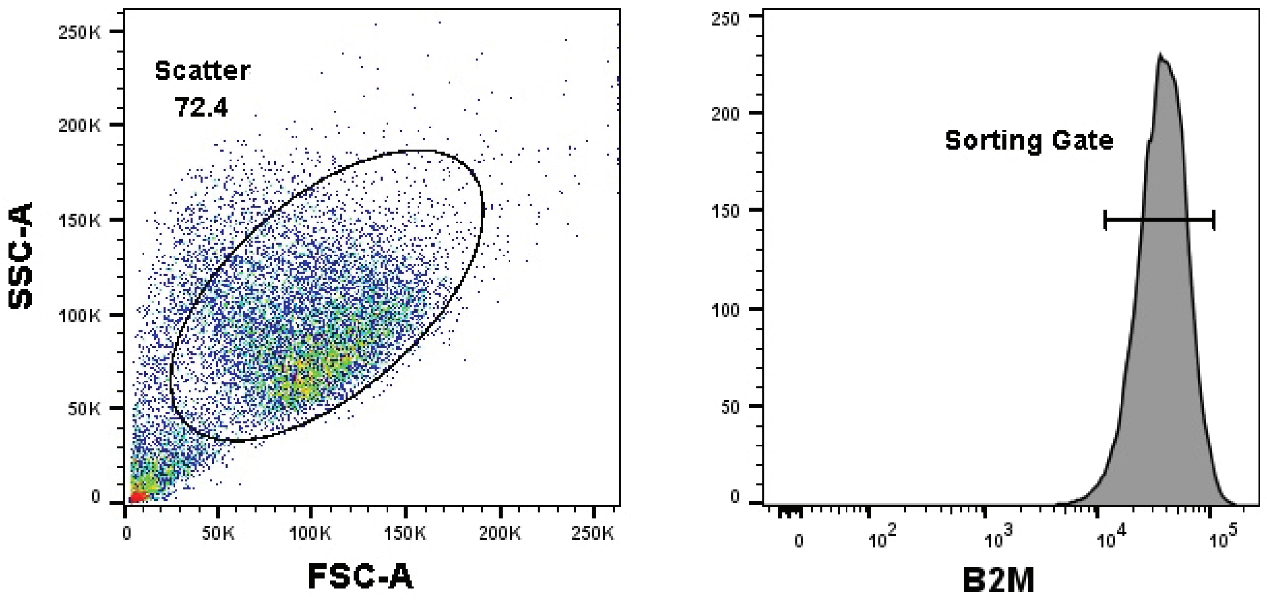


**b**

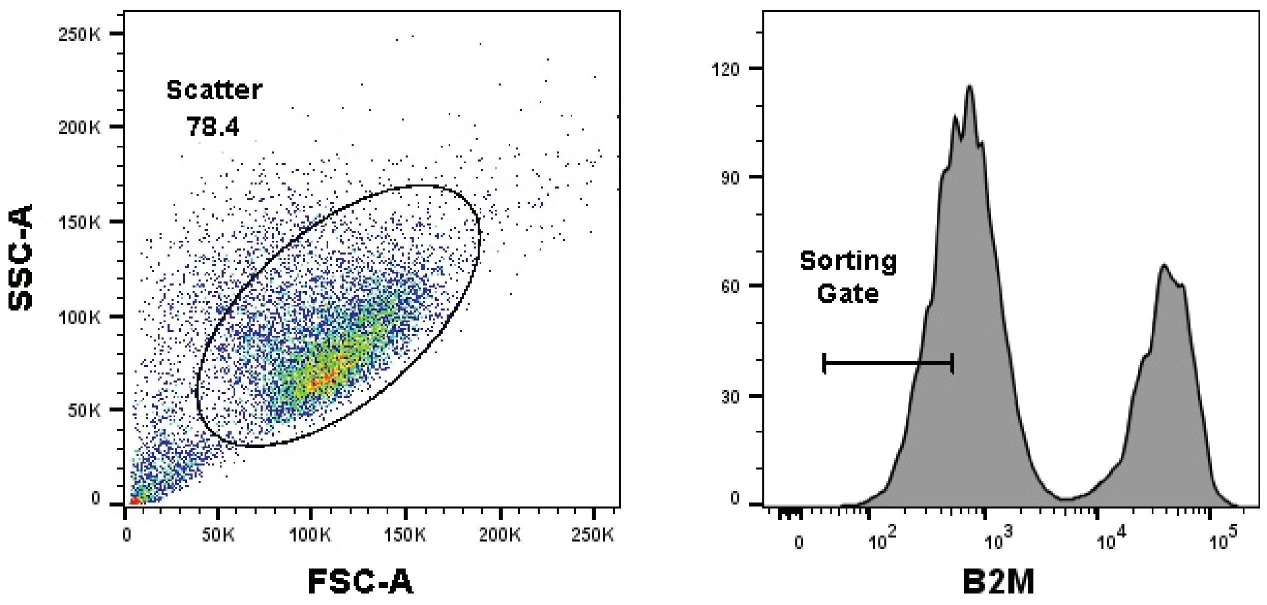


**Supplementary Figure 29 | Representative examples of gates used to flow sort B2M-positive and B2M-negative cells prior to whole genome sequencing. a:** representative plot and gate for live, B2M-positive HEK293T cells sorted into single cell clones for the untreated condition. **b**: representative plot and gate for live, B2M-negative HEK293T cells sorted for the all treated conditions (ABE, CBE or Cas9-treated cells)

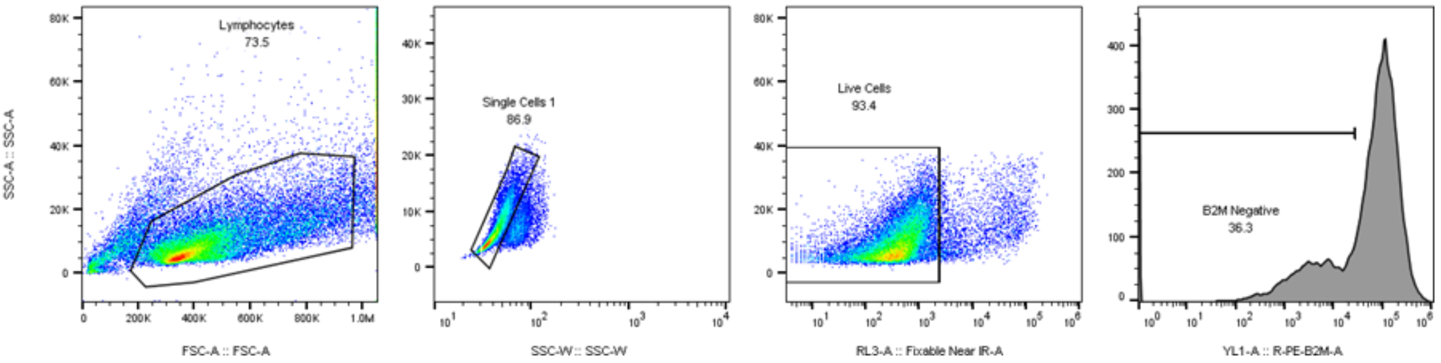


**Supplementary Figure 30 | Examples of gates used for assessment of protein knockdown in T cells.** Representative gating strategy for population analysis on live, single, lymphocytes in order to determine surface protein reduction via flow cytometry.

**Supplementary Table 1:** Sequences of sgRNAs used for HEK293T mammalian cell transfection. The 20-nt target protospacer is shown in red. When a target DNA sequence did not start with a G a G was added to the 5' end of the primer since it has been established that the human U6 promoter prefers a G at the transcription start site[1]. The pFYF sgRNA plasmid described previously was used as a template for PCR amplification[2].

| Site | RNA protospacer sequence | Cas9 scaffold | PAM |
|---|---|---|---|
| **1** | GAACACAAAGCAUAGACUGC | *S. pyogenes* | NGG |
| **2** | GGGAAAGACCCAGCAUCCGU | *S. pyogenes* | NGG |
| **3** | GCUCCCAUCACAUCAACCGG | *S. pyogenes* | NGG |
| **4** | GGUGAGUGAGUGUGUGCGUG | *S. pyogenes* | NGG |
| **5** | GGCUUCAGGUUCUAAAUGAG | *S. pyogenes* | NGG |
| **6** | GCAGAGAGUCGCCGUCUCCA | *S. pyogenes* | NGG |
| **7** | GUGUAAGACCUCAAAAGCAC | *S. pyogenes* | NGG |
| **8** | GAUGAGAAGGAGAAGUUCUU | *S. pyogenes* | NGG |
| **9** | GAGGACAAAGUACAAACGGC | *S. pyogenes* | AGA |
| **10** | GCCACCACAGGGAAGCUGGG | *S. pyogenes* | TGA |
| **11** | GCUCUCAGGCCCUGUCCGCA | *S. pyogenes* | CGT |
| **12** | GAGCAAAUACCAGAGAUAAG | *S. pyogenes* | AGA |
| **13** | GAUCAGGAAAUAGAGCCACA | *S. pyogenes* | GGC |
| **14** | GCCCAUCCCUGAGUCCAGCG | *S. pyogenes* | AGC |
| **15** | GAACACGAAGACAUCUGAAGGUA | *S. aureus* | TTGAAT |
| **16** | GAUUUACAGCCUGGCCUUUGGGG | *S. aureus* | TCGGGT |
| **17** | GGAGAGAAAGAGAAGUUGAUUG | *S. aureus* | ATGGGT |
| **18** | GAGGGUGAGGGAUGAGAUAAUG | *S. aureus* | ATGAGT |
| **19** | GGUGGAGGAGGGUGCAUGGGGU | *S. aureus* | CAGAAT |
| **20** | GCUGUUGCAUGAGGAAAGGGAC | *S. aureus* | TAGAGT |
| **HEK2** | GAACACAAAGCAUAGACUGC | *S. pyogenes* | CGG |
| **HEK3** | GGCCCAGACUGAGCACGUGA | *S. pyogenes* | TGG |
| **HEK4** | GGCACUGCGGCUGGAGGUGG | *S. pyogenes* | GGG |
| **LDLR** | GCAGAGCACUGGAAUUCGUCA | *S. pyogenes* | GGG |

**sgRNA scaffold sequences:**

***S. pyogenes:***
GUUUUAGAGCUAGAAAUAGCAAGUUAAAAUAAGGCUAGUCCGUUAUCAACUUGAAAAAGUGGCACCGAGUCGGUGC

***S. aureus:***
GUUUUAGUACUCUGUAAUGAAAAUUACAGAAUCUACUAAAACAAGGCAAAAUGCCGUGUUUAUCUCGUCAACUUGUUGGCGAGA

**Supplementary Table 2:** Sequences of sgRNAs used for CD34+ and T cell transfections

| Site | RNA protospacer sequence | Cas9 scaffold | supplier |
|---|---|---|---|
| **21** | csususACCCCACUUAACUAUCU | *S. pyogenes* | Synthego |
| **22** | cscscsUACCUGUCACCAGGACC | *S. pyogenes* | Synthego |
| **23** | csascsCUACCUAAGAACCAUCC | *S. pyogenes* | Synthego |
| **24** | csascsUCACCUUAGCCUGAGCA | *S. pyogenes* | Synthego |
| **25** | csususACCUGGGCUGGGGAAGA | *S. pyogenes* | Synthego |
| **26** | asususAUACCUGCCAUGCCGUA | *S. pyogenes* | Synthego |
| **-198 HBG1/2** | gsusgsGGGAAGGGGCCCCCAAG | *S. pyogenes* | Biospring |

a,c,g,u: 2'-O-methyl residues
s: phosphorothioate

**sgRNA scaffold sequences:**
***S. pyogenes:***
5'GUUUUAGAGCUAGAAAUAGCAAGUUAAAAUAAGGCUAGUCCGUUAUCAACUUGAAAAAGUGGCACCGAGUC
GGUGCUsususu-3'

**Supplementary Table 3:** Primers used for ABE8 T7 in vitro transcription reactions

| name | sequence |
|---|---|
| **fwd_IVT** | TCGAGCTCGGTACCTAATACGACTCAC |
| **rev_IVT** | TTTTTTTTTTTTTTTTTTTTTTTTTTTTTTTTTTTTTTTTTTTTTTTTTTTTTTTTTTTTTTTTT<br>TTTTTTTTTTTTTTTTTTTTTTTTTTTTTTTTTTTTTTTTTTTTTTTTTTTTTTTTTTTTTTTTT<br>CTTCCTACTCAGGCTTTATTCAAAGACCA |

**Supplementary Table 4**: HTS Primers used to amplify genomic sites:

| primer name | sequence |
|---|---|
| fwd_site_1 | ACACTCTTTCCCTACACGACGCTCTTCCGATCTNNNNCCAGCCCCATCTGTCAAACT |
| rev_site_1 | TGGAGTTCAGACGTGTGCTCTTCCGATCTTGAATGGATTCCTTGGAAACAATGA |
| fwd_site_2 | ACACTCTTTCCCTACACGACGCTCTTCCGATCTNNNNTGAGGGAGAGCCGTGTAGTT |
| rev_site_2 | TGGAGTTCAGACGTGTGCTCTTCCGATCTGCCTCTCAAAGTGCTGGGAT |
| fwd_site_3 | ACACTCTTTCCCTACACGACGCTCTTCCGATCTNNNNCCATCAGGCTCTCAGCTCAG |
| rev_site_3 | TGGAGTTCAGACGTGTGCTCTTCCGATCTCTCGTGGGTTTGTGGTTGC |
| fwd_site_4 | ACACTCTTTCCCTACACGACGCTCTTCCGATCTNNNNGCCCATTCCCTCTTTAGCCA |
| rev_site_4 | TGGAGTTCAGACGTGTGCTCTTCCGATCTGAGCCGTTCCCTCTTTGCTA |
| fwd_site_5 | ACACTCTTTCCCTACACGACGCTCTTCCGATCTNNNNAACCTGTGTGACACTTGGCA |
| rev_site_5 | TGGAGTTCAGACGTGTGCTCTTCCGATCTGTCTGGCCCAAGATCACACA |
| fwd_site_6 | ACACTCTTTCCCTACACGACGCTCTTCCGATCTNNNCACGGATAAAGACGCTGGGA |
| rev_site_6 | TGGAGTTCAGACGTGTGCTCTTCCGATCTGGGGTCCCAGGTGCTGAC |
| fwd_site_7 | ACACTCTTTCCCTACACGACGCTCTTCCGATCTNNNTTGATTGTCTCCTTTGCCGC |
| rev_site_7 | TGGAGTTCAGACGTGTGCTCTTCCGATCTTGACCCAGTGTTTGATAGATCAGT |
| fwd_site_8 | ACACTCTTTCCCTACACGACGCTCTTCCGATCTNNNCACCCCTTCAGTCCATGCTT |
| rev_site_8 | TGGAGTTCAGACGTGTGCTCTTCCGATCTTCTGATGGGGAGGAACGAGT |
| fwd_site_9 | ACACTCTTTCCCTACACGACGCTCTTCCGATCTNNNNCAGCTCAGCCTGAGTGTTGA |
| rev_site_9 | TGGAGTTCAGACGTGTGCTCTTCCGATCTGCCCACCCTAGTCATTGGAG |
| fwd_site_10 | ACACTCTTTCCCTACACGACGCTCTTCCGATCTNNNNGTCAGAGGGACACACTGTGG |
| rev_site_10 | TGGAGTTCAGACGTGTGCTCTTCCGATCTCACACTCACTCACCCACACA |
| fwd_site_11 | ACACTCTTTCCCTACACGACGCTCTTCCGATCTNNNNTGTGTGGGTGAGTGAGTGTG |
| rev_site_11 | TGGAGTTCAGACGTGTGCTCTTCCGATCTCACCAAGGTTCACAGCCTGA |
| fwd_site_12 | ACACTCTTTCCCTACACGACGCTCTTCCGATCTNNNNTTGTCTCTGCCTGTAGCTGC |
| rev_site_12 | TGGAGTTCAGACGTGTGCTCTTCCGATCTCGCTCTGGGCTTCATCTTCA |
| fwd_site_13 | ACACTCTTTCCCTACACGACGCTCTTCCGATCTNNNNTGGGATTATGGGTGTGAGCC |
| rev_site_13 | TGGAGTTCAGACGTGTGCTCTTCCGATCTTGCCTTCCTCCTCTCTCTCC |
| fwd_site_14 | ACACTCTTTCCCTACACGACGCTCTTCCGATCTNNNNTGCAGACCAGATTCGGAGAA |
| rev_site_14 | TGGAGTTCAGACGTGTGCTCTTCCGATCTGTTCAGTTTCCAGGGGGTCC |
| fwd_site_15 | ACACTCTTTCCCTACACGACGCTCTTCCGATCTNNNNTCCGCACAGCCTTAGTTCAA |
| rev_site_15 | TGGAGTTCAGACGTGTGCTCTTCCGATCTAACTTGAAGAGACGGCAGCA |
| fwd_site_16 | ACACTCTTTCCCTACACGACGCTCTTCCGATCTNNNNCCCCAGCTACAGAAAGGTC |
| rev_site_16 | TGGAGTTCAGACGTGTGCTCTTCCGATCTATTTCCACCGCAAAATGGCC |
| fwd_site_17 | ACACTCTTTCCCTACACGACGCTCTTCCGATCTNNNNTCACTTCAGCCCAGGAGTAT |
| rev_site_17 | TGGAGTTCAGACGTGTGCTCTTCCGATCTTGTGTATGGTGAGAGGTAGGGA |
| fwd_site_18 | ACACTCTTTCCCTACACGACGCTCTTCCGATCTNNNNGTCTGAGGTCACACAGTGGG |
| rev_site_18 | TGGAGTTCAGACGTGTGCTCTTCCGATCTCTGAGAGCAGGGACCACATC |
| fwd_site_19 | ACACTCTTTCCCTACACGACGCTCTTCCGATCTNNNNGGGAGGTGGAGAGAGGATGT |

| | |
|---|---|
| **rev_site_19** | TGGAGTTCAGACGTGTGCTCTTCCGATCTACTCTTCCTGAGGTCTAGGAACCCG |
| **fwd_site_20** | ACACTCTTTCCCTACACGACGCTCTTCCGATCTNNNNCCCTGTTCCTAAAGCCCACC |
| **rev_site_20** | TGGAGTTCAGACGTGTGCTCTTCCGATCTACTCTCTGGTTCTGTTTGTGGCCA |
| **fwd_CTNNB1** | ACACTCTTTCCCTACACGACGCTCTTCCGATCTNNNNATTTGATGGAGTTGGACATGGCC |
| **rev_CTNNB1** | TGGAGTTCAGACGTGTGCTCTCCAGCTACTTGTTCTTGAGTGAAGG |
| **fwd_IP90** | ACACTCTTTCCCTACACGACGCTCTTCCGATCTNNNNCTGGTTGACCAATCTGTGGTG |
| **rev_IP90** | TGGAGTTCAGACGTGTGCTCTCTGCGTCTGGATCAGGTACG |
| **fwd_HEK293_site2_off1** | ACACTCTTTCCCTACACGACGCTCTTCCGATCTNNNNGTGTGGAGAGTGAGTAAGCCA |
| **rev_HEK293_site2_off1** | TGGAGTTCAGACGTGTGCTCTTCCGATCTACGGTAGGATGATTTCAGGCA |
| **fwd_HEK293_site2_off2** | ACACTCTTTCCCTACACGACGCTCTTCCGATCTNNNNCACAAAGCAGTGTAGCTCAGG |
| **rev_HEK293_site2_off2** | TGGAGTTCAGACGTGTGCTCTTCCGATCTTTTTTGGTACTCGAGTGTTATTCAG |
| **fwd_HEK293_site3_off1** | ACACTCTTTCCCTACACGACGCTCTTCCGATCTNNNNTCCCCTGTTGACCTGGAGAA |
| **rev_HEK293_site3_off1** | TGGAGTTCAGACGTGTGCTCTTCCGATCTCACTGTACTTGCCCTGACCA |
| **fwd_HEK293_site3_off2** | ACACTCTTTCCCTACACGACGCTCTTCCGATCTNNNNTTGGTGTTGACAGGGAGCAA |
| **rev_HEK293_site3_off2** | TGGAGTTCAGACGTGTGCTCTTCCGATCTCTGAGATGTGGGCAGAAGGG |
| **fwd_HEK293_site3_off3** | ACACTCTTTCCCTACACGACGCTCTTCCGATCTNNNNTGAGAGGGAACAGAAGGGCT |
| **rev_HEK293_site3_off3** | TGGAGTTCAGACGTGTGCTCTTCCGATCTGTCCAAAGGCCCAAGAACCT |
| **fwd_HEK293_site3_off4** | ACACTCTTTCCCTACACGACGCTCTTCCGATCTNNNNTCCTAGCACTTTGGAAGGTCG |
| **rev_HEK293_site3_off4** | TGGAGTTCAGACGTGTGCTCTTCCGATCTGCTCATCTTAATCTGCTCAGCC |
| **fwd_HEK293_site3_off5** | ACACTCTTTCCCTACACGACGCTCTTCCGATCTNNNNAAAGGAGCAGCTCTTCCTGG |
| **rev_HEK293_site3_off5** | TGGAGTTCAGACGTGTGCTCTTCCGATCTGTCTGCACCATCTCCCACAA |
| **fwd_HEK293_site4_off1** | ACACTCTTTCCCTACACGACGCTCTTCCGATCTNNNNGGCATGGCTTCTGAGACTCA |
| **rev_HEK293_site4_off1** | TGGAGTTCAGACGTGTGCTCTTCCGATCTGTCTCCCTTGCACTCCCTGTCTTT |
| **fwd_HEK293_site4_off2** | ACACTCTTTCCCTACACGACGCTCTTCCGATCTNNNNTTTGGCAATGGAGGCATTGG |
| **rev_HEK293_site4_off2** | TGGAGTTCAGACGTGTGCTCTTCCGATCTGAAGAGGCTGCCCATGAGAG |
| **fwd_HEK293_site4_off3** | ACACTCTTTCCCTACACGACGCTCTTCCGATCTNNNNGGTCTGAGGCTCGAATCCTG |
| **rev_HEK293_site4_off3** | TGGAGTTCAGACGTGTGCTCTTCCGATCTCTGTGGCCTCCATATCCCTG |
| **fwd_HEK293_site4_off4** | ACACTCTTTCCCTACACGACGCTCTTCCGATCTNNNNTTTCCACCAGAACTCAGCCC |
| **rev_HEK293_site4_off4** | TGGAGTTCAGACGTGTGCTCTTCCGATCTCCTCGGTTCCTCCACAACAC |
| **fwd_HEK293_site4_off5** | ACACTCTTTCCCTACACGACGCTCTTCCGATCTNNNNCACGGGAAGGACAGGAGAAG |
| **rev_HEK293_site4_off5** | TGGAGTTCAGACGTGTGCTCTTCCGATCTGCAGGGGAGGGATAAAGCAG |
| **fwd_HEK_site_3** | ACACTCTTTCCCTACACGACGCTCTTCCGATCTNNNNGGAAACGCCCATGCAATTAGTC |
| **rev_HEK_site_3** | TGGAGTTCAGACGTGTGCTCTTCCGATCTCTTGTCAACCAGTATCCCGGTG |
| **fwd_HEK_site_2** | ACACTCTTTCCCTACACGACGCTCTTCCGATCTNNNNTGAATGGATTCCTTGGAAACAATG |
| **rev_HEK_site_2** | TGGAGTTCAGACGTGTGCTCTTCCGATCTCCAGCCCCATCTGTCAAACT |
| **fwd_HEK_site_4** | TGGAGTTCAGACGTGTGCTCTTCCGATCTTCCTTTCAACCCGAACGGAG |
| **rev_HEK_site_4** | ACACTCTTTCCCTACACGACGCTCTTCCGATCTNNNNGCTGGTCTTCTTTCCCCTCC |
| **fwd_LDLR** | ACACTCTTTCCCTACACGACGCTCTTCCGATCTNNNNGCCCTGCTTCTTTTTCTCTGGT |
| **rev_LDLR** | TGGAGTTCAGACGTGTGCTCTTCCGATCTACCATTAACGCAGCCAACTTCA |
| **fwd_TRAC** | ACACTCTTTCCCTACACGACGCTCTTCCGATCTCATGAGGTCTATGGACTTCAAGAGCAA |

| | |
|---|---|
| **Rev_TRAC** | TGGAGTTCAGACGTGTGCTCTTCCGATCTCATCATTGACCAGAGCTCTGGGCAGAA |
| **fwd_CBLB** | ACACTCTTTCCCTACACGACGCTCTTCCGATCTGCACTTACCAGCATTACTTCCTAAACC |
| **Rev_CBLB** | TGGAGTTCAGACGTGTGCTCTTCCGATCTATGGGCTCCACTTTTCAGCTCTGTAA |
| **fwd_CD7** | ACACTCTTTCCCTACACGACGCTCTTCCGATCTCAGTTCAGGCACATGTAGGAGGGA |
| **Rev_CD7** | TGGAGTTCAGACGTGTGCTCTTCCGATCTACCGCCTGCAGCTGTCGGACACTGGCA |
| **fwd_B2M** | ACACTCTTTCCCTACACGACGCTCTTCCGATCTAAAAGATGAGTATGCCTGCCGTG |
| **Rev_B2M** | TGGAGTTCAGACGTGTGCTCTTCCGATCTCAGATTGTTTATATCAGATGGGATGGG |
| **fwd_CIITA** | ACACTCTTTCCCTACACGACGCTCTTCCGATCTATGCAAGTTTGGTCCTGAGCCCTCCC |
| **Rev_CIITA** | TGGAGTTCAGACGTGTGCTCTTCCGATCTGATGTGGGTTCCCTGCGCTCTGCA |
| **fwd_PDCD1** | ACACTCTTTCCCTACACGACGCTCTTCCGATCTCCAGGGACTGAGGGTGGAAGGTCC |
| **Rev_PDCD1** | TGGAGTTCAGACGTGTGCTCTTCCGATCTACCTCCGCCTGAGCAGTGGAGAA |

**Supplementary Table 5:** UHPLC UV-Vis trace (220 nm) and integration of globin chain levels of untreated differentiated CD34+ cells (donor 1)

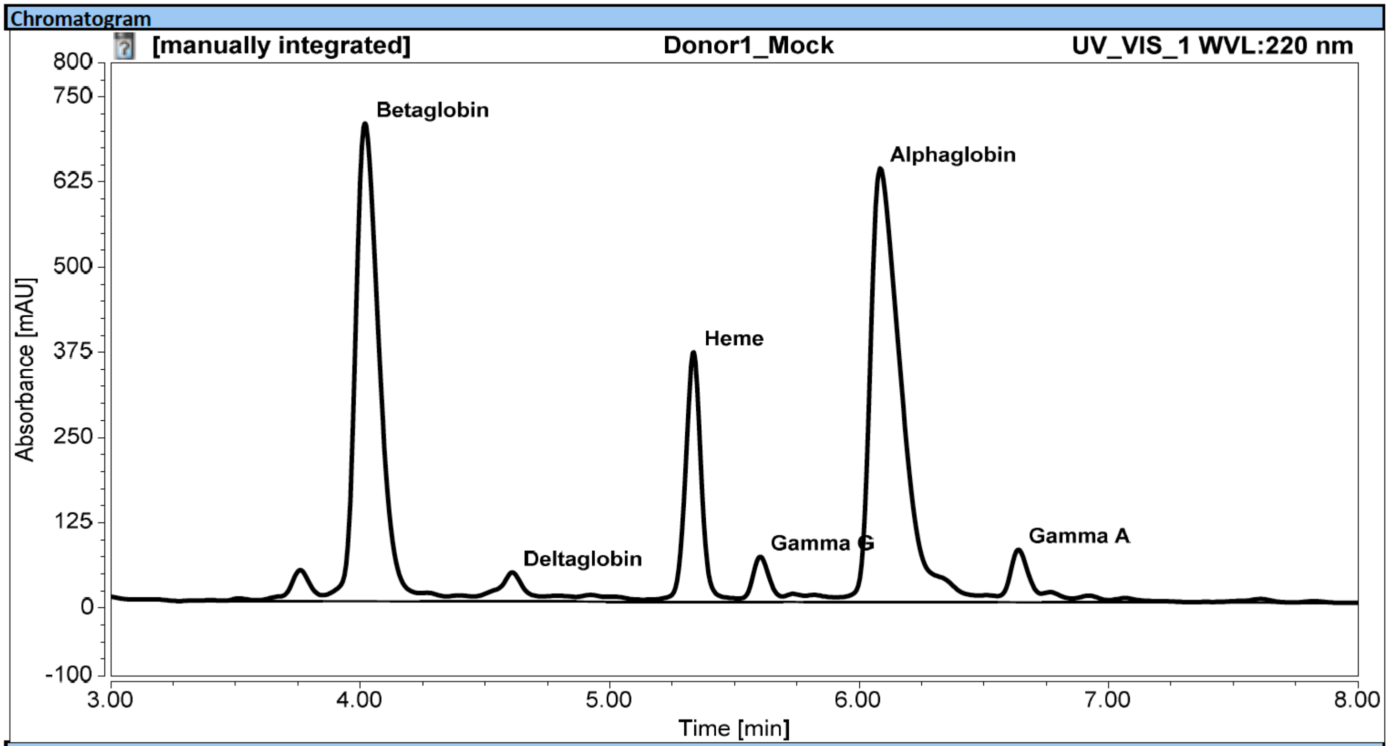


Integration Results

| No. | Peak Name | Retention Time<br>min | Area<br>mAU*min | Height<br>mAU | Relative Area<br>% | Relative Height<br>% |
|---|---|---|---|---|---|---|
| 6 | Betaglobin | 4.020 | 77.631 | 701.634 | 33.61 | 32.98 |
| n.a. | HbS | n.a. | n.a. | n.a. | n.a. | n.a. |
| 10 | Deltaglobin | 4.610 | 3.780 | 42.943 | 1.64 | 2.02 |
| 14 | Heme | 5.337 | 27.191 | 366.542 | 11.77 | 17.23 |
| 15 | Gamma G | 5.603 | 5.248 | 66.622 | 2.27 | 3.13 |
| 18 | Alphaglobin | 6.083 | 88.477 | 636.811 | 38.31 | 29.94 |
| 21 | Gamma A | 6.637 | 6.751 | 77.427 | 2.92 | 3.64 |

**Supplementary Table 6:** UHPLC UV-Vis trace (220 nm) and integration of globin chain levels of differentiated CD34+ cells treated with ABE7.10-m (donor1)

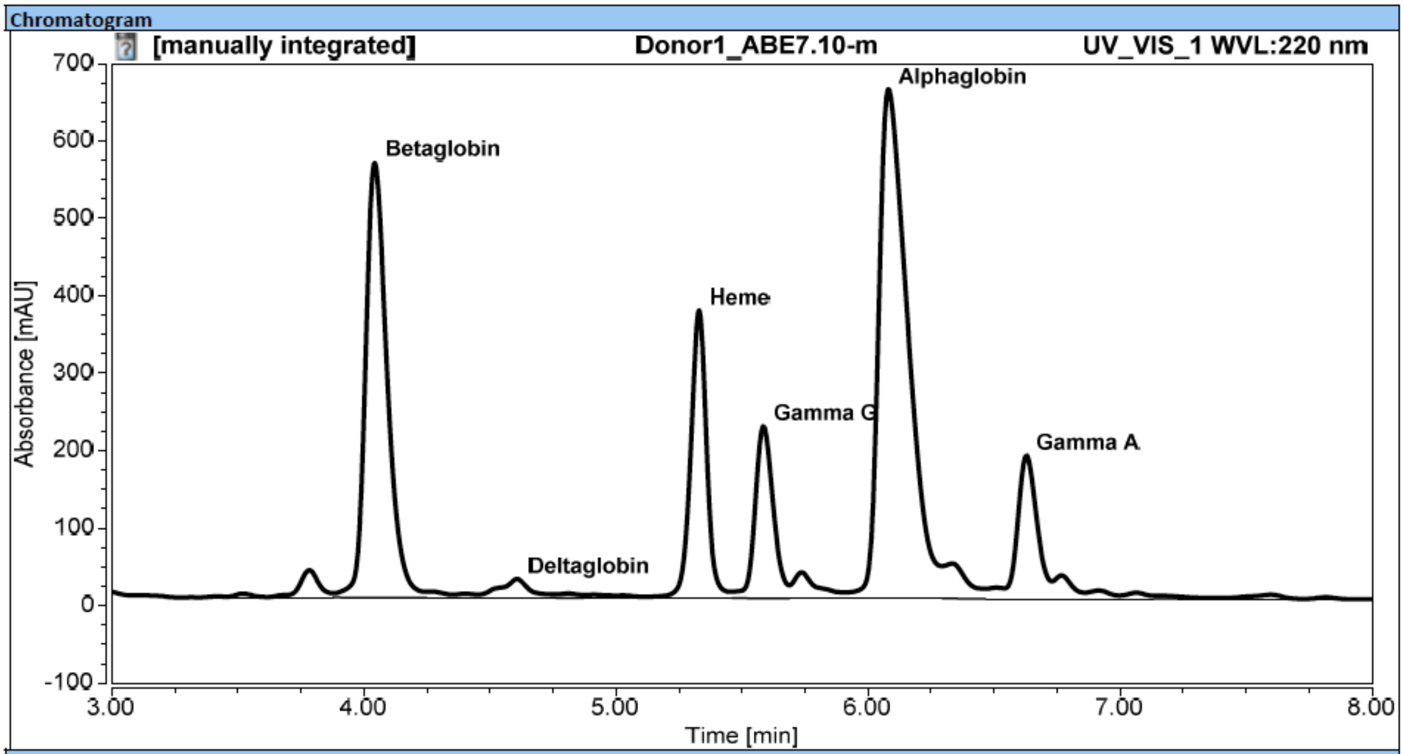


Integration Results

| No. | Peak Name | Retention Time min | Area mAU*min | Height mAU | Relative Area % | Relative Height % |
|---|---|---|---|---|---|---|
| 5 | Betaglobin | 4.043 | 56.404 | 561.657 | 24.21 | 24.80 |
| n.a. | HbS | n.a. | n.a. | n.a. | n.a. | n.a. |
| 9 | Deltaglobin | 4.607 | 2.463 | 24.382 | 1.06 | 1.08 |
| 13 | Heme | 5.330 | 27.334 | 371.802 | 11.73 | 16.42 |
| 14 | Gamma G | 5.583 | 18.764 | 222.410 | 8.06 | 9.82 |
| 16 | Alphaglobin | 6.080 | 88.236 | 657.695 | 37.88 | 29.04 |
| 19 | Gamma A | 6.627 | 16.470 | 185.055 | 7.07 | 8.17 |

**Supplementary Table 7:** UHPLC UV-Vis trace (220 nm) and integration of globin chain levels of differentiated CD34+ cells treated with ABE7.10-d (donor1).

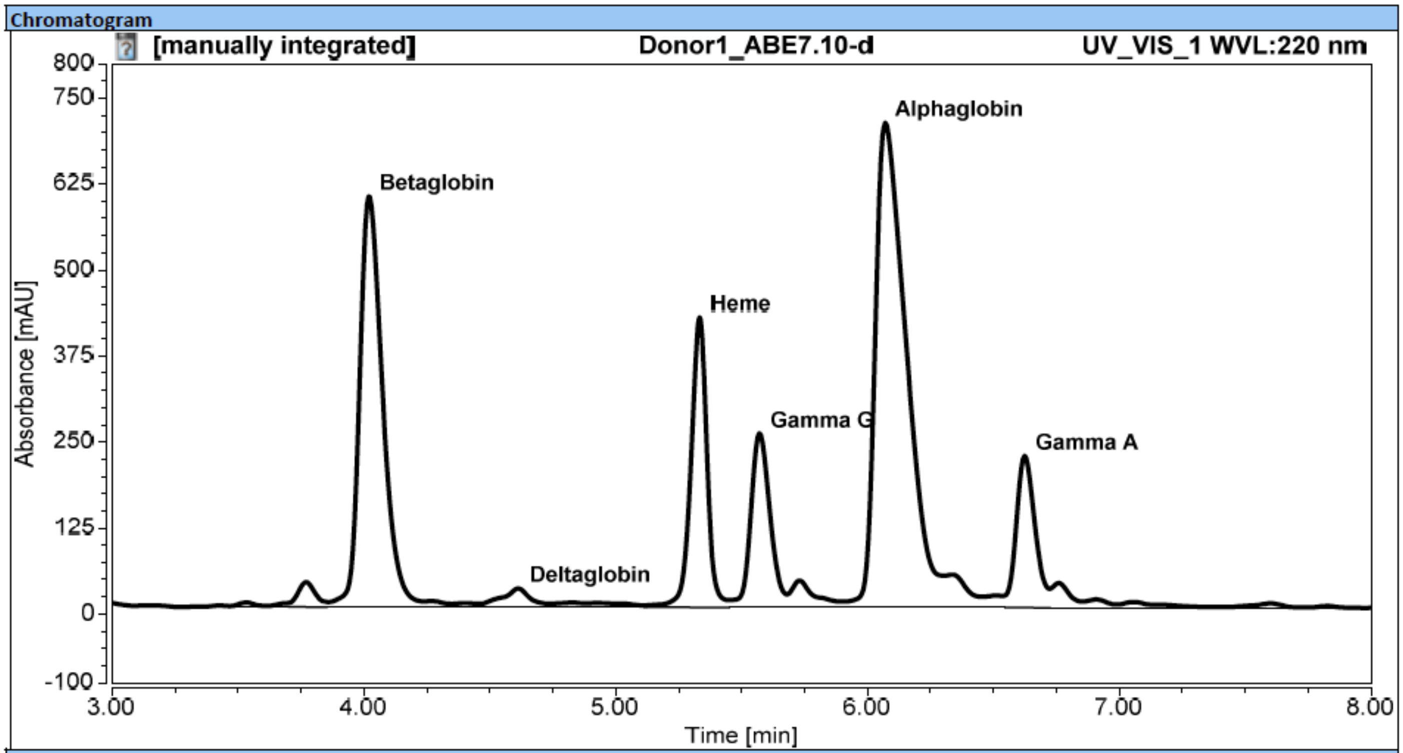


Integration Results

| No. | Peak Name | Retention Time min | Area mAU*min | Height mAU | Relative Area % | Relative Height % |
|---|---|---|---|---|---|---|
| 6 | Betaglobin | 4.020 | 63.540 | 596.818 | 23.88 | 23.74 |
| n.a. | HbS | n.a. | n.a. | n.a. | n.a. | n.a. |
| 10 | Deltaglobin | 4.613 | 2.822 | 27.354 | 1.06 | 1.09 |
| 14 | Heme | 5.333 | 31.163 | 421.735 | 11.71 | 16.77 |
| 15 | Gamma G | 5.570 | 22.558 | 253.399 | 8.48 | 10.08 |
| 18 | Alphaglobin | 6.070 | 100.612 | 704.910 | 37.81 | 28.03 |
| 21 | Gamma A | 6.623 | 19.220 | 220.846 | 7.22 | 8.78 |

**Supplementary Table 8:** UHPLC UV-Vis trace (220 nm) and integration of globin chain levels of differentiated CD34+ cells treated with ABE8.8-m (donor1)

Chromatogram

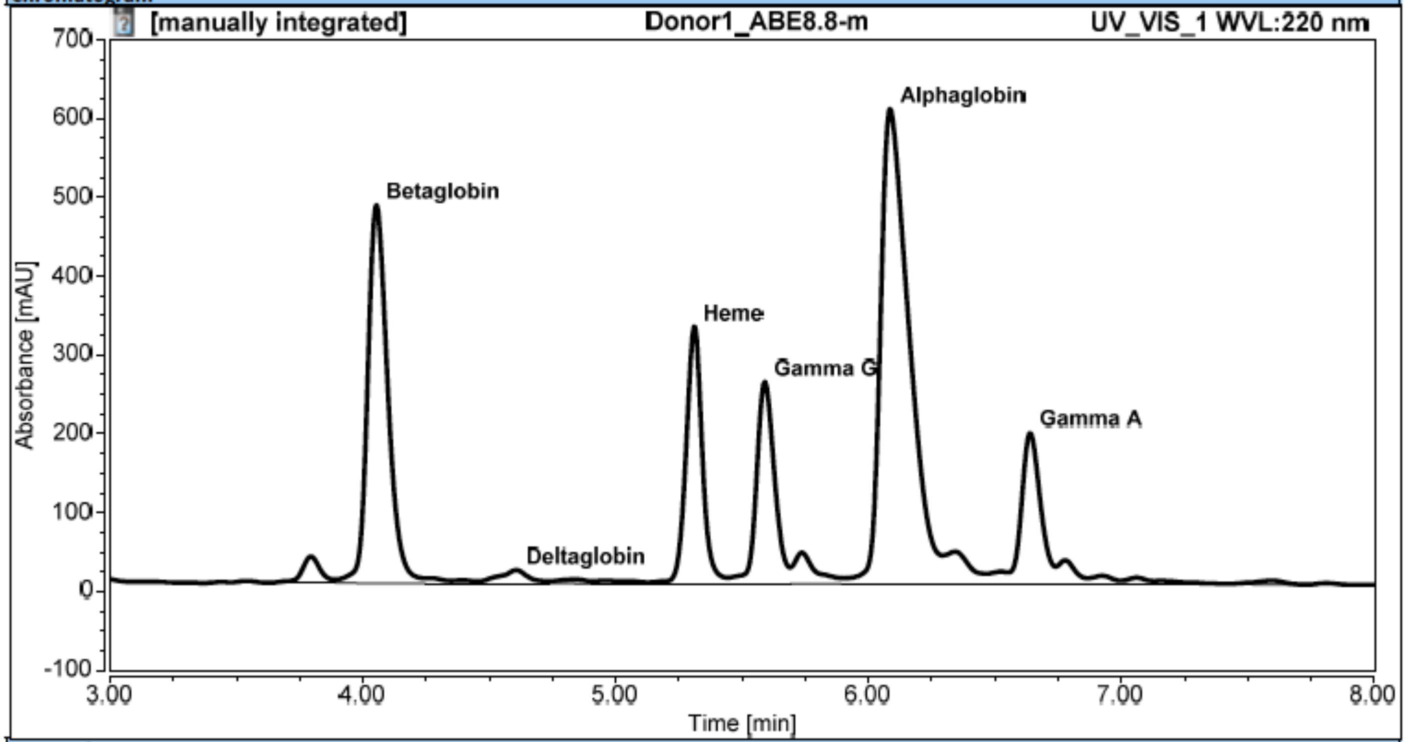


Integration Results

| No. | Peak Name | Retention Time<br>min | Area<br>mAU*min | Height<br>mAU | Relative Area<br>% | Relative Height<br>% |
|---|---|---|---|---|---|---|
| 5 | Betaglobin | 4.053 | 44.227 | 479.648 | 21.03 | 22.78 |
| n.a. | HbS | n.a. | n.a. | n.a. | n.a. | n.a. |
| 9 | Deltaglobin | 4.607 | 1.704 | 17.056 | 0.81 | 0.81 |
| 13 | Heme | 5.310 | 24.221 | 326.068 | 11.52 | 15.48 |
| 14 | Gamma G | 5.590 | 21.346 | 256.462 | 10.15 | 12.18 |
| 16 | Alphaglobin | 6.087 | 79.706 | 602.792 | 37.90 | 28.62 |
| 19 | Gamma A | 6.640 | 16.631 | 192.016 | 7.91 | 9.12 |

**Supplementary Table 9:** UHPLC UV-Vis trace (220 nm) and integration of globin chain levels of differentiated CD34+ cells treated with ABE8.8-d (donor1).

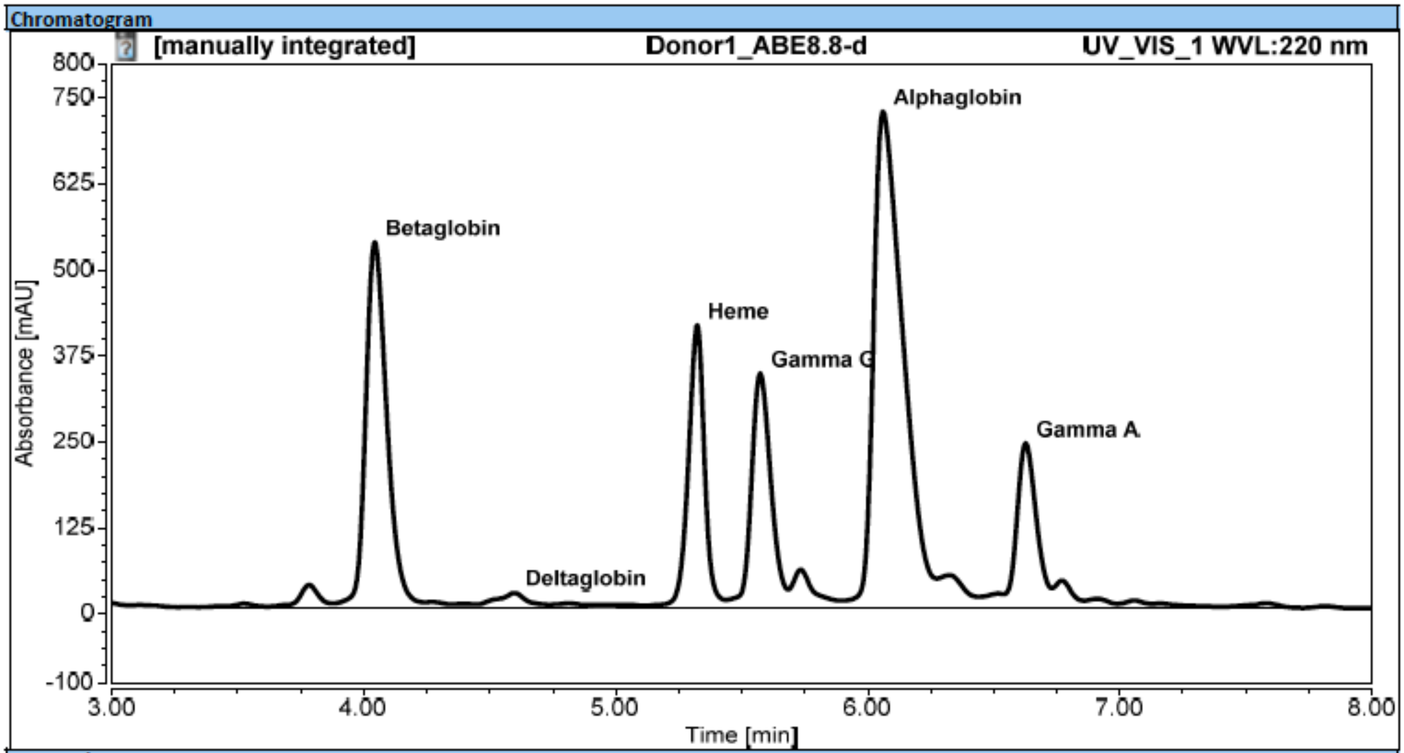


Integration Results

| No. | Peak Name | Retention Time<br>min | Area<br>mAU*min | Height<br>mAU | Relative Area<br>% | Relative Height<br>% |
|---|---|---|---|---|---|---|
| 5 | Betaglobin | 4.043 | 51.802 | 530.669 | 19.60 | 20.80 |
| n.a. | HbS | n.a. | n.a. | n.a. | n.a. | n.a. |
| 9 | Deltaglobin | 4.597 | 2.269 | 21.040 | 0.86 | 0.82 |
| 13 | Heme | 5.323 | 30.637 | 410.481 | 11.59 | 16.09 |
| 14 | Gamma G | 5.573 | 30.747 | 340.883 | 11.63 | 13.36 |
| 16 | Alphaglobin | 6.060 | 98.417 | 721.350 | 37.24 | 28.28 |
| 19 | Gamma A | 6.627 | 21.595 | 239.647 | 8.17 | 9.39 |

**Supplementary Table 10:** UHPLC UV-Vis trace (220 nm) and integration of globin chain levels of differentiated CD34+ cells treated with ABE8.13-m (donor1).

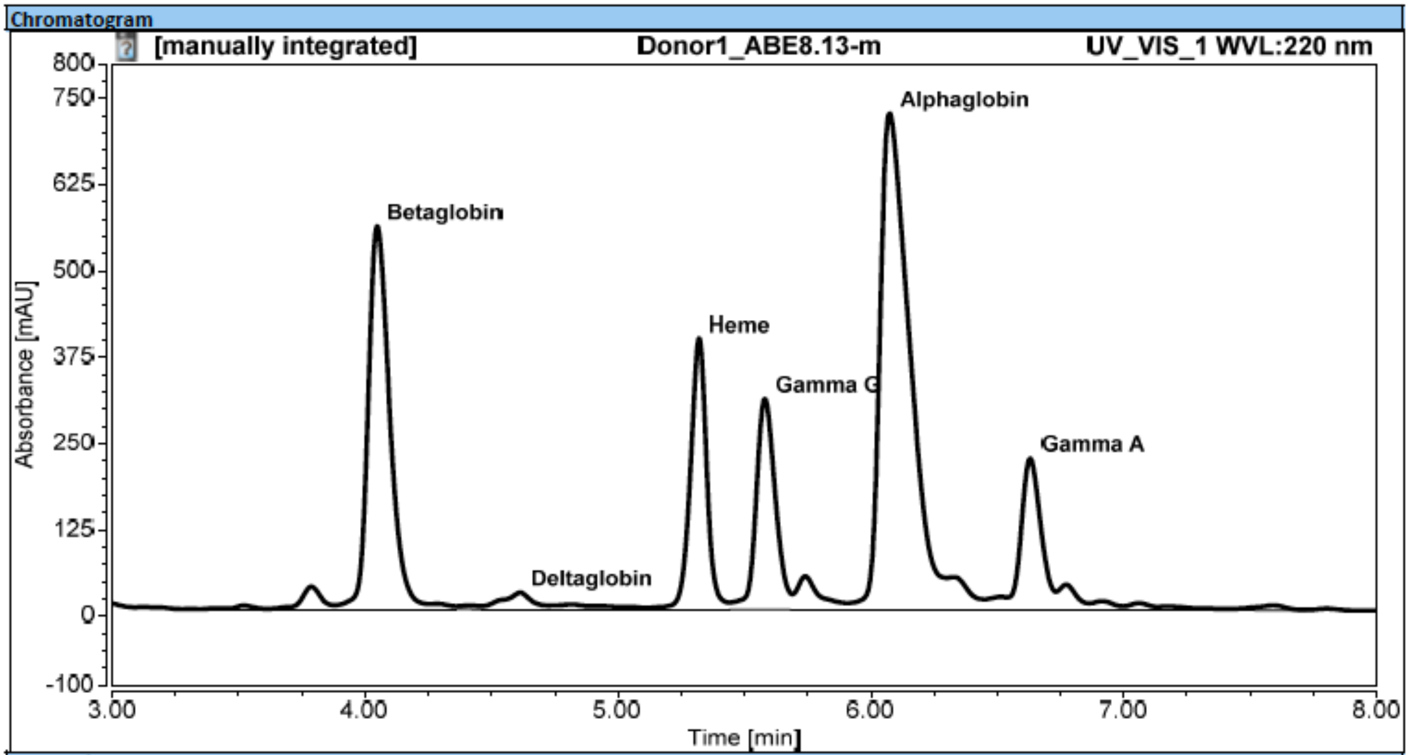


Integration Results

| No. | Peak Name | Retention Time min | Area mAU*min | Height mAU | Relative Area % | Relative Height % |
|---|---|---|---|---|---|---|
| 5 | Betaglobin | 4.047 | 55.084 | 554.598 | 21.23 | 22.22 |
| n.a. | HbS | n.a. | n.a. | n.a. | n.a. | n.a. |
| 9 | Deltaglobin | 4.613 | 2.417 | 24.372 | 0.93 | 0.98 |
| 13 | Heme | 5.320 | 29.500 | 393.066 | 11.37 | 15.75 |
| 14 | Gamma G | 5.580 | 27.177 | 306.655 | 10.48 | 12.29 |
| 16 | Alphaglobin | 6.073 | 98.048 | 719.982 | 37.79 | 28.85 |
| 19 | Gamma A | 6.630 | 19.949 | 219.869 | 7.69 | 8.81 |

**Supplementary Table 11:** UHPLC UV-Vis trace (220 nm) and integration of globin chain levels of differentiated CD34+ cells treated with ABE8.13-d (donor1).

Chromatogram

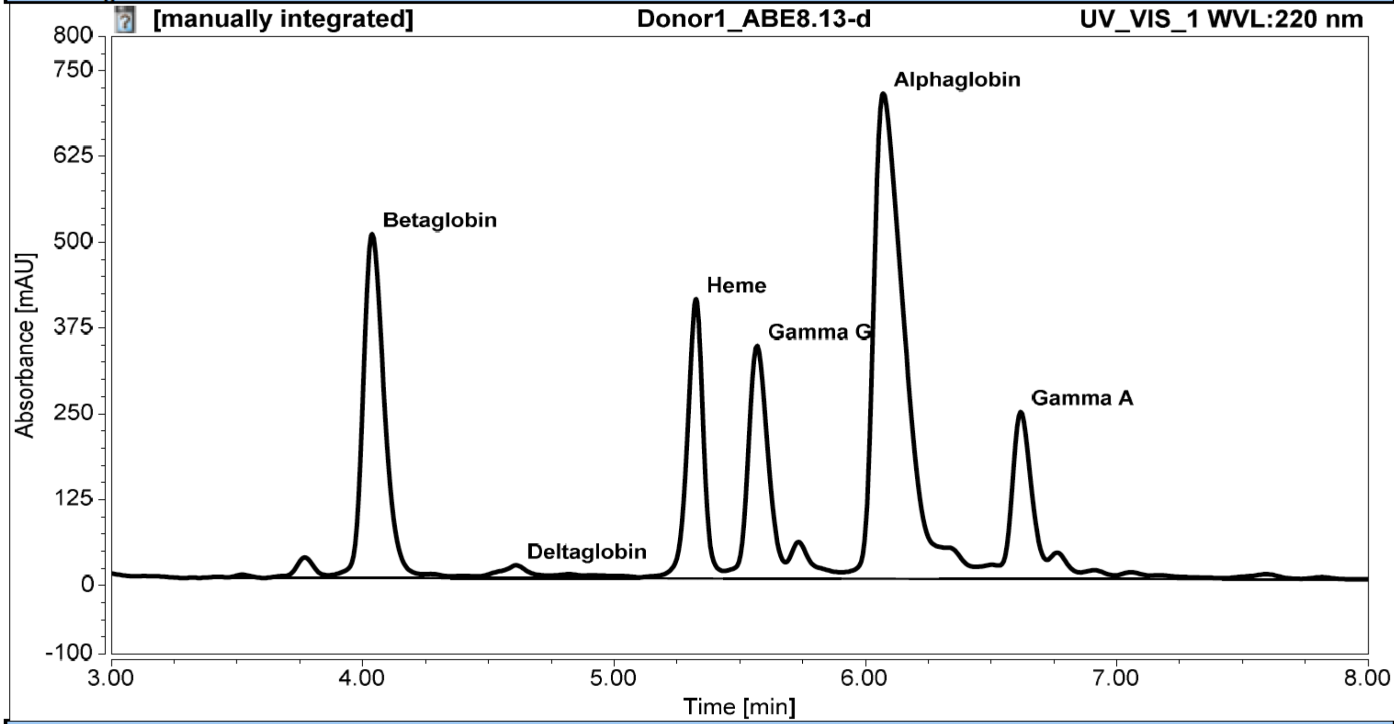


Integration Results

| No. | Peak Name | Retention Time min | Area mAU*min | Height mAU | Relative Area % | Relative Height % |
|---|---|---|---|---|---|---|
| 5 | Betaglobin | 4.037 | 49.284 | 501.306 | 19.04 | 20.14 |
| n.a. | HbS | n.a. | n.a. | n.a. | n.a. | n.a. |
| 9 | Deltaglobin | 4.610 | 1.960 | 18.855 | 0.76 | 0.76 |
| 13 | Heme | 5.327 | 30.274 | 407.530 | 11.69 | 16.37 |
| 14 | Gamma G | 5.570 | 31.059 | 339.163 | 12.00 | 13.63 |
| 16 | Alphaglobin | 6.070 | 98.109 | 707.464 | 37.89 | 28.42 |
| 19 | Gamma A | 6.617 | 21.869 | 243.715 | 8.45 | 9.79 |

**Supplementary Table 12:** UHPLC UV-Vis trace (220 nm) and integration of globin chain levels of differentiated CD34+ cells treated with ABE8.17-m (donor1).

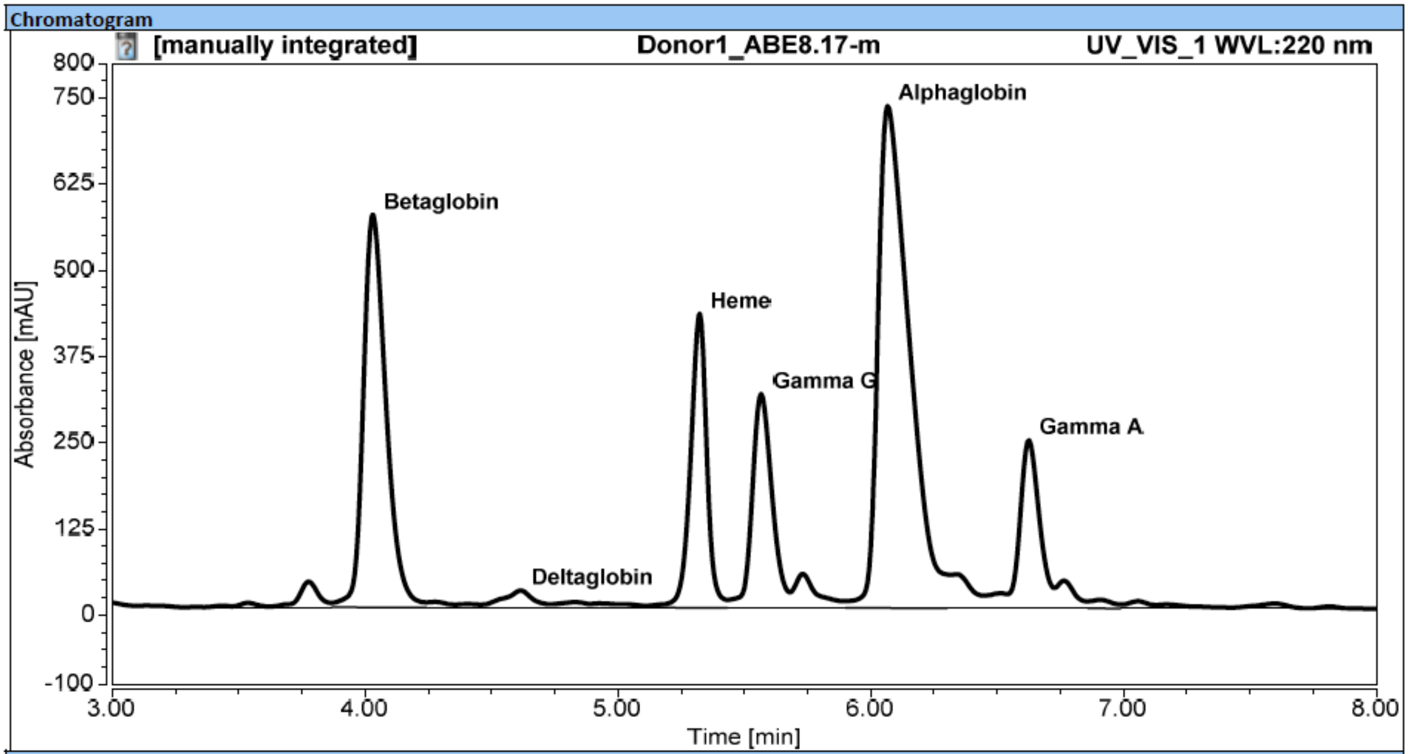


Integration Results

| No. | Peak Name | Retention Time<br>min | Area<br>mAU*min | Height<br>mAU | Relative Area<br>% | Relative Height<br>% |
|---|---|---|---|---|---|---|
| 5 | Betaglobin | 4.030 | 59.000 | 569.475 | 21.35 | 21.91 |
| n.a. | HbS | n.a. | n.a. | n.a. | n.a. | n.a. |
| 9 | Deltaglobin | 4.617 | 2.483 | 24.660 | 0.90 | 0.95 |
| 13 | Heme | 5.323 | 31.792 | 426.360 | 11.50 | 16.40 |
| 14 | Gamma G | 5.567 | 28.258 | 310.274 | 10.22 | 11.94 |
| 16 | Alphaglobin | 6.067 | 104.586 | 728.120 | 37.84 | 28.01 |
| 19 | Gamma A | 6.627 | 21.512 | 243.710 | 7.78 | 9.37 |

**Supplementary Table 13:** UHPLC UV-Vis trace (220 nm) and integration of globin chain levels of differentiated CD34+ cells treated with ABE8.17-d (donor1).

Chromatogram

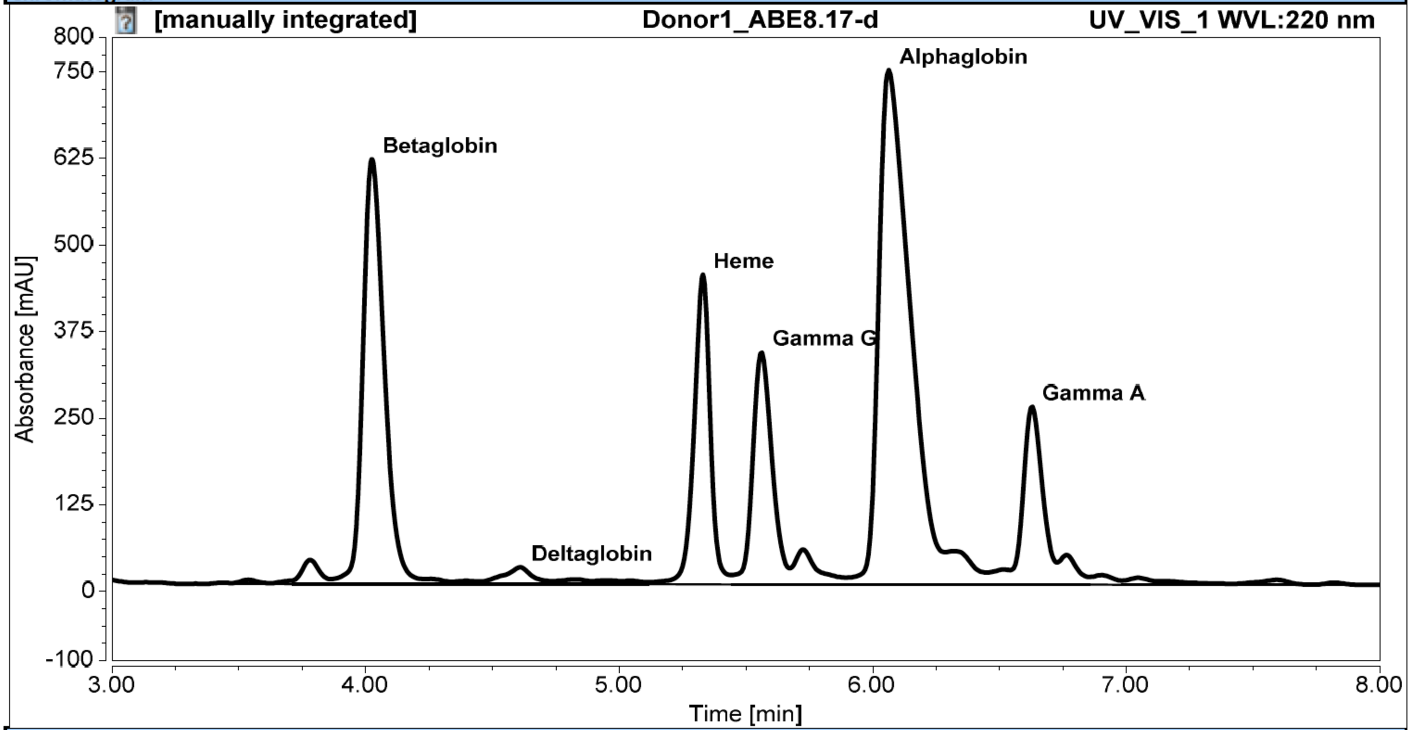


Integration Results

| No. | Peak Name | Retention Time min | Area mAU*min | Height mAU | Relative Area % | Relative Height % |
|---|---|---|---|---|---|---|
| 6 | Betaglobin | 4.023 | 60.046 | 613.639 | 21.18 | 22.63 |
| n.a. | HbS | n.a. | n.a. | n.a. | n.a. | n.a. |
| 10 | Deltaglobin | 4.610 | 2.491 | 24.450 | 0.88 | 0.90 |
| 14 | Heme | 5.330 | 33.236 | 446.899 | 11.72 | 16.48 |
| 15 | Gamma G | 5.560 | 29.840 | 335.709 | 10.52 | 12.38 |
| 17 | Alphaglobin | 6.063 | 106.077 | 742.712 | 37.41 | 27.38 |
| 20 | Gamma A | 6.627 | 22.570 | 257.472 | 7.96 | 9.49 |

**Supplementary Table 14:** UHPLC UV-Vis trace (220 nm) and integration of globin chain levels of differentiated CD34+ cells treated with ABE8.20-m (donor1).

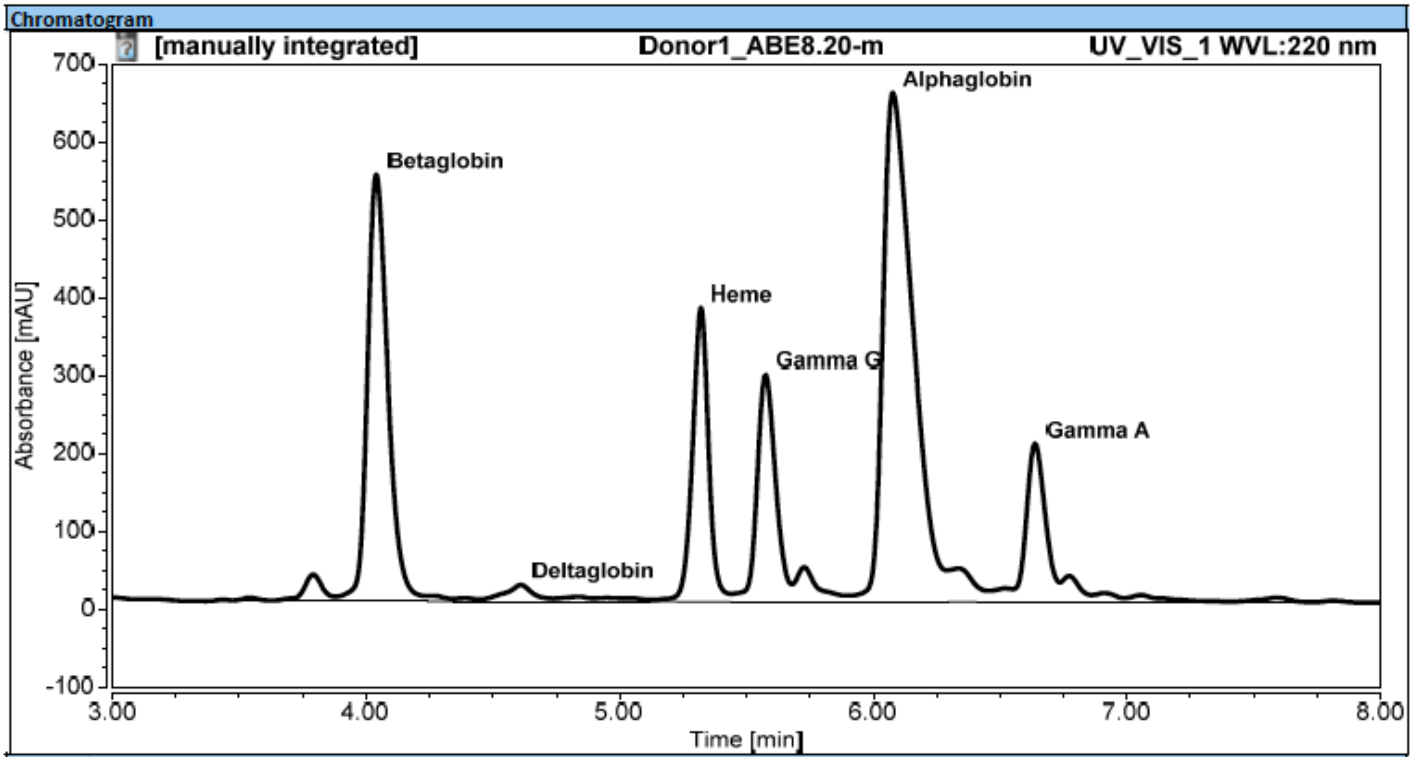


Integration Results

| No. | Peak Name | Retention Time min | Area mAU*min | Height mAU | Relative Area % | Relative Height % |
|---|---|---|---|---|---|---|
| 5 | Betaglobin | 4.040 | 51.610 | 548.050 | 21.57 | 23.31 |
| n.a. | HbS | n.a. | n.a. | n.a. | n.a. | n.a. |
| 9 | Deltaglobin | 4.610 | 2.233 | 21.329 | 0.93 | 0.91 |
| 13 | Heme | 5.320 | 27.916 | 377.668 | 11.67 | 16.06 |
| 14 | Gamma G | 5.573 | 24.931 | 291.767 | 10.42 | 12.41 |
| 16 | Alphaglobin | 6.077 | 90.099 | 653.942 | 37.66 | 27.81 |
| 19 | Gamma A | 6.637 | 17.798 | 203.835 | 7.44 | 8.67 |

**Supplementary Table 15:** UHPLC UV-Vis trace (220 nm) and integration of globin chain levels of differentiated CD34+ cells treated with ABE8.20-d (donor 1).

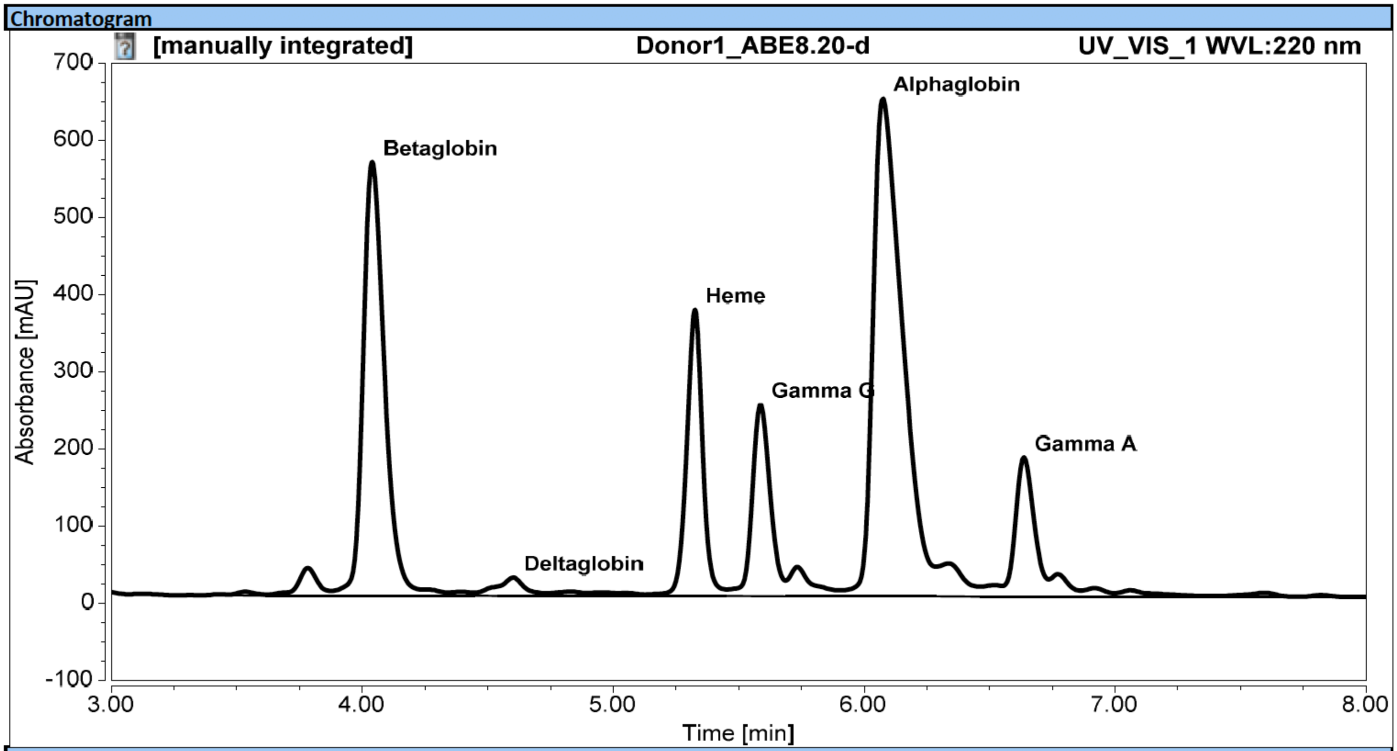


Integration Results

| No. | Peak Name | Retention Time<br>min | Area<br>mAU*min | Height<br>mAU | Relative Area<br>% | Relative Height<br>% |
|---|---|---|---|---|---|---|
| 6 | Betaglobin | 4.040 | 55.490 | 562.369 | 23.82 | 24.71 |
| n.a. | HbS | n.a. | n.a. | n.a. | n.a. | n.a. |
| 10 | Deltaglobin | 4.600 | 2.399 | 23.713 | 1.03 | 1.04 |
| 14 | Heme | 5.327 | 27.308 | 371.229 | 11.72 | 16.31 |
| 15 | Gamma G | 5.587 | 20.808 | 248.715 | 8.93 | 10.93 |
| 17 | Alphaglobin | 6.073 | 87.517 | 645.669 | 37.56 | 28.38 |
| 20 | Gamma A | 6.637 | 15.700 | 180.637 | 6.74 | 7.94 |

**Supplementary Table 16:** UHPLC UV-Vis trace (220 nm) and integration of globin chain levels of differentiated CD34+ cells untreated (donor 2). Note: donor 2 is heterozygous for sickle cell disease.

Chromatogram

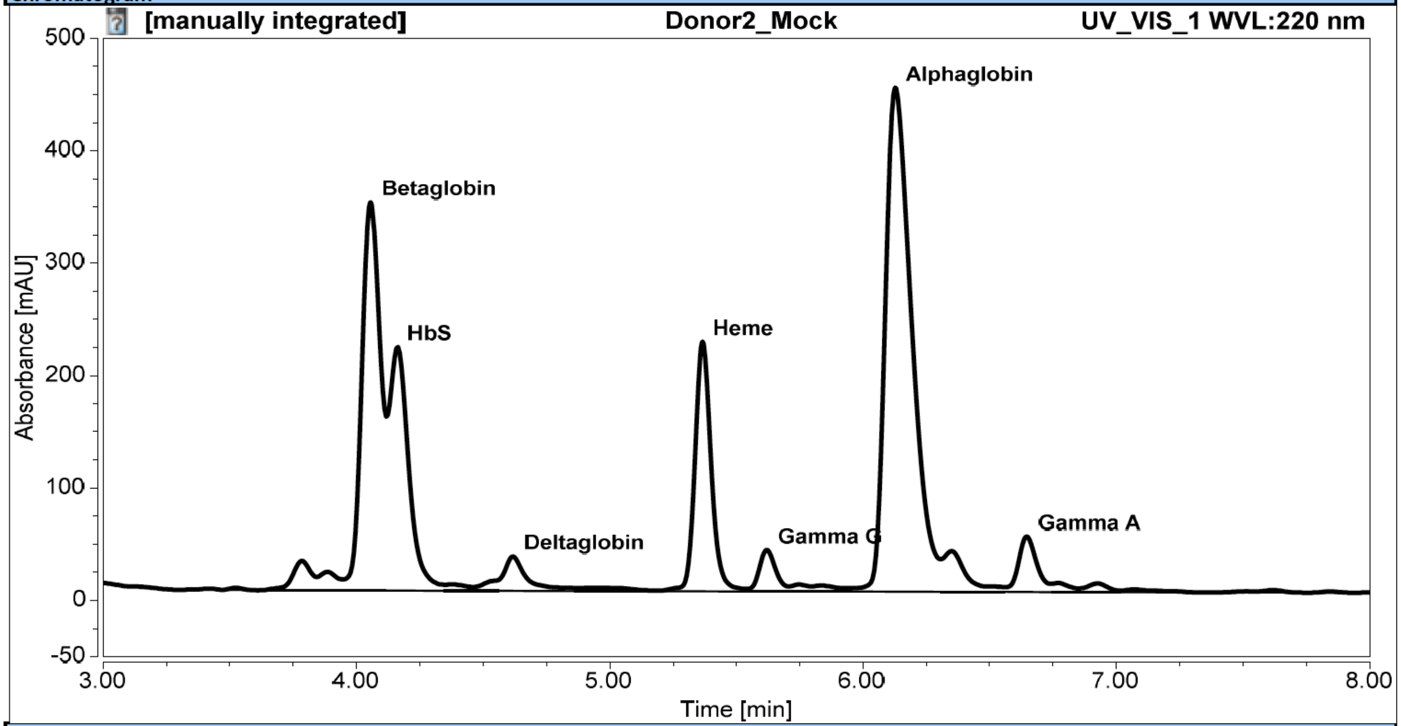


Integration Results

| No. | Peak Name | Retention Time min | Area mAU*min | Height mAU | Relative Area % | Relative Height % |
|---|---|---|---|---|---|---|
| 5 | Betaglobin | 4.057 | 28.943 | 345.410 | 20.25 | 23.19 |
| 6 | HbS | 4.160 | 19.794 | 216.879 | 13.85 | 14.56 |
| 9 | Deltaglobin | 4.617 | 3.170 | 30.593 | 2.22 | 2.05 |
| 11 | Heme | 5.367 | 16.498 | 222.155 | 11.54 | 14.91 |
| 12 | Gamma G | 5.620 | 2.805 | 37.002 | 1.96 | 2.48 |
| 15 | Alphaglobin | 6.127 | 54.386 | 448.385 | 38.05 | 30.10 |
| 18 | Gamma A | 6.647 | 4.170 | 49.520 | 2.92 | 3.32 |

**Supplementary Table 17:** UHPLC UV-Vis trace (220 nm) and integration of globin chain levels of differentiated CD34+ cells treated with ABE7.10-m (donor 2). Note: donor 2 is heterozygous for sickle cell disease.

Chromatogram

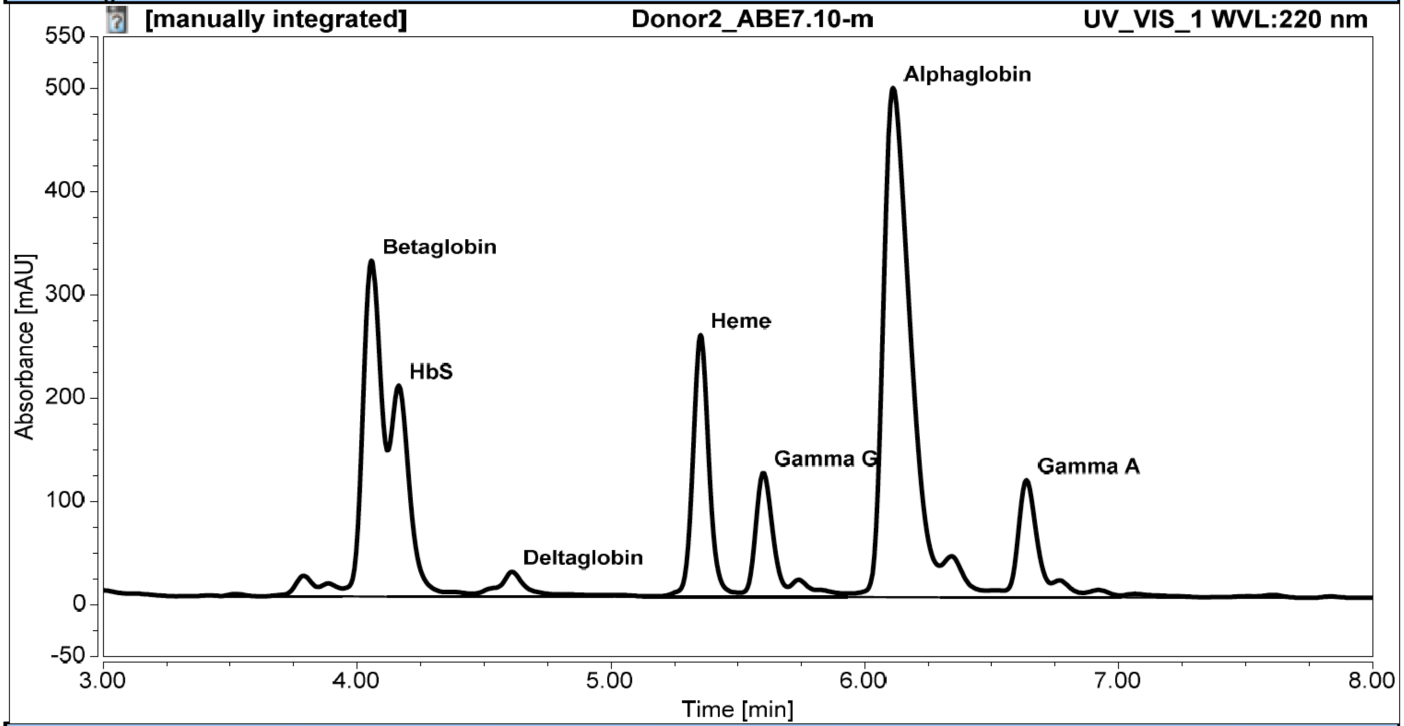


Integration Results

| No. | Peak Name | Retention Time<br>min | Area<br>mAU*min | Height<br>mAU | Relative Area<br>% | Relative Height<br>% |
|---|---|---|---|---|---|---|
| 5 | Betaglobin | 4.057 | 27.210 | 325.305 | 16.96 | 19.17 |
| 6 | HbS | 4.163 | 18.038 | 204.594 | 11.24 | 12.05 |
| 9 | Deltaglobin | 4.610 | 2.458 | 24.147 | 1.53 | 1.42 |
| 14 | Heme | 5.353 | 18.788 | 253.663 | 11.71 | 14.94 |
| 15 | Gamma G | 5.600 | 9.475 | 120.558 | 5.91 | 7.10 |
| 18 | Alphaglobin | 6.110 | 60.688 | 493.180 | 37.83 | 29.06 |
| 21 | Gamma A | 6.637 | 9.526 | 113.714 | 5.94 | 6.70 |

**Supplementary Table 18:** UHPLC UV-Vis trace (220 nm) and integration of globin chain levels of differentiated CD34+ cells treated with ABE7.10-d (donor 2). Note: donor 2 is heterozygous for sickle cell disease.

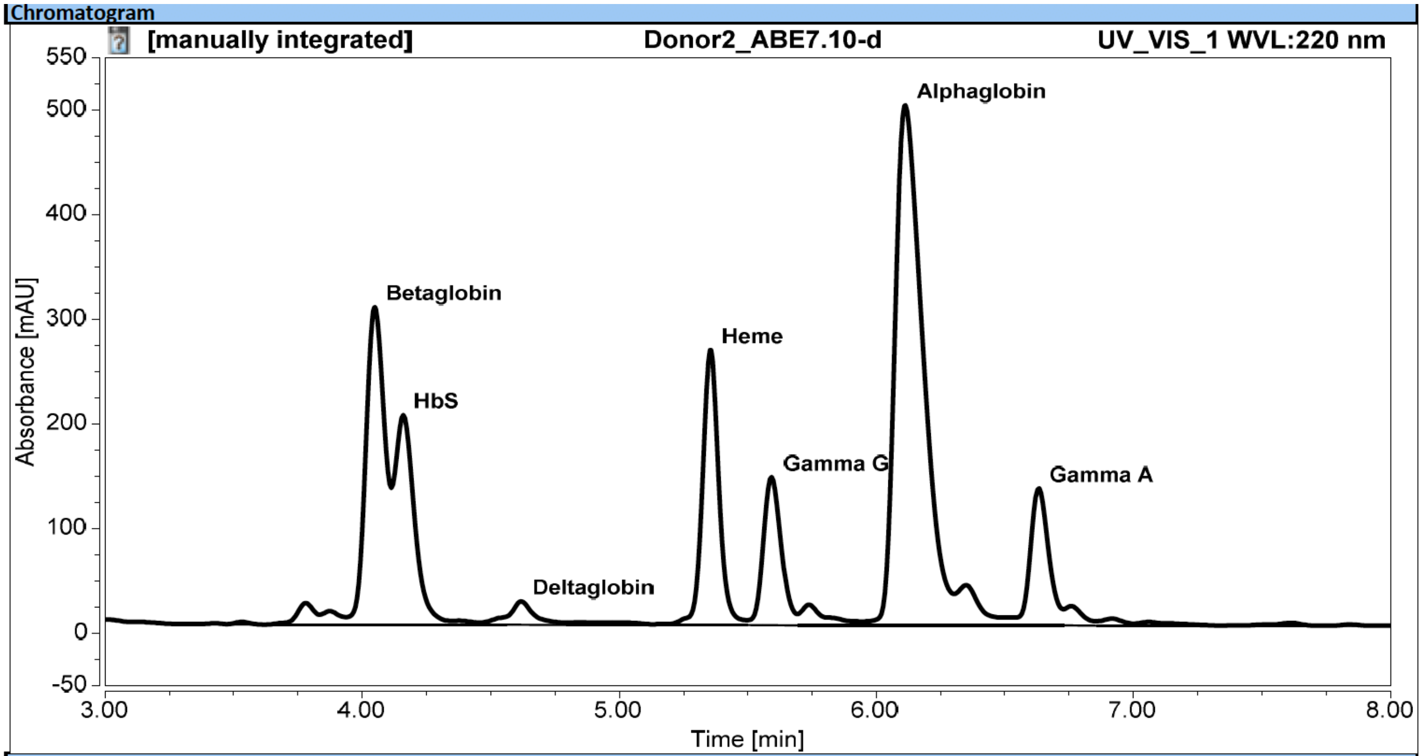


Integration Results

| No. | Peak Name | Retention Time min | Area mAU*min | Height mAU | Relative Area % | Relative Height % |
|---|---|---|---|---|---|---|
| 5 | Betaglobin | 4.050 | 26.525 | 303.461 | 15.99 | 17.61 |
| 6 | HbS | 4.160 | 17.741 | 200.547 | 10.70 | 11.64 |
| 9 | Deltaglobin | 4.620 | 2.209 | 22.450 | 1.33 | 1.30 |
| 14 | Heme | 5.357 | 19.493 | 262.669 | 11.75 | 15.24 |
| 15 | Gamma G | 5.593 | 11.528 | 141.729 | 6.95 | 8.23 |
| 18 | Alphaglobin | 6.113 | 62.859 | 497.019 | 37.90 | 28.85 |
| 21 | Gamma A | 6.630 | 10.827 | 131.062 | 6.53 | 7.61 |

**Supplementary Table 19:** UHPLC UV-Vis trace (220 nm) and integration of globin chain levels of differentiated CD34+ cells treated with ABE8.8-m (donor 2). Note: donor 2 is heterozygous for sickle cell disease.

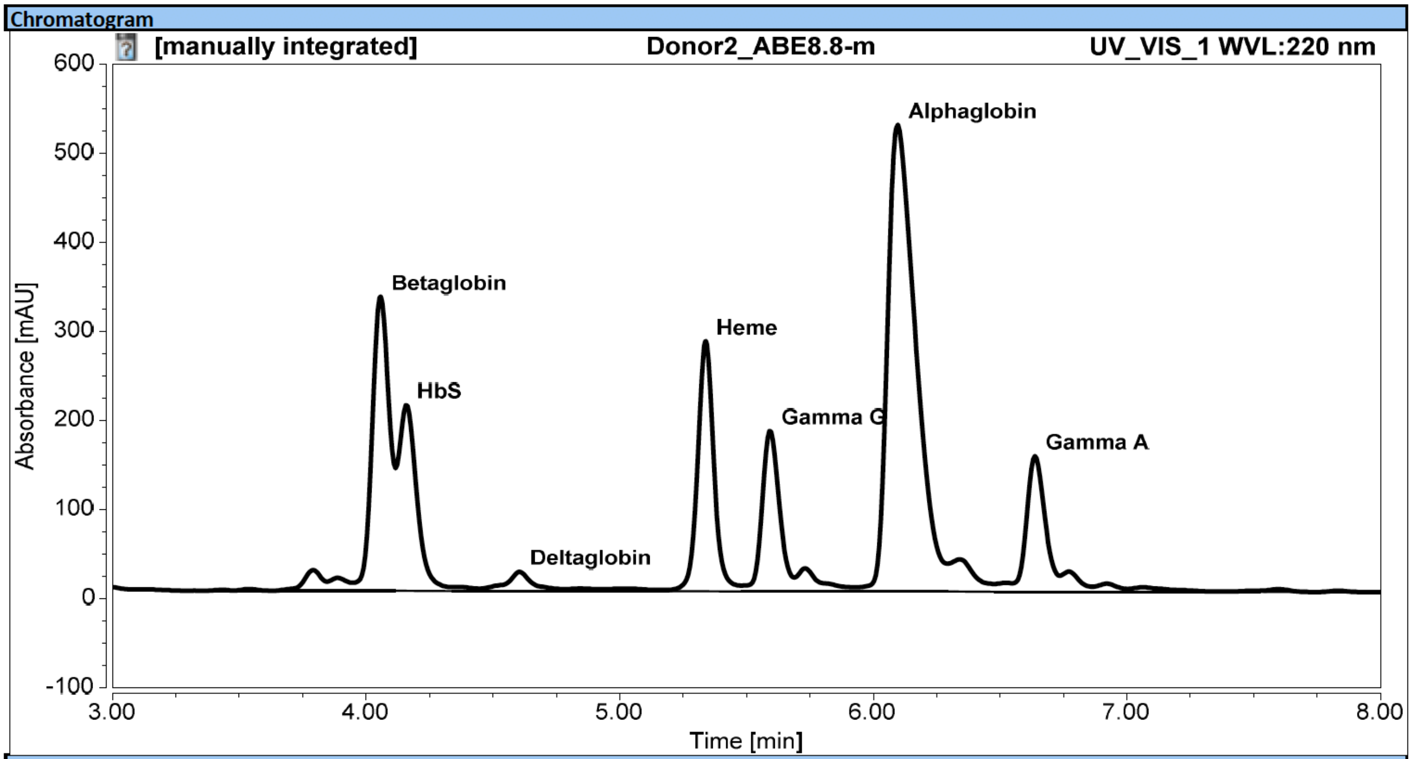


Integration Results

| No. | Peak Name | Retention Time min | Area mAU*min | Height mAU | Relative Area % | Relative Height % |
|---|---|---|---|---|---|---|
| 5 | Betaglobin | 4.057 | 26.492 | 330.353 | 14.76 | 17.48 |
| 6 | HbS | 4.160 | 17.650 | 209.154 | 9.84 | 11.07 |
| 9 | Deltaglobin | 4.603 | 2.190 | 21.674 | 1.22 | 1.15 |
| 14 | Heme | 5.340 | 20.930 | 280.930 | 11.66 | 14.87 |
| 15 | Gamma G | 5.593 | 14.357 | 180.573 | 8.00 | 9.56 |
| 18 | Alphaglobin | 6.097 | 67.616 | 524.013 | 37.68 | 27.73 |
| 21 | Gamma A | 6.637 | 12.935 | 152.286 | 7.21 | 8.06 |

**Supplementary Table 20:** UHPLC UV-Vis trace (220 nm) and integration of globin chain levels of differentiated CD34+ cells treated with ABE8.8-d (donor 2). Note: donor 2 is heterozygous for sickle cell disease.

Chromatogram

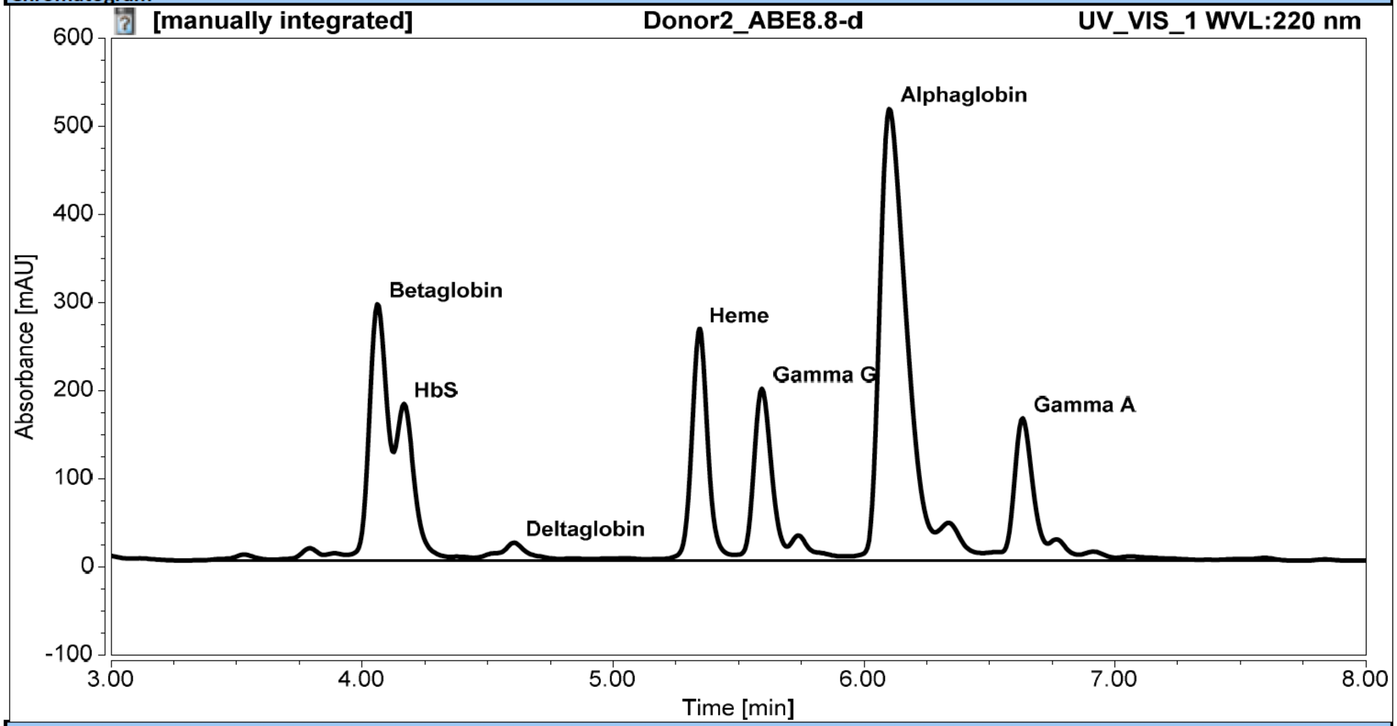


Integration Results

| No. | Peak Name | Retention Time<br>min | Area<br>mAU*min | Height<br>mAU | Relative Area<br>% | Relative Height<br>% |
|---|---|---|---|---|---|---|
| 6 | Betaglobin | 4.063 | 23.888 | 291.371 | 13.90 | 16.06 |
| 7 | HbS | 4.167 | 15.512 | 178.139 | 9.02 | 9.82 |
| 10 | Deltaglobin | 4.607 | 2.249 | 20.296 | 1.31 | 1.12 |
| 15 | Heme | 5.343 | 19.534 | 262.940 | 11.36 | 14.49 |
| 16 | Gamma G | 5.593 | 16.034 | 195.308 | 9.33 | 10.77 |
| 19 | Alphaglobin | 6.103 | 63.055 | 512.839 | 36.68 | 28.27 |
| 22 | Gamma A | 6.633 | 13.860 | 161.449 | 8.06 | 8.90 |

**Supplementary Table 21:** UHPLC UV-Vis trace (220 nm) and integration of globin chain levels of differentiated CD34+ cells treated with ABE8.13-m (donor 2). Note: donor 2 is heterozygous for sickle cell disease.

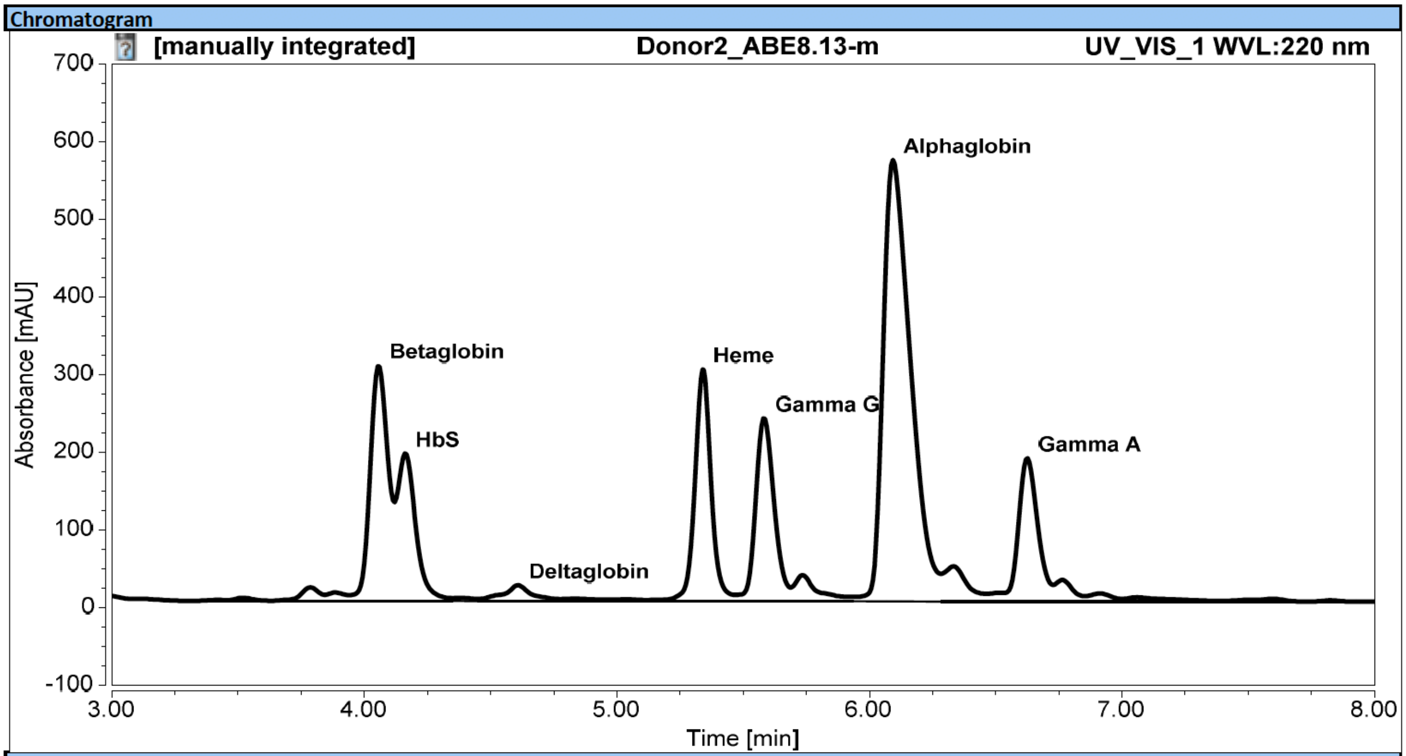


Integration Results

| No. | Peak Name | Retention Time min | Area mAU*min | Height mAU | Relative Area % | Relative Height % |
|---|---|---|---|---|---|---|
| 6 | Betaglobin | 4.057 | 24.649 | 303.680 | 12.77 | 15.05 |
| 7 | HbS | 4.163 | 16.705 | 190.174 | 8.65 | 9.43 |
| 10 | Deltaglobin | 4.607 | 2.114 | 20.478 | 1.10 | 1.01 |
| 15 | Heme | 5.340 | 22.158 | 298.385 | 11.48 | 14.79 |
| 16 | Gamma G | 5.583 | 20.107 | 236.242 | 10.41 | 11.71 |
| 19 | Alphaglobin | 6.093 | 71.587 | 568.640 | 37.08 | 28.18 |
| 22 | Gamma A | 6.623 | 16.259 | 184.774 | 8.42 | 9.16 |

**Supplementary Table 22:** UHPLC UV-Vis trace (220 nm) and integration of globin chain levels of differentiated CD34+ cells treated with ABE8.13-d (donor 2). Note: donor 2 is heterozygous for sickle cell disease.

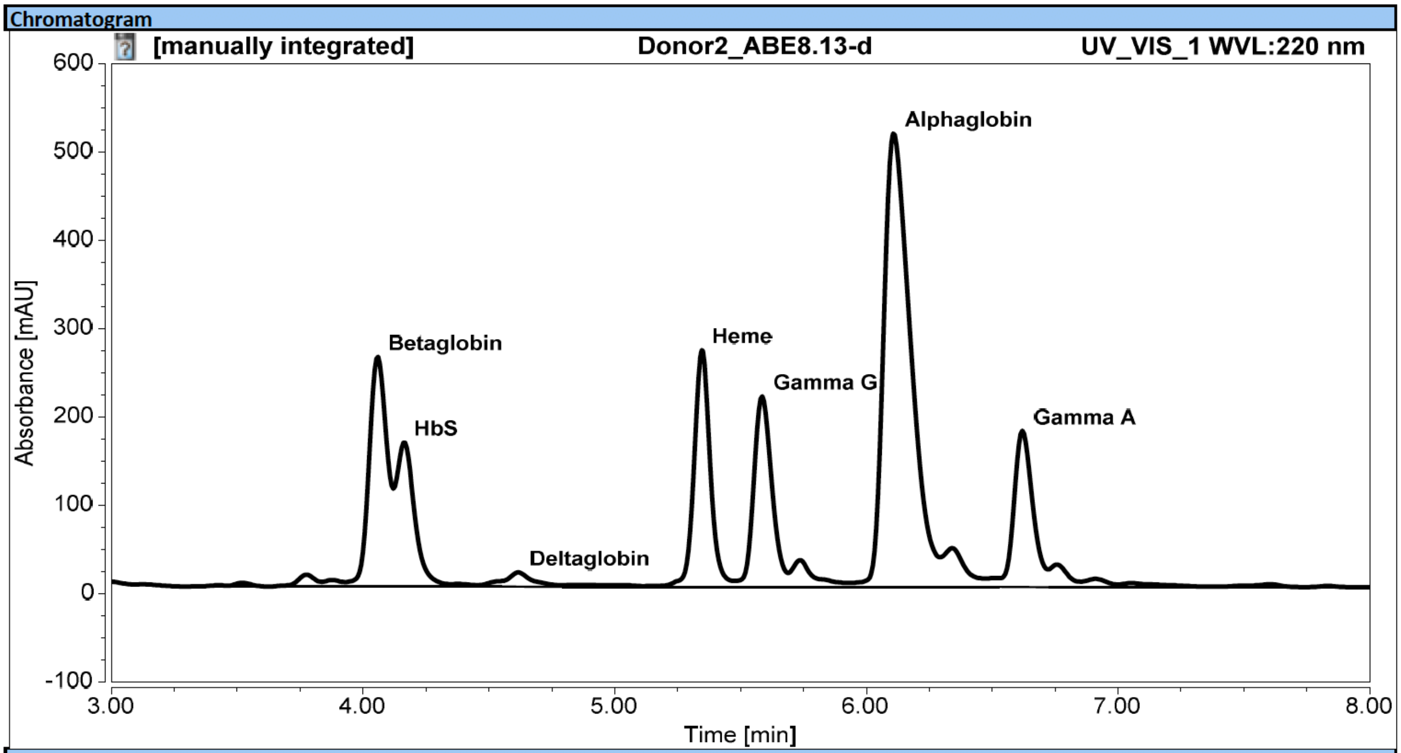


Integration Results

| No. | Peak Name | Retention Time min | Area mAU*min | Height mAU | Relative Area % | Relative Height % |
|---|---|---|---|---|---|---|
| 5 | Betaglobin | 4.057 | 21.912 | 260.274 | 12.79 | 14.49 |
| 6 | HbS | 4.163 | 13.772 | 163.531 | 8.04 | 9.10 |
| 9 | Deltaglobin | 4.617 | 1.827 | 16.318 | 1.07 | 0.91 |
| 13 | Heme | 5.347 | 19.932 | 268.078 | 11.64 | 14.92 |
| 14 | Gamma G | 5.587 | 18.026 | 215.798 | 10.52 | 12.01 |
| 17 | Alphaglobin | 6.110 | 64.064 | 513.833 | 37.40 | 28.61 |
| 20 | Gamma A | 6.620 | 15.335 | 177.074 | 8.95 | 9.86 |

**Supplementary Table 23:** UHPLC UV-Vis trace (220 nm) and integration of globin chain levels of differentiated CD34+ cells treated with ABE8.17-m (donor 2). Note: donor 2 is heterozygous for sickle cell disease.

Chromatogram

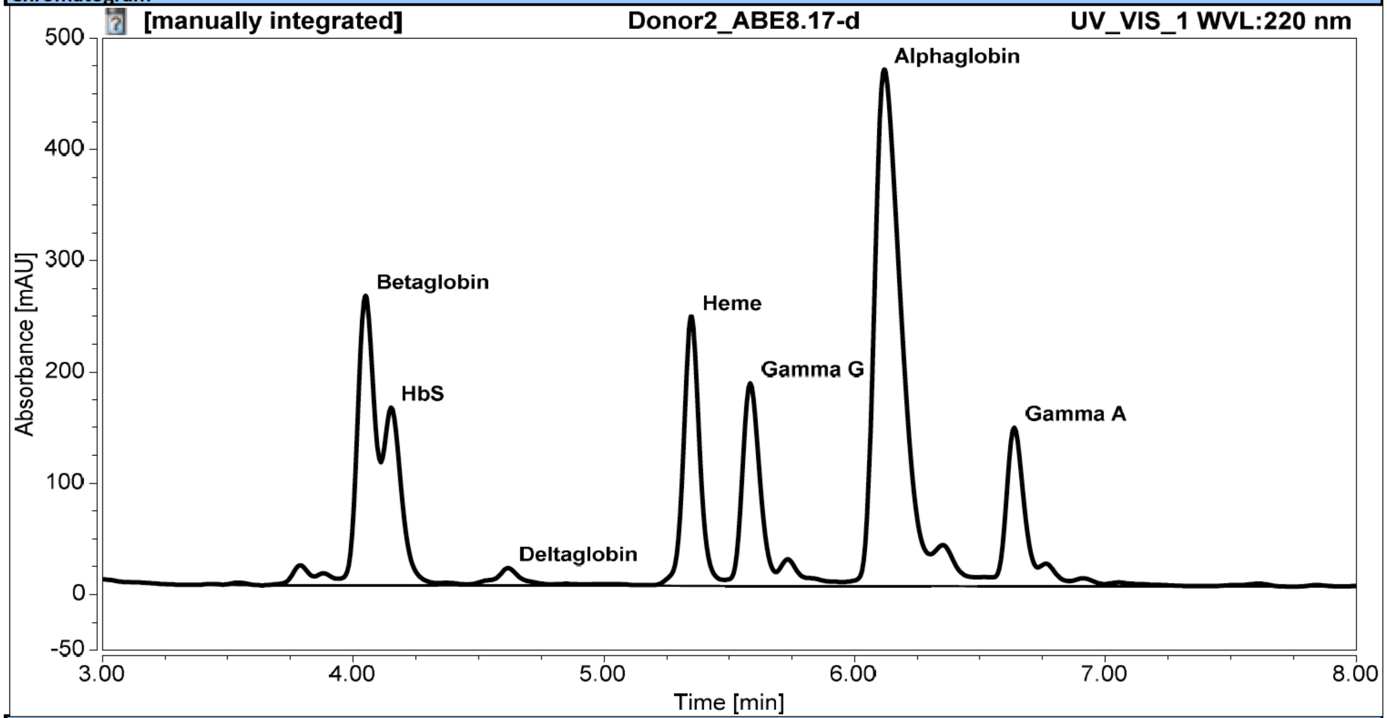


Integration Results

| No. | Peak Name | Retention Time<br>min | Area<br>mAU*min | Height<br>mAU | Relative Area<br>% | Relative Height<br>% |
|---|---|---|---|---|---|---|
| 5 | Betaglobin | 4.050 | 20.688 | 260.391 | 13.65 | 15.97 |
| 6 | HbS | 4.150 | 13.530 | 159.994 | 8.93 | 9.81 |
| 9 | Deltaglobin | 4.617 | 1.590 | 15.794 | 1.05 | 0.97 |
| 14 | Heme | 5.350 | 18.069 | 242.095 | 11.92 | 14.85 |
| 15 | Gamma G | 5.583 | 14.825 | 182.475 | 9.78 | 11.19 |
| 18 | Alphaglobin | 6.117 | 57.055 | 464.210 | 37.65 | 28.48 |
| 21 | Gamma A | 6.637 | 11.691 | 142.851 | 7.72 | 8.76 |

**Supplementary Table 24:** UHPLC UV-Vis trace (220 nm) and integration of globin chain levels of differentiated CD34+ cells treated with ABE8.17-d (donor 2). Note: donor 2 is heterozygous for sickle cell disease.

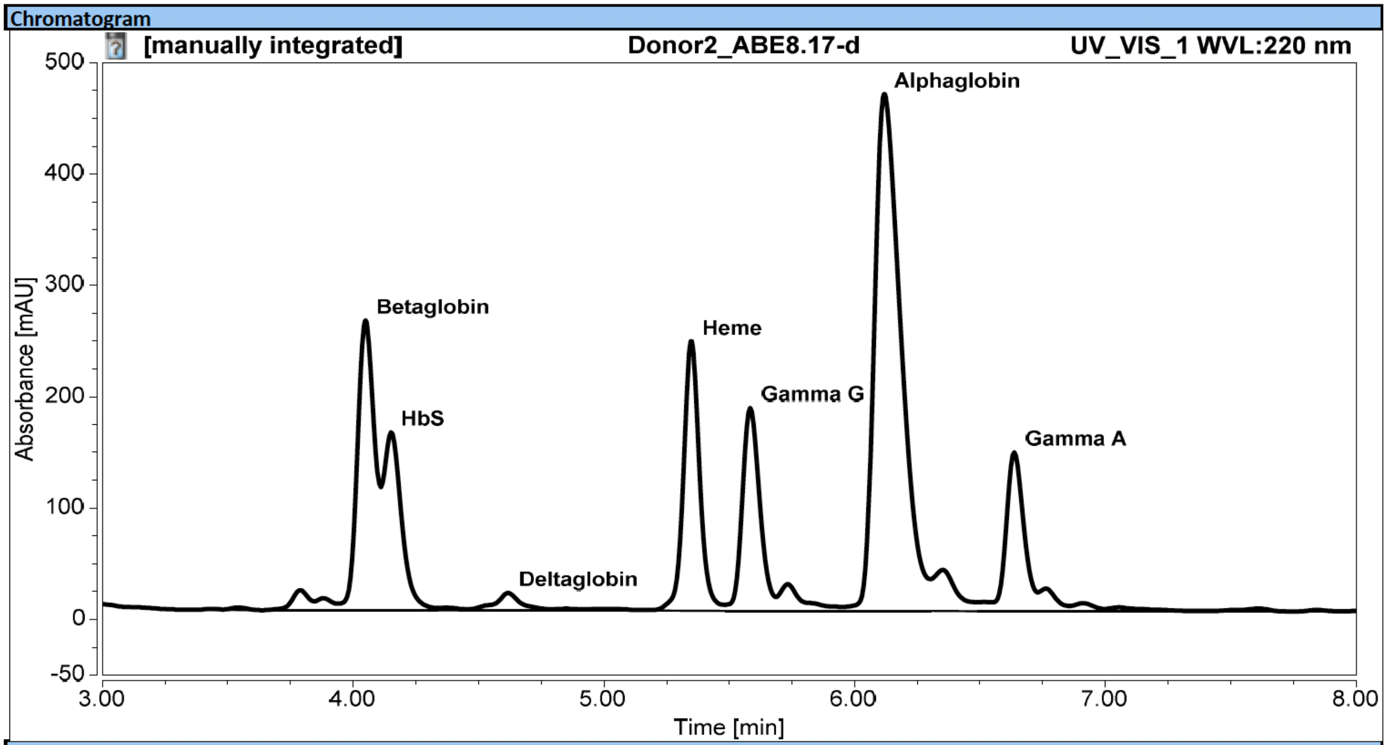


Integration Results

| No. | Peak Name | Retention Time min | Area mAU*min | Height mAU | Relative Area % | Relative Height % |
|---|---|---|---|---|---|---|
| 5 | Betaglobin | 4.050 | 20.688 | 260.391 | 13.65 | 15.97 |
| 6 | HbS | 4.150 | 13.530 | 159.994 | 8.93 | 9.81 |
| 9 | Deltaglobin | 4.617 | 1.590 | 15.794 | 1.05 | 0.97 |
| 14 | Heme | 5.350 | 18.069 | 242.095 | 11.92 | 14.85 |
| 15 | Gamma G | 5.583 | 14.825 | 182.475 | 9.78 | 11.19 |
| 18 | Alphaglobin | 6.117 | 57.055 | 464.210 | 37.65 | 28.48 |
| 21 | Gamma A | 6.637 | 11.691 | 142.851 | 7.72 | 8.76 |

**Supplementary Table 25:** UHPLC UV-Vis trace (220 nm) and integration of globin chain levels of differentiated CD34+ cells treated with ABE8.20-m (donor 2). Note: donor 2 is heterozygous for sickle cell disease.

Chromatogram

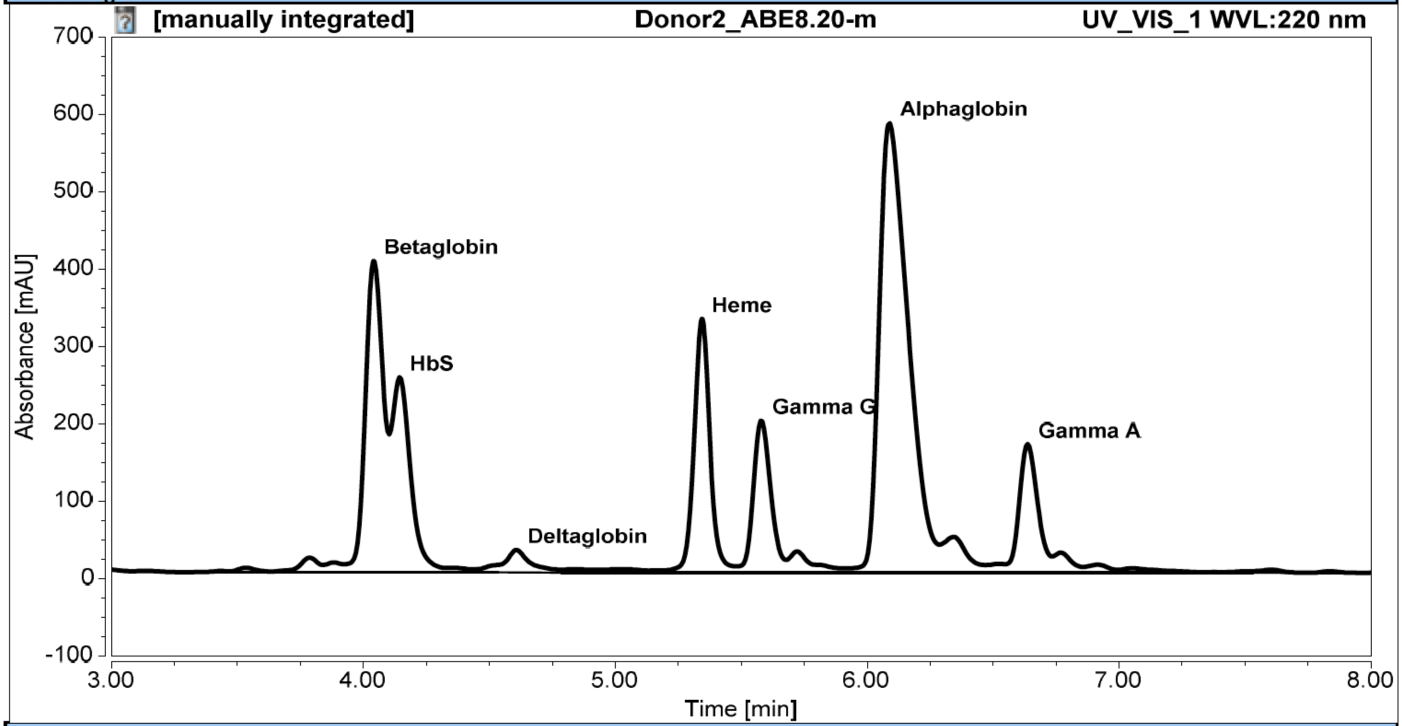


Integration Results

| No. | Peak Name | Retention Time min | Area mAU*min | Height mAU | Relative Area % | Relative Height % |
|---|---|---|---|---|---|---|
| 6 | Betaglobin | 4.040 | 33.339 | 402.775 | 15.75 | 18.52 |
| 7 | HbS | 4.143 | 22.030 | 252.466 | 10.41 | 11.61 |
| 10 | Deltaglobin | 4.607 | 3.256 | 29.080 | 1.54 | 1.34 |
| 14 | Heme | 5.343 | 24.399 | 328.265 | 11.53 | 15.10 |
| 15 | Gamma G | 5.580 | 15.959 | 197.001 | 7.54 | 9.06 |
| 18 | Alphaglobin | 6.087 | 78.149 | 581.487 | 36.93 | 26.74 |
| 21 | Gamma A | 6.637 | 14.315 | 166.369 | 6.76 | 7.65 |

**Supplementary Table 25:** UHPLC UV-Vis trace (220 nm) and integration of globin chain levels of differentiated CD34+ cells treated with ABE8.20-d (donor 2). Note: donor 2 is heterozygous for sickle cell disease.

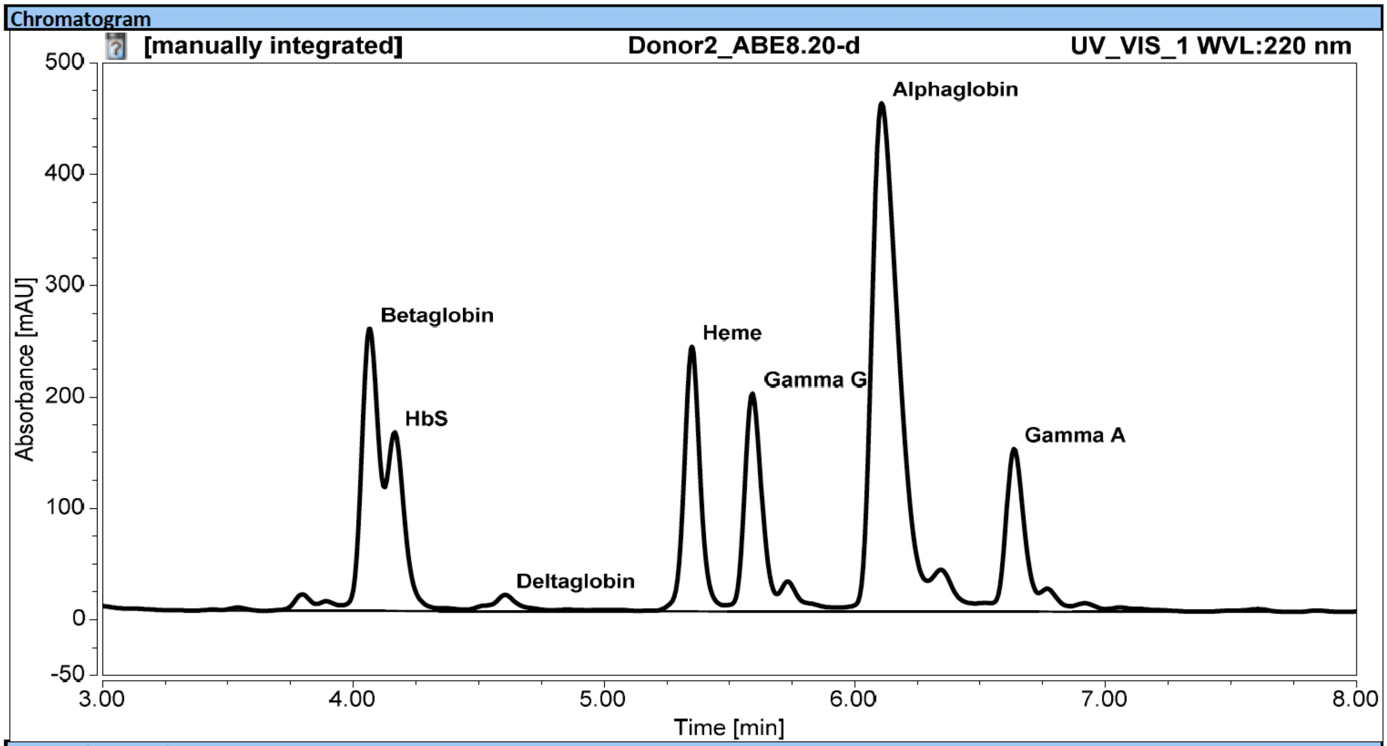


Integration Results

| No. | Peak Name | Retention Time min | Area mAU*min | Height mAU | Relative Area % | Relative Height % |
|---|---|---|---|---|---|---|
| 5 | Betaglobin | 4.067 | 20.307 | 253.509 | 13.46 | 15.57 |
| 6 | HbS | 4.163 | 12.923 | 160.772 | 8.57 | 9.87 |
| 9 | Deltaglobin | 4.607 | 1.534 | 14.720 | 1.02 | 0.90 |
| 14 | Heme | 5.350 | 17.781 | 237.713 | 11.79 | 14.60 |
| 15 | Gamma G | 5.593 | 15.582 | 195.753 | 10.33 | 12.02 |
| 18 | Alphaglobin | 6.107 | 56.188 | 456.808 | 37.25 | 28.05 |
| 21 | Gamma A | 6.633 | 12.248 | 146.288 | 8.12 | 8.98 |

**Supplementary Table 26:** P-values resulting from comparisons between relative cytosine and adenine deamination-induced mutation frequencies amongst treatment groups subjected to whole genome sequencing

| | **ODDS RATIO** | | **%** | |
|---|---|---|---|---|
| Treatment | A->G | C->T | A->G | C->T |
| BE4 | 0.991 | 0.018 | 0.995 | 0.010 |
| ABE7.10-d | 0.883 | 0.883 | 0.911 | 0.944 |
| ABE8.17-m | 0.259 | 0.817 | 0.375 | 0.880 |
| ABE8.20-m | 0.617 | 0.725 | 0.643 | 0.804 |
| Cas9 | 0.275 | 0.383 | 0.270 | 0.643 |
| No Treatment | 0.590 | 0.590 | 0.559 | 0.559 |

**Supplementary Table 27**: Total number of mutations, relative to untreated control, detected in each whole genome sequencing sample

| Sample ID | Treatment | Number of mutations relative to no treatment control |
|---|---|---|
| H17 | ABE7.10-d | 3386 |
| H18 | ABE7.10-d | 3260 |
| H19 | ABE7.10-d | 3237 |
| H21 | ABE7.10-d | 3357 |
| H23 | ABE7.10-d | 3097 |
| H10 | ABE8.17-m | 3558 |
| H12 | ABE8.17-m | 3761 |
| H13 | ABE8.17-m | 3043 |
| H15 | ABE8.17-m | 3623 |
| H16 | ABE8.17-m | 2595 |
| H9 | ABE8.17-m | 3539 |
| H25 | ABE8.20-m | 5101 |
| H26 | ABE8.20-m | 4163 |
| H27 | ABE8.20-m | 2598 |
| H28 | ABE8.20-m | 2996 |
| H29 | ABE8.20-m | 3035 |
| H1 | BE4 | 5947 |
| H3 | BE4 | 6056 |
| H4 | BE4 | 6024 |
| H7 | BE4 | 5394 |
| H8 | BE4 | 5180 |
| H33 | Cas9 | 9532 |
| H34 | Cas9 | 3329 |
| H35 | Cas9 | 3224 |
| H36 | Cas9 | 3872 |
| H37 | Cas9 | 3807 |
| H44 | No Treatment | 3145 |
| H45 | No Treatment | 3659 |
| H46 | No Treatment | 3532 |
| H47 | No Treatment | 3786 |

**Supplementary Note 1**: Discussion of codon optimization and NLS choice for ABE8 design

It has been established that Cas9 codon usage[3,4] and nuclear localization sequence[5] can dramatically alter genome editing efficiencies in eukaryotes. The original Cas9n component of base editors contains six potential polyadenylation sites, leading to poor expression in eukaryotes [2,3,6]. Replacing this with an extensively optimized codon sequence[1] improves base editing efficiencies[7,8]. We assessed the DNA on-target (Supplementary Fig. 4a, 4b), DNA off-target (Supplementary Fig. 4c, 4d) and RNA off-target (Supplementary Fig. 4e) base editing frequencies associated with four ABE constructs, all of which contain a codon-optimized Cas9(D10A)[1]. 1. ABE7.10, which has a single C-terminal BP-SV40 NLS 2. monoABE7.10 which lacks the 5' TadA wild-type portion of ABE7.10, 3. ABEmax, which contains codon-optimized TadA regions and two BPNLS sequences 4. ABEmax(-BPNLS), which has the TadA codon optimization as ABEmax but contains a single C-terminal BP-SV40 NLS.

All four constructs displayed remarkably similar on-target editing efficiencies, indicating that the NLS architecture and TadA codon optimization are not determining for on-target editing efficiency (Supplementary Fig. 4a and 4b). The off-target profiles were also highly similar, but ABEmax displayed significantly greater DNA off-target editing ($p=0.00027$, Students' two-tailed T test) at one site when compared to ABE7.10 (Supplementary Fig. 4c and 4d). ABEmax(-NBPNLS) displayed a 1.6-fold greater mean frequency of RNA off-target editing than ABE7.10 (Supplementary Fig. 4e). Therefore, we used the ABE7.10 architecture for all of our subsequent assays since it demonstrated the most favorable on- and off- target editing profile.

**Supplementary Note 2**: Modeling $\gamma$-globin induction as a function of editor-induced allele frequencies

We sought to explain the relationship between $\gamma$-globin induction (as quantified by UPLC) and the frequency of specific target site edits (or alleles) created by each ABE variant (as quantified by NGS) across two donors. We fitted a linear regression model to the data from the 144-hour timepoint, using the "lm" function in R, with the exact formula being: lm(gamma_over_alpha ~ `7G` + `7G+8G`).

The resulting (best fitting) model ($R^2 = 0.84$) had the following form:
% gamma globin / % alpha globin = 18.50578 + 0.46683 * [Percent 7G] + 0.78554 * [Percent 7G+8G]

The frequency of the "8G" allele was also included in an otherwise identical model and was not found to be statistically significant (P-value = 0.07, two-tailed t-test). Therefore, the model selected for Fig. 3c does not include the "8G" allele.

The finding that the "7G" edit contributes to $\gamma$-globin induction is consistent with what is known from previous human genetics studies, where this allele is known as the British HPFH variant and typically yields 3.5-10% fetal hemoglobin in heterozygous individuals[9]. In addition, the regression model implies that the "7G+8G" edit may have an even larger contribution to $\gamma$-globin induction. Since ABE8 editors generally create higher levels of the "7G+8G" edit, this mechanism is likely to contribute to their overall higher $\gamma$-globin induction.

**Supplementary Note 3:** Discussion of whole genome sequencing data

To generate whole genome sequencing data from ABE7.10-d, ABE8.20-m, ABE8.17-m, Cas9 and BE4-treated cells, we sequenced genomic DNA from five (BE4, Cas9, no treatment, ABE7.10-d and ABE8.20-m) or six (ABE8.17-m) single cell HEK293T cell clones with >20X coverage of the genome. As described in the Methods, the mutational profiles for each sample were calculated relative to one no treatment sample, referred to as the background control, and are displayed in Supplementary Fig. 26 along with the absolute number of relative mutations (Supplementary Table 27).

To determine whether base editor treatment leads to a detectable level of unguided spurious DNA deamination, we used the highly transformed cell line HEK293T as potentially a worst-case scenario, assuming a high degree of chromatin accessibility. As such, we use this system to compare the relative rates of mutations rather than rely on the absolute numbers as a reflection of true mutation rates. In these experiments we calculated whether each sample displayed a statistically significant (Fisher's Exact Test, Figure 4c) increase or decrease in C-to-T mutation relative to the background control (in the case of BE4) or A-to-G (in the case of ABE and Cas9-treated samples). Consistent with prior work[10-13], this analysis confirmed that 5/5 BE4 clones display a significant increase in genome-wide C-to-T mutation (denoted with ** in Supplementary Fig. 27), and that 0/5 ABE7.10-d, 0/4 no treatment, 1/6 ABE8.17-m, 2/5 ABE8.20-m and 1/5 Cas9-treated clones display an increase in A-to-G mutation (denoted with ** in Supplementary Fig. 27). Of note, the Cas9-treated sample that shows an increased number of A-to-G mutations relative to the background control shows a greatly increased number of detected mutations (9,532) when compared to the rest of the sequence samples; the 29 remaining samples fell in the range 3,043-6,056. Intriguingly, 1/5 ABE8.20-m-treated clone displayed a significant reduction (denoted with * in Supplementary Fig. 27) in the frequency of genome-wide A-to-G mutation. Together with the Cas9-treated sample showing a statistically significant increase in A-to-G mutation, these two unexpected data points lead us to believe that there may be variability between cells in our data which could be caused by pre-existing genetic variation within the initial group of HEK293T cells that were transfected with mRNA. For this reason, we preferred to express the data as median rather than mean to mitigate the impact of possible outliers in the dataset.

The protocol we employed (see Methods) was largely based upon prior work from Ye and colleagues[14]. A key difference between our experimental procedures is that we did not begin with HEK293T cells that had grown up from a single cell clone; instead we worked with a sample of

HEK293T cells that were obtained from ATCC and maintained in culture for approximately 4 weeks prior to transfection.

To glean more insight from our data, we performed an analysis of the statistical significance of differences in deamination frequencies of cytosine or adenine residues between treatment groups, using the Mann-Whitney U test (Supplementary Table 26). Our results show that the only group with significant editing of this type (relative to the untreated control group) is the BE4 treatment group, which shows a significant ($p = 0.018$) increase in C•G-to-T•A (or G•C-to-A•T) mutations relative to the untreated control. This assay suggests that the A base editors tested here do not cause spurious deamination in the genome on a comparable level to BE4. However, we recognize that cellular heterogeneity or some other experimental limitation may cause false positive or negative results, and we anticipate that more experimentation in this area will be illuminating.

**Supplementary Note 4:** Targeted NGS analysis details

1. To generate FASTQ files from the base call files (BCF) generated by the MiSeq, demultiplexing was performed by running Illumina bcl2fastq (v2.20.0.422) with the following parameters:

```
bcl2fastq \
        --ignore-missing-bcls \
        --ignore-missing-filter \
        --ignore-missing-positions \
        --ignore-missing-controls \
        --auto-set-to-zero-barcode-mismatches \
        --find-adapters-with-sliding-window \
        --adapter-stringency 0.9 \
        --mask-short-adapter-reads 35 \
        --minimum-trimmed-read-length 35 \
```

2. The FASTQ files created in step (1) were processed using trimmomatic (v0.39)[15] with parameters set up to clip Illumina TruSeq adapters, exclude reads shorter than 20 bases, and trim the remaining 3' end of reads if the average base quality (Phred score) in a 4-bp sliding window dropped below 15. In addition, any bases with quality scores of 3 or lower at the end of reads were removed. Finally, because the round 1 PCR primers include four randomized bases after the read 1 primer sequence, the first four bases of each read were trimmed. The command used to execute trimmomatic is shown below:

```
trimmomatic SE -phred33 $input_fastq $output_fastq \
        ILLUMINACLIP:illumine_adapters.fa:2:30:10 \
        LEADING:3 TRAILING:3 \
        SLIDINGWINDOW:4:15 \
        MINLEN:20 \
        HEADCROP:4
```

3. Reads were aligned to amplicon sequences using bowtie2 (v2.35)[16], in end-to-end mode with the alignment parameters specified by the --very sensitive flag. Reference sequences were determined as the expected amplicon sequences (including primers) for each primer pair based on the human

genome (GRCh38).The SAM files created by bowtie2 were converted to BAM files, sorted, and indexed using the samtools package (v1.9)[17]. Only samples with at least 5,000 aligned reads were considered for analysis.

4. The BAM files created in step (3) were processed using the bam-readcounts tool (https://github.com/genome/bam-readcount) to generate plain text files summarizing the number of non-reference bases, deletions and insertions at each position in the alignment. The minimum base quality (Phred score) for counting a non-reference base was set to 29 in order to exclude low confidence base calls from statistics about editing rates. Only reads with insertions and/or deletions that overlapped the base editor target site (defined as its protospacer + PAM sequence) were counted towards insertion and deletion rates. Editing rates for each position in the target site were calculated as the fraction of non-reference bases of a given type (e.g., G) to the total number of bases passing the base quality threshold at a given position in the alignment.

**Supplementary Note 5:** Whole transcriptome analysis details

1. Lane level FASTQ files were separately aligned to the human genome (Gencode GRCh38v31 primary assembly) using STAR (v2.7.2a) with parameters set to specify the ReadGroup and output both a genome aligned BAM file and a transcriptome aligned BAM file.

2. Lane level genome alignments for each sample created in step (1) were merged, sorted by coordinate, and duplicate marked using Picard (v2.20.5).

3. Reads containing Ns in their cigar string because they span splicing junctions were split using GATK (v4.1.3.0) SplitNCigarReads.

4. Base quality scores were recalibrated using Picard with default settings.

5. Variants were called using GATK HaplotypeCaller. Only reads with a mapping quality ≥ 30 were considered and the minimum base quality (Phred score) for counting a non-reference base was set to 20. Standard settings for variant calling in RNA-seq were used: minimum-base-quality = 20, minimum-mapping-quality = 30, don't-use-soft-clipped-bases, standard-call-conf = 20.

6. Mutations private to base-editor treated samples were identified using background filtration. The highest coverage 'No Treatment' sample was used as the background sample. Only substitutions on canonical chromosomes were considered. Mutations were considered private to the base-editor treated sample if they met the following criteria:

    a. The genomic position of the mutation had coverage ≥ 30 reads in the treated sample and ≥ 20 reads in the untreated sample
    b. The untreated sample had ≥ 99% of reads supporting the reference, non-mutant, base at the position of the mutation
    c. The variant allele frequency of the mutation in the treated sample was ≥ 20%

**Supplementary Note 6:** Whole genome sequencing analysis details

1. Lane level FASTQ files were separately aligned to the human genome (Gencode GRCh38v31 primary assembly) using BWA (0.7.17-r1188) mem with parameters set to specify the ReadGroup. The -M flag was also set to mark shorted split hits at secondary alignments.

2. Lane level genome alignments for each sample created in step (1) were merged, sorted by coordinate, and duplicate marked using Picard (v2.20.5) using default settings.

3. Variants were called using GATK (v4.1.3.0) HaplotypeCaller. Only reads with a mapping quality ≥ 30 were considered and the minimum base quality (Phred score) for counting a non-reference base was set to 20. Standard settings for variant calling in DNA-seq were used.

4. Mutations private to base-editor treated samples were identified using background filtration. The highest coverage 'No Treatment' sample was used as the background sample. Only substitutions on canonical chromosomes were considered. Mutations were considered private to the base-editor treated sample if they met the following criteria:

    a. The genomic position of the mutation had coverage ≥ 10 reads in the treated and untreated sample
    b. The untreated sample had ≥ 99% of reads supporting the reference, non-mutant, base at the position of the mutation

**Supplementary Sequence 1: inactivated kanamycin resistance gene used in ABE8 evolution**

ccggaattgccagctggggcgccctctggtaaggttgggaagccctgcaaagtaaactggatggctttcttgcc
gccaaggatctgatggcgcaggggatcaagatctgatcaagagacaggatgaggatccttcgcATGATCGAAT
AAGATGGATTGCACGCAGGTTCTCCGGCCGCTTAGGTGGAGCGCCTATTCGGCTATGACTGGGCACAACAGACA
ATCGGCTGCTCTGATGCCGCCGTGTTCCGGCTGTCAGCGCAGGGGCGCCCGGTTCTTTTTGTCAAGACCGACCT
GTCCGGTGCCCTGAATGAACTGCAGGACGAGGCAGCGCGGCTATCGTGGCTGGCCACGACGGGCGTTCCTTGCG
CAGCTGTGCTCGACGTTGTCACTGAAGCGGGAAGGGACTGGCTGCTATTGGGCGAAGTGCCGGGGCAGGATCTC
CTGTCATCTCACCTTGCTCCTGCCGAGAAAGTATCCATCATGGCTGATGCAATGCGGCGGCTGCATACGCTTGA
TCCGGCTACCTGCCCATTCGACCACCAAGCGAAACATCGCATCGAGCGAGCACGTACTCGGATGGAAGCCGGTC
TTGTCGATCAGGATGATCTGGACGAAGAGCATCAGGGGCTCGCGCCAGCCGAACTGTTCGCCAGGCTCAAGGCG
CGCATGCCCGACGGCGAGGATCTCGTCGTGACCCATGGCGATGCCTGCTTGCCGAATATCATGGTGGAAAATGG
CCGCTTTTCTGGATTCATTAACTGTGGCCGGCTGGGTGTGGCGGACCGCTATCAGGACATAGCGTTGGCTACCC
GTGATATTGCTGAAGAGCTTGGCGGCGAATGGGCTGACCGCTTCCTCGTGCTTTACGGTATCGCCGCTCCCGAT
TCGCAGCGCATCGCCTTCTATCGCCTTCTTGACGAGTTCTTCTAA

lower case = kanamycin resistance promoter region
red = targeted inactivated portion (Q4* and W15*)
blue = targeted inactive active site of kanamycin resistance gene (D208N)
underline = PAM

#### Supplementary Sequence 2: ABE8.8-m

MSEVEFSHEYWMRHALTLAKRARDEREVPVGAVLVLNNRVIGEGWNRAIGLHDPTAHAEIMALRQGGLVMQNYR
LIDATLYVTFEPCVMCAGAMIHSRIGRVVFGVRNAKTGAAGSLMDVLH**H**PGMNHRVEITEGILADECAALLC**R**F
FRMPR**R**VFNAQKKAQSSTDSGGSSGGSSGSETPGTSESATPESSGGSSGGSDKKYSIGLAIGTNSVGWAVITDE
YKVPSKKFKVLGNTDRHSIKKNLIGALLFDSGETAEATRLKRTARRRYTRRKNRICYLQEIFSNEMAKVDDSFF
HRLEESFLVEEDKKHERHPIFGNIVDEVAYHEKYPTIYHLRKKLVDSTDKADLRLIYLALAHMIKFRGHFLIEG
DLNPDNSDVDKLFIQLVQTYNQLFEENPINASGVDAKAILSARLSKSRRLENLIAQLPGEKKNGLFGNLIALSL
GLTPNFKSNFDLAEDAKLQLSKDTYDDDLDNLLAQIGDQYADLFLAAKNLSDAILLSDILRVNTEITKAPLSAS
MIKRYDEHHQDLTLLKALVRQQLPEKYKEIFFDQSKNGYAGYIDGGASQEEFYKFIKPILEKMDGTEELLVKLN
REDLLRKQRTFDNGSIPHQIHLGELHAILRRQEDFYPFLKDNREKIEKILTFRIPYYVGPLARGNSRFAWMTRK
SEETITPWNFEEVVDKGASAQSFIERMTNFDKNLPNEKVLPKHSLLYEYFTVYNELTKVKYVTEGMRKPAFLSG
EQKKAIVDLLFKTNRKVTVKQLKEDYFKKIECFDSVEISGVEDRFNASLGTYHDLLKIIKDKDFLDNEENEDIL
EDIVLTLTLFEDREMIEERLKTYAHLFDDKVMKQLKRRRYTGWGRLSRKLINGIRDKQSGKTILDFLKSDGFAN
RNFMQLIHDDSLTFKEDIQKAQVSGQGDSLHEHIANLAGSPAIKKGILQTVKVVDELVKVMGRHKPENIVIEMA
RENQTTQKGQKNSRERMKRIEEGIKELGSQILKEHPVENTQLQNEKLYLYYLQNGRDMYVDQELDINRLSDYDV
DHIVPQSFLKDDSIDNKVLTRSDKNRGKSDNVPSEEVVKKMKNYWRQLLNAKLITQRKFDNLTKAERGGLSELD
KAGFIKRQLVETRQITKHVAQILDSRMNTKYDENDKLIREVKVITLKSKLVSDFRKDFQFYKVREINNYHHAHD
AYLNAVVGTALIKKYPKLESEFVYGDYKVYDVRKMIAKSEQEIGKATAKYFFYSNIMNFFKTEITLANGEIRKR
PLIETNGETGEIVWDKGRDFATVRKVLSMPQVNIVKKTEVQTGGFSKESILPKRNSDKLIARKKDWDPKKYGGF
DSPTVAYSVLVVAKVEKGKSKKLKSVKELLGITIMERSSFEKNPIDFLEAKGYKEVKKDLIIKLPKYSLFELEN
GRKRMLASAGELQKGNELALPSKYVNFLYLASHYEKLKGSPEDNEQKQLFVEQHKHYLDEIIEQISEFSKRVIL
ADANLDKVLSAYNKHRDKPIREQAENIIHLFTLTNLGAPAAFKYFDTTIDRKRYTSTKEVLDATLIHQSITGLY
ETRIDLSQLGGDEGADKRTADGSEFESPKKKRKV*

#### Supplementary Sequence 3: ABE8.8-d

MSEVEFSHEYWMRHALTLAKRAWDEREVPVGAVLVHNNRVIGEGWNRPIGRHDPTAHAEIMALRQGGLVMQNYR
LIDATLYVTLEPCVMCAGAMIHSRIGRVVFGARDAKTGAAGSLMDVLHHPGMNHRVEITEGILADECAALLSDF
FRMRRQEIKAQKKAQSSTDSGGSSGGSSGSETPGTSESATPESSGGSSGGSSEVEFSHEYWMRHALTLAKRARD
EREVPVGAVLVLNNRVIGEGWNRAIGLHDPTAHAEIMALRQGGLVMQNYRLIDATLYVTFEPCVMCAGAMIHSR
IGRVVFGVRNAKTGAAGSLMDVLH**H**PGMNHRVEITEGILADECAALLC**R**FFRMPR**R**VFNAQKKAQSSTDSGGSS
GGSSGSETPGTSESATPESSGGSSGGSDKKYSIGLAIGTNSVGWAVITDEYKVPSKKFKVLGNTDRHSIKKNLI
GALLFDSGETAEATRLKRTARRRYTRRKNRICYLQEIFSNEMAKVDDSFFHRLEESFLVEEDKKHERHPIFGNI
VDEVAYHEKYPTIYHLRKKLVDSTDKADLRLIYLALAHMIKFRGHFLIEGDLNPDNSDVDKLFIQLVQTYNQLF
EENPINASGVDAKAILSARLSKSRRLENLIAQLPGEKKNGLFGNLIALSLGLTPNFKSNFDLAEDAKLQLSKDT
YDDDLDNLLAQIGDQYADLFLAAKNLSDAILLSDILRVNTEITKAPLSASMIKRYDEHHQDLTLLKALVRQQLP
EKYKEIFFDQSKNGYAGYIDGGASQEEFYKFIKPILEKMDGTEELLVKLNREDLLRKQRTFDNGSIPHQIHLGE
LHAILRRQEDFYPFLKDNREKIEKILTFRIPYYVGPLARGNSRFAWMTRKSEETITPWNFEEVVDKGASAQSFI
ERMTNFDKNLPNEKVLPKHSLLYEYFTVYNELTKVKYVTEGMRKPAFLSGEQKKAIVDLLFKTNRKVTVKQLKE
DYFKKIECFDSVEISGVEDRFNASLGTYHDLLKIIKDKDFLDNEENEDILEDIVLTLTLFEDREMIEERLKTYA
HLFDDKVMKQLKRRRYTGWGRLSRKLINGIRDKQSGKTILDFLKSDGFANRNFMQLIHDDSLTFKEDIQKAQVS
GQGDSLHEHIANLAGSPAIKKGILQTVKVVDELVKVMGRHKPENIVIEMARENQTTQKGQKNSRERMKRIEEGI
KELGSQILKEHPVENTQLQNEKLYLYYLQNGRDMYVDQELDINRLSDYDVDHIVPQSFLKDDSIDNKVLTRSDK
NRGKSDNVPSEEVVKKMKNYWRQLLNAKLITQRKFDNLTKAERGGLSELDKAGFIKRQLVETRQITKHVAQILD
SRMNTKYDENDKLIREVKVITLKSKLVSDFRKDFQFYKVREINNYHHAHDAYLNAVVGTALIKKYPKLESEFVY
GDYKVYDVRKMIAKSEQEIGKATAKYFFYSNIMNFFKTEITLANGEIRKRPLIETNGETGEIVWDKGRDFATVR
KVLSMPQVNIVKKTEVQTGGFSKESILPKRNSDKLIARKKDWDPKKYGGFDSPTVAYSVLVVAKVEKGKSKKLK
SVKELLGITIMERSSFEKNPIDFLEAKGYKEVKKDLIIKLPKYSLFELENGRKRMLASAGELQKGNELALPSKY
VNFLYLASHYEKLKGSPEDNEQKQLFVEQHKHYLDEIIEQISEFSKRVILADANLDKVLSAYNKHRDKPIREQA
ENIIHLFTLTNLGAPAAFKYFDTTIDRKRYTSTKEVLDATLIHQSITGLYETRIDLSQLGGDEGADKRTADGSE
FESPKKKRKV*

##### Supplementary Sequence 4: ABE8.13-m

MSEVEFSHEYWMRHALTLAKRARDEREVPVGAVLVLNNRVIGEGWNRAIGLHDPTAHAEIMALRQGGLVMQNYR
L**Y**DATLYVTFEPCVMCAGAMIHSRIGRVVFGVRNAKTGAAGSLMDVLH**H**PGMNHRVEITEGILADECAALLC**R**F
FRMPR**R**VFNAQKKAQSSTDSGGSSGGSSGSETPGTSESATPESSGGSSGGSDKKYSIGLAIGTNSVGWAVITDE
YKVPSKKFKVLGNTDRHSIKKNLIGALLFDSGETAEATRLKRTARRRYTRRKNRICYLQEIFSNEMAKVDDSFF
HRLEESFLVEEDKKHERHPIFGNIVDEVAYHEKYPTIYHLRKKLVDSTDKADLRLIYLALAHMIKFRGHFLIEG
DLNPDNSDVDKLFIQLVQTYNQLFEENPINASGVDAKAILSARLSKSRRLENLIAQLPGEKKNGLFGNLIALSL
GLTPNFKSNFDLAEDAKLQLSKDTYDDDLDNLLAQIGDQYADLFLAAKNLSDAILLSDILRVNTEITKAPLSAS
MIKRYDEHHQDLTLLKALVRQQLPEKYKEIFFDQSKNGYAGYIDGGASQEEFYKFIKPILEKMDGTEELLVKLN
REDLLRKQRTFDNGSIPHQIHLGELHAILRRQEDFYPFLKDNREKIEKILTFRIPYYVGPLARGNSRFAWMTRK
SEETITPWNFEEVVDKGASAQSFIERMTNFDKNLPNEKVLPKHSLLYEYFTVYNELTKVKYVTEGMRKPAFLSG
EQKKAIVDLLFKTNRKVTVKQLKEDYFKKIECFDSVEISGVEDRFNASLGTYHDLLKIIKDKDFLDNEENEDIL
EDIVLTLTLFEDREMIEERLKTYAHLFDDKVMKQLKRRRYTGWGRLSRKLINGIRDKQSGKTILDFLKSDGFAN
RNFMQLIHDDSLTFKEDIQKAQVSGQGDSLHEHIANLAGSPAIKKGILQTVKVVDELVKVMGRHKPENIVIEMA
RENQTTQKGQKNSRERMKRIEEGIKELGSQILKEHPVENTQLQNEKLYLYYLQNGRDMYVDQELDINRLSDYDV
DHIVPQSFLKDDSIDNKVLTRSDKNRGKSDNVPSEEVVKKMKNYWRQLLNAKLITQRKFDNLTKAERGGLSELD
KAGFIKRQLVETRQITKHVAQILDSRMNTKYDENDKLIREVKVITLKSKLVSDFRKDFQFYKVREINNYHHAHD
AYLNAVVGTALIKKYPKLESEFVYGDYKVYDVRKMIAKSEQEIGKATAKYFFYSNIMNFFKTEITLANGEIRKR
PLIETNGETGEIVWDKGRDFATVRKVLSMPQVNIVKKTEVQTGGFSKESILPKRNSDKLIARKKDWDPKKYGGF
DSPTVAYSVLVVAKVEKGKSKKLKSVKELLGITIMERSSFEKNPIDFLEAKGYKEVKKDLIIKLPKYSLFELEN
GRKRMLASAGELQKGNELALPSKYVNFLYLASHYEKLKGSPEDNEQKQLFVEQHKHYLDEIIEQISEFSKRVIL
ADANLDKVLSAYNKHRDKPIREQAENIIHLFTLTNLGAPAAFKYFDTTIDRKRYTSTKEVLDATLIHQSITGLY
ETRIDLSQLGGDEGADKRTADGSEFESPKKKRKV*

##### Supplementary Sequence 5: ABE8.13-d

MSEVEFSHEYWMRHALTLAKRAWDEREVPVGAVLVHNNRVIGEGWNRPIGRHDPTAHAEIMALRQGGLVMQNYR
LIDATLYVTLEPCVMCAGAMIHSRIGRVVFGARDAKTGAAGSLMDVLHHPGMNHRVEITEGILADECAALLSDF
FRMRRQEIKAQKKAQSSTDSGGSSGGSSGSETPGTSESATPESSGGSSGGSSEVEFSHEYWMRHALTLAKRARD
EREVPVGAVLVLNNRVIGEGWNRAIGLHDPTAHAEIMALRQGGLVMQNYRL**Y**DATLYVTFEPCVMCAGAMIHSR
IGRVVFGVRNAKTGAAGSLMDVLH**H**PGMNHRVEITEGILADECAALLC**R**FFRMPR**R**VFNAQKKAQSSTDSGGSS
GGSSGSETPGTSESATPESSGGSSGGSDKKYSIGLAIGTNSVGWAVITDEYKVPSKKFKVLGNTDRHSIKKNLI
GALLFDSGETAEATRLKRTARRRYTRRKNRICYLQEIFSNEMAKVDDSFFHRLEESFLVEEDKKHERHPIFGNI
VDEVAYHEKYPTIYHLRKKLVDSTDKADLRLIYLALAHMIKFRGHFLIEGDLNPDNSDVDKLFIQLVQTYNQLF
EENPINASGVDAKAILSARLSKSRRLENLIAQLPGEKKNGLFGNLIALSLGLTPNFKSNFDLAEDAKLQLSKDT
YDDDLDNLLAQIGDQYADLFLAAKNLSDAILLSDILRVNTEITKAPLSASMIKRYDEHHQDLTLLKALVRQQLP
EKYKEIFFDQSKNGYAGYIDGGASQEEFYKFIKPILEKMDGTEELLVKLNREDLLRKQRTFDNGSIPHQIHLGE
LHAILRRQEDFYPFLKDNREKIEKILTFRIPYYVGPLARGNSRFAWMTRKSEETITPWNFEEVVDKGASAQSFI
ERMTNFDKNLPNEKVLPKHSLLYEYFTVYNELTKVKYVTEGMRKPAFLSGEQKKAIVDLLFKTNRKVTVKQLKE
DYFKKIECFDSVEISGVEDRFNASLGTYHDLLKIIKDKDFLDNEENEDILEDIVLTLTLFEDREMIEERLKTYA
HLFDDKVMKQLKRRRYTGWGRLSRKLINGIRDKQSGKTILDFLKSDGFANRNFMQLIHDDSLTFKEDIQKAQVS
GQGDSLHEHIANLAGSPAIKKGILQTVKVVDELVKVMGRHKPENIVIEMARENQTTQKGQKNSRERMKRIEEGI
KELGSQILKEHPVENTQLQNEKLYLYYLQNGRDMYVDQELDINRLSDYDVDHIVPQSFLKDDSIDNKVLTRSDK
NRGKSDNVPSEEVVKKMKNYWRQLLNAKLITQRKFDNLTKAERGGLSELDKAGFIKRQLVETRQITKHVAQILD
SRMNTKYDENDKLIREVKVITLKSKLVSDFRKDFQFYKVREINNYHHAHDAYLNAVVGTALIKKYPKLESEFVY
GDYKVYDVRKMIAKSEQEIGKATAKYFFYSNIMNFFKTEITLANGEIRKRPLIETNGETGEIVWDKGRDFATVR
KVLSMPQVNIVKKTEVQTGGFSKESILPKRNSDKLIARKKDWDPKKYGGFDSPTVAYSVLVVAKVEKGKSKKLK
SVKELLGITIMERSSFEKNPIDFLEAKGYKEVKKDLIIKLPKYSLFELENGRKRMLASAGELQKGNELALPSKY
VNFLYLASHYEKLKGSPEDNEQKQLFVEQHKHYLDEIIEQISEFSKRVILADANLDKVLSAYNKHRDKPIREQA
ENIIHLFTLTNLGAPAAFKYFDTTIDRKRYTSTKEVLDATLIHQSITGLYETRIDLSQLGGDEGADKRTADGSE
FESPKKKRKV*

##### Supplementary Sequence 6: ABE8.17-m

MSEVEFSHEYWMRHALTLAKRARDEREVPVGAVLVLNNRVIGEGWNRAIGLHDPTAHAEIMALRQGGLVMQNYR
LIDATLY**S**TFEPCVMCAGAMIHSRIGRVVFGVRNAKTGAAGSLMDVLHYPGMNHRVEITEGILADECAALLCYF
FRMPR**R**VFNAQKKAQSSTDSGGSSGGSSGSETPGTSESATPESSGGSSGGSDKKYSIGLAIGTNSVGWAVITDE
YKVPSKKFKVLGNTDRHSIKKNLIGALLFDSGETAEATRLKRTARRRYTRRKNRICYLQEIFSNEMAKVDDSFF
HRLEESFLVEEDKKHERHPIFGNIVDEVAYHEKYPTIYHLRKKLVDSTDKADLRLIYLALAHMIKFRGHFLIEG
DLNPDNSDVDKLFIQLVQTYNQLFEENPINASGVDAKAILSARLSKSRRLENLIAQLPGEKKNGLFGNLIALSL
GLTPNFKSNFDLAEDAKLQLSKDTYDDDLDNLLAQIGDQYADLFLAAKNLSDAILLSDILRVNTEITKAPLSAS
MIKRYDEHHQDLTLLKALVRQQLPEKYKEIFFDQSKNGYAGYIDGGASQEEFYKFIKPILEKMDGTEELLVKLN
REDLLRKQRTFDNGSIPHQIHLGELHAILRRQEDFYPFLKDNREKIEKILTFRIPYYVGPLARGNSRFAWMTRK
SEETITPWNFEEVVDKGASAQSFIERMTNFDKNLPNEKVLPKHSLLYEYFTVYNELTKVKYVTEGMRKPAFLSG
EQKKAIVDLLFKTNRKVTVKQLKEDYFKKIECFDSVEISGVEDRFNASLGTYHDLLKIIKDKDFLDNEENEDIL
EDIVLTLTLFEDREMIEERLKTYAHLFDDKVMKQLKRRRYTGWGRLSRKLINGIRDKQSGKTILDFLKSDGFAN
RNFMQLIHDDSLTFKEDIQKAQVSGQGDSLHEHIANLAGSPAIKKGILQTVKVVDELVKVMGRHKPENIVIEMA
RENQTTQKGQKNSRERMKRIEEGIKELGSQILKEHPVENTQLQNEKLYLYYLQNGRDMYVDQELDINRLSDYDV
DHIVPQSFLKDDSIDNKVLTRSDKNRGKSDNVPSEEVVKKMKNYWRQLLNAKLITQRKFDNLTKAERGGLSELD
KAGFIKRQLVETRQITKHVAQILDSRMNTKYDENDKLIREVKVITLKSKLVSDFRKDFQFYKVREINNYHHAHD
AYLNAVVGTALIKKYPKLESEFVYGDYKVYDVRKMIAKSEQEIGKATAKYFFYSNIMNFFKTEITLANGEIRKR
PLIETNGETGEIVWDKGRDFATVRKVLSMPQVNIVKKTEVQTGGFSKESILPKRNSDKLIARKKDWDPKKYGGF
DSPTVAYSVLVVAKVEKGKSKKLKSVKELLGITIMERSSFEKNPIDFLEAKGYKEVKKDLIIKLPKYSLFELEN
GRKRMLASAGELQKGNELALPSKYVNFLYLASHYEKLKGSPEDNEQKQLFVEQHKHYLDEIIEQISEFSKRVIL
ADANLDKVLSAYNKHRDKPIREQAENIIHLFTLTNLGAPAAFKYFDTTIDRKRYTSTKEVLDATLIHQSITGLY
ETRIDLSQLGGDEGADKRTADGSEFESPKKKRKV*

##### Supplementary Sequence 7: ABE8.17-d

MSEVEFSHEYWMRHALTLAKRAWDEREVPVGAVLVHNNRVIGEGWNRPIGRHDPTAHAEIMALRQGGLVMQNYR
LIDATLYVTLEPCVMCAGAMIHSRIGRVVFGARDAKTGAAGSLMDVLHHPGMNHRVEITEGILADECAALLSDF
FRMRRQEIKAQKKAQSSTDSGGSSGGSSGSETPGTSESATPESSGGSSGGSSEVEFSHEYWMRHALTLAKRARD
EREVPVGAVLVLNNRVIGEGWNRAIGLHDPTAHAEIMALRQGGLVMQNYRLIDATLY**S**TFEPCVMCAGAMIHSR
IGRVVFGVRNAKTGAAGSLMDVLHYPGMNHRVEITEGILADECAALLCYFFRMPR**R**VFNAQKKAQSSTDSGGSS
GGSSGSETPGTSESATPESSGGSSGGSDKKYSIGLAIGTNSVGWAVITDEYKVPSKKFKVLGNTDRHSIKKNLI
GALLFDSGETAEATRLKRTARRRYTRRKNRICYLQEIFSNEMAKVDDSFFHRLEESFLVEEDKKHERHPIFGNI
VDEVAYHEKYPTIYHLRKKLVDSTDKADLRLIYLALAHMIKFRGHFLIEGDLNPDNSDVDKLFIQLVQTYNQLF
EENPINASGVDAKAILSARLSKSRRLENLIAQLPGEKKNGLFGNLIALSLGLTPNFKSNFDLAEDAKLQLSKDT
YDDDLDNLLAQIGDQYADLFLAAKNLSDAILLSDILRVNTEITKAPLSASMIKRYDEHHQDLTLLKALVRQQLP
EKYKEIFFDQSKNGYAGYIDGGASQEEFYKFIKPILEKMDGTEELLVKLNREDLLRKQRTFDNGSIPHQIHLGE
LHAILRRQEDFYPFLKDNREKIEKILTFRIPYYVGPLARGNSRFAWMTRKSEETITPWNFEEVVDKGASAQSFI
ERMTNFDKNLPNEKVLPKHSLLYEYFTVYNELTKVKYVTEGMRKPAFLSGEQKKAIVDLLFKTNRKVTVKQLKE
DYFKKIECFDSVEISGVEDRFNASLGTYHDLLKIIKDKDFLDNEENEDILEDIVLTLTLFEDREMIEERLKTYA
HLFDDKVMKQLKRRRYTGWGRLSRKLINGIRDKQSGKTILDFLKSDGFANRNFMQLIHDDSLTFKEDIQKAQVS
GQGDSLHEHIANLAGSPAIKKGILQTVKVVDELVKVMGRHKPENIVIEMARENQTTQKGQKNSRERMKRIEEGI
KELGSQILKEHPVENTQLQNEKLYLYYLQNGRDMYVDQELDINRLSDYDVDHIVPQSFLKDDSIDNKVLTRSDK
NRGKSDNVPSEEVVKKMKNYWRQLLNAKLITQRKFDNLTKAERGGLSELDKAGFIKRQLVETRQITKHVAQILD
SRMNTKYDENDKLIREVKVITLKSKLVSDFRKDFQFYKVREINNYHHAHDAYLNAVVGTALIKKYPKLESEFVY
GDYKVYDVRKMIAKSEQEIGKATAKYFFYSNIMNFFKTEITLANGEIRKRPLIETNGETGEIVWDKGRDFATVR
KVLSMPQVNIVKKTEVQTGGFSKESILPKRNSDKLIARKKDWDPKKYGGFDSPTVAYSVLVVAKVEKGKSKKLK
SVKELLGITIMERSSFEKNPIDFLEAKGYKEVKKDLIIKLPKYSLFELENGRKRMLASAGELQKGNELALPSKY
VNFLYLASHYEKLKGSPEDNEQKQLFVEQHKHYLDEIIEQISEFSKRVILADANLDKVLSAYNKHRDKPIREQA

ENIIHLFTLTNLGAPAAFKYFDTTIDRKRYTSTKEVLDATLIHQSITGLYETRIDLSQLGGDEGADKRTADGSE
FESPKKKRKV*

#### Supplementary Sequence 8: ABE8.20-m

MSEVEFSHEYWMRHALTLAKRARDEREVPVGAVLVLNNRVIGEGWNRAIGLHDPTAHAEIMALRQGGLVMQNYR
LYDATLYSTFEPCVMCAGAMIHSRIGRVVFGVRNAKTGAAGSLMDVLHHPGMNHRVEITEGILADECAALLCRF
FRMPRRVFNAQKKAQSSTDSGGSSGGSSGSETPGTSESATPESSGGSSGGSDKKYSIGLAIGTNSVGWAVITDE
YKVPSKKFKVLGNTDRHSIKKNLIGALLFDSGETAEATRLKRTARRRYTRRKNRICYLQEIFSNEMAKVDDSFF
HRLEESFLVEEDKKHERHPIFGNIVDEVAYHEKYPTIYHLRKKLVDSTDKADLRLIYLALAHMIKFRGHFLIEG
DLNPDNSDVDKLFIQLVQTYNQLFEENPINASGVDAKAILSARLSKSRRLENLIAQLPGEKKNGLFGNLIALSL
GLTPNFKSNFDLAEDAKLQLSKDTYDDDLDNLLAQIGDQYADLFLAAKNLSDAILLSDILRVNTEITKAPLSAS
MIKRYDEHHQDLTLLKALVRQQLPEKYKEIFFDQSKNGYAGYIDGGASQEEFYKFIKPILEKMDGTEELLVKLN
REDLLRKQRTFDNGSIPHQIHLGELHAILRRQEDFYPFLKDNREKIEKILTFRIPYYVGPLARGNSRFAWMTRK
SEETITPWNFEEVVDKGASAQSFIERMTNFDKNLPNEKVLPKHSLLYEYFTVYNELTKVKYVTEGMRKPAFLSG
EQKKAIVDLLFKTNRKVTVKQLKEDYFKKIECFDSVEISGVEDRFNASLGTYHDLLKIIKDKDFLDNEENEDIL
EDIVLTLTLFEDREMIEERLKTYAHLFDDKVMKQLKRRRYTGWGRLSRKLINGIRDKQSGKTILDFLKSDGFAN
RNFMQLIHDDSLTFKEDIQKAQVSGQGDSLHEHIANLAGSPAIKKGILQTVKVVDELVKVMGRHKPENIVIEMA
RENQTTQKGQKNSRERMKRIEEGIKELGSQILKEHPVENTQLQNEKLYLYYLQNGRDMYVDQELDINRLSDYDV
DHIVPQSFLKDDSIDNKVLTRSDKNRGKSDNVPSEEVVKKMKNYWRQLLNAKLITQRKFDNLTKAERGGLSELD
KAGFIKRQLVETRQITKHVAQILDSRMNTKYDENDKLIREVKVITLKSKLVSDFRKDFQFYKVREINNYHHAHD
AYLNAVVGTALIKKYPKLESEFVYGDYKVYDVRKMIAKSEQEIGKATAKYFFYSNIMNFFKTEITLANGEIRKR
PLIETNGETGEIVWDKGRDFATVRKVLSMPQVNIVKKTEVQTGGFSKESILPKRNSDKLIARKKDWDPKKYGGF
DSPTVAYSVLVVAKVEKGKSKKLKSVKELLGITIMERSSFEKNPIDFLEAKGYKEVKKDLIIKLPKYSLFELEN
GRKRMLASAGELQKGNELALPSKYVNFLYLASHYEKLKGSPEDNEQKQLFVEQHKHYLDEIIEQISEFSKRVIL
ADANLDKVLSAYNKHRDKPIREQAENIIHLFTLTNLGAPAAFKYFDTTIDRKRYTSTKEVLDATLIHQSITGLY
ETRIDLSQLGGDEGADKRTADGSEFESPKKKRKV*

#### Supplementary Sequence 9: ABE8.20-d

MSEVEFSHEYWMRHALTLAKRAWDEREVPVGAVLVHNNRVIGEGWNRPIGRHDPTAHAEIMALRQGGLVMQNYR
LIDATLYVTLEPCVMCAGAMIHSRIGRVVFGARDAKTGAAGSLMDVLHHPGMNHRVEITEGILADECAALLSDF
FRMRRQEIKAQKKAQSSTDSGGSSGGSSGSETPGTSESATPESSGGSSGGSSEVEFSHEYWMRHALTLAKRARD
EREVPVGAVLVLNNRVIGEGWNRAIGLHDPTAHAEIMALRQGGLVMQNYRLYDATLYSTFEPCVMCAGAMIHSR
IGRVVFGVRNAKTGAAGSLMDVLHHPGMNHRVEITEGILADECAALLCRFFRMPRRVFNAQKKAQSSTDSGGSS
GGSSGSETPGTSESATPESSGGSSGGSDKKYSIGLAIGTNSVGWAVITDEYKVPSKKFKVLGNTDRHSIKKNLI
GALLFDSGETAEATRLKRTARRRYTRRKNRICYLQEIFSNEMAKVDDSFFHRLEESFLVEEDKKHERHPIFGNI
VDEVAYHEKYPTIYHLRKKLVDSTDKADLRLIYLALAHMIKFRGHFLIEGDLNPDNSDVDKLFIQLVQTYNQLF
EENPINASGVDAKAILSARLSKSRRLENLIAQLPGEKKNGLFGNLIALSLGLTPNFKSNFDLAEDAKLQLSKDT
YDDDLDNLLAQIGDQYADLFLAAKNLSDAILLSDILRVNTEITKAPLSASMIKRYDEHHQDLTLLKALVRQQLP
EKYKEIFFDQSKNGYAGYIDGGASQEEFYKFIKPILEKMDGTEELLVKLNREDLLRKQRTFDNGSIPHQIHLGE
LHAILRRQEDFYPFLKDNREKIEKILTFRIPYYVGPLARGNSRFAWMTRKSEETITPWNFEEVVDKGASAQSFI
ERMTNFDKNLPNEKVLPKHSLLYEYFTVYNELTKVKYVTEGMRKPAFLSGEQKKAIVDLLFKTNRKVTVKQLKE
DYFKKIECFDSVEISGVEDRFNASLGTYHDLLKIIKDKDFLDNEENEDILEDIVLTLTLFEDREMIEERLKTYA
HLFDDKVMKQLKRRRYTGWGRLSRKLINGIRDKQSGKTILDFLKSDGFANRNFMQLIHDDSLTFKEDIQKAQVS
GQGDSLHEHIANLAGSPAIKKGILQTVKVVDELVKVMGRHKPENIVIEMARENQTTQKGQKNSRERMKRIEEGI
KELGSQILKEHPVENTQLQNEKLYLYYLQNGRDMYVDQELDINRLSDYDVDHIVPQSFLKDDSIDNKVLTRSDK
NRGKSDNVPSEEVVKKMKNYWRQLLNAKLITQRKFDNLTKAERGGLSELDKAGFIKRQLVETRQITKHVAQILD
SRMNTKYDENDKLIREVKVITLKSKLVSDFRKDFQFYKVREINNYHHAHDAYLNAVVGTALIKKYPKLESEFVY

GDYKVYDVRKMIAKSEQEIGKATAKYFFYSNIMNFFKTEITLANGEIRKRPLIETNGETGEIVWDKGRDFATVR
KVLSMPQVNIVKKTEVQTGGFSKESILPKRNSDKLIARKKDWDPKKYGGFDSPTVAYSVLVVAKVEKGKSKKLK
SVKELLGITIMERSSFEKNPIDFLEAKGYKEVKKDLIIKLPKYSLFELENGRKRMLASAGELQKGNELALPSKY
VNFLYLASHYEKLKGSPEDNEQKQLFVEQHKHYLDEIIEQISEFSKRVILADANLDKVLSAYNKHRDKPIREQA
ENIIHLFTLTNLGAPAAFKYFDTTIDRKRYTSTKEVLDATLIHQSITGLYETRIDLSQLGGDEGADKRTADGSE
FESPKKKRKV*

Black – wild type TadA
Green – linker
Blue – evolved TadA (ABE8 mutations in red)
Orange – D10A S. pyogenes Cas9
Purple – BP-NLS nuclear localization tag
